# Coordinately regulated interbacterial antagonism defense pathways constitute a bacterial innate immune system

**DOI:** 10.1101/2021.08.22.457286

**Authors:** See-Yeun Ting, Kaitlyn D. LaCourse, Hannah E. Ledvina, Rutan Zhang, Matthew C. Radey, Hemantha D. Kulasekara, Rahul Somavanshi, Savannah K. Bertolli, Larry A. Gallagher, Jennifer Kim, Kelsi M. Penewit, Stephen J. Salipante, Libin Xu, S. Brook Peterson, Joseph D. Mougous

**Affiliations:** Department of Microbiology, University of Washington School of Medicine, Seattle, WA 98109, USA; Department of Medicinal Chemistry, University of Washington School of Pharmacy, Seattle, WA 98195, USA; Department of Laboratory Medicine and Pathology, University of Washington School of Medicine, Seattle, WA 98195; Department of Biochemistry, University of Washington School of Medicine, Seattle, WA 98195, USA; Howard Hughes Medical Institute, University of Washington, Seattle, WA 98195, USA

## Abstract

Bacterial survival is fraught with antagonism, including that deriving from viruses and competing bacterial cells^1–3 4^. It is now appreciated that bacteria mount complex antiviral responses; however, whether a coordinated defense against bacterial threats is undertaken is not well understood. Previously we showed that *Pseudomonas aeruginosa* possess a danger sensing pathway that is a critical fitness determinant during competition against other bacteria^5, 6^. Here, we conducted genome-wide screens in *P. aeruginosa* that reveal three conserved and widespread interbacterial antagonism resistance clusters (*arc1-3*). We find that although *arc1-3* are coordinately activated by the Gac/Rsm danger sensing system, they function independently and provide idiosyncratic defense capabilities, distinguishing them from general stress response pathways. Our findings demonstrate that Arc3 family proteins provide specific protection against phospholipase toxins by preventing the accumulation of lysophospholipids in a manner distinct from previously characterized membrane repair systems. These findings liken the response of *P. aeruginosa* to bacterial threats to that of eukaryotic innate immunity, wherein threat detection leads to the activation of specialized defense systems.

---

Antagonism from other organisms is a threat faced nearly universally by bacteria living in mixed populations, yet we are only beginning to understand the mechanisms employed in defense against these assaults^1, 7, 8^. One defense mechanism that is common in the gut microbiome and potentially other habitats is the production of immunity proteins that grant protection against specific toxins delivered by the type VI secretion system (T6SS)^9^. These “orphan” immunity proteins share homology with and are likely evolved from cognate immunity proteins that protect bacteria from undergoing self-intoxication, which is inherent to the indiscriminate nature of the T6SS delivery mechanism. Core cellular structures have also been implicated in promoting survival during interbacterial antagonism. Extracellular polysaccharide capsules protect *Vibrio cholerae* and *E. coli* from T6SS-based attack of other species^10, 11^, and in *Acinetobacter baumanii*, lysine modification at peptidoglycan crosslinks was suggested to render the cell wall resistant to degradation by amidase toxins^12^. Finally, characterized stress response pathways also appear to contribute to antagonism defense. Indeed, it has been suggested that bacterial stress responses evolved in part as a means of detecting and responding to antagonistic competitors ^13, 14^. A candidate-based approach applied to *E. coli* and *P. aeruginosa* found that stress response genes involved in envelope integrity and acid stress, among others can contribute to resistance against a *V. cholerae* phospholipase toxin^15^. While the *P. aeruginosa* genes identified impacted survival when the toxin was produced heterologously within the organism, they did not influence the fitness of the bacterium during interbacterial competition with *V. cholerae*.

We previously discovered that *P. aeruginosa* mounts an effective, but largely undefined defense upon exposure to an antagonistic competitor, which we named the *<u>P</u>. <u>a</u>eruginosa* <u>r</u>esponse to <u>a</u>ntagonism (PARA) ^5^. The response is activated when a subpopulation of *P. aeruginosa* cells are lysed, releasing intracellular contents that trigger the response in neighboring survivors^6^. PARA is coordinated by the Gac/Rsm global regulatory pathway, which posttranscriptionally controls the expression of nearly 400 genes^16, 17^. This regulon includes the Hcp secretion island I-encoded type VI secretion system (H1-T6SS), the activity of which enables *P. aeruginosa* to kill or disable competing bacteria through the delivery of a cocktail of toxic effector proteins^18^. While an important component of PARA, data suggest that the H1-T6SS is just one feature of the response; *P. aeruginosa* strains lacking functionality of the two-component system required to initiate Gac/Rsm signaling are severely crippled in interbacterial antagonism defense relative to those lacking only the H1-T6SS^6^. The majority the genes under Gac/Rsm control encode proteins of unknown function and we hypothesized that these could represent uncharacterized, novel mechanisms by which *P. aeruginosa* defends against interbacterial antagonism.

Prior work by our laboratory has shown that *B. thailandensis* (*B. thai*) fiercely antagonizes *P. aeruginosa* using its antibacterial T6SS^5^. Thus, to identify its interbacterial defense factors, we subjected a transposon library of *P. aeruginosa* to antagonism by *B. thai* or a derivative lacking antibacterial T6SS activity (ΔT6S) and used high-throughput sequencing to measure gene-level fitness changes. Duplicate screening of this library revealed 34 genes critical for the survival of *P. aeruginosa* specifically while undergoing antagonism by *B. thai* (Fig. 1A, Supplemental Fig. 1A,B, Supplemental Table 1). Strikingly, 25 of these belong to the Gac/Rsm regulon and three additional hits included genes directly involved in the Gac/Rsm signaling system^16, 17^. Indeed, disruption of genes encoding GacS and GacA, the central sensor kinase and response regulator required for activation of the pathway, respectively, elicited the strongest interbacterial defense defect among all genes in the *P. aeruginosa* genome (Fig. 1A,B). On the contrary, inactivating insertions in *retS*, encoding a hybrid sensor kinase that negatively regulates GacS, increased interbacterial competitiveness.

**Fig. 1.**
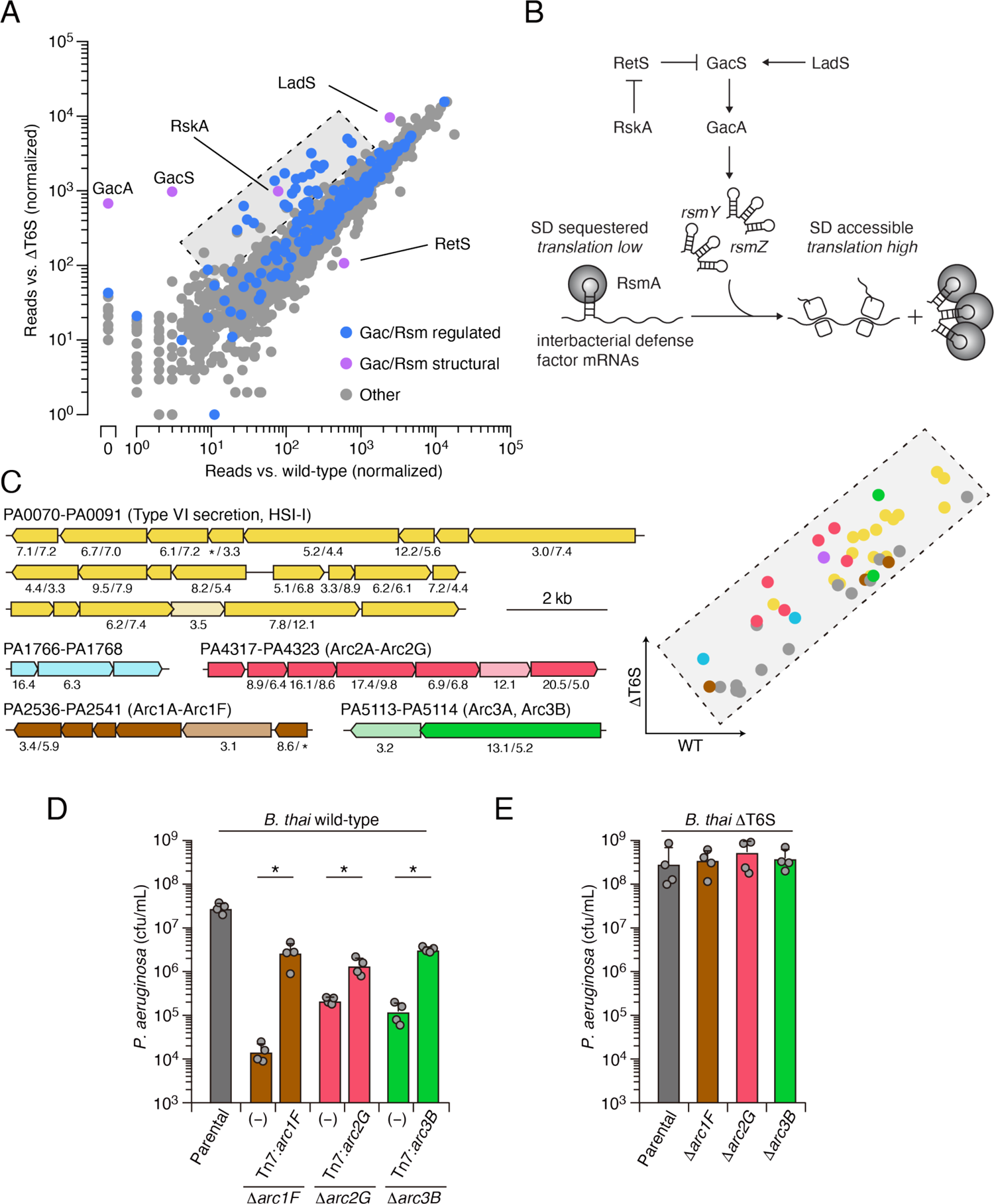
Multiple pathways under Gac/Rsm control contribute to *P. aeruginosa* defense against antagonism. (**A**) Transposon library sequencing-based comparison of the fitness contribution of individual *P. aeruginosa* genes during growth competition with wild-type *B. thai* versus *B. thai* ΔT6S. Genes under Gac/Rsm control (blue) and those encoding core Gac/Rsm regulatory factors (purple, labeled) are indicated. (**B**) Overview of the Gac/Rsm pathway (SD, Shine–Dalgarno). The *rsmY* and *rsmZ* genes encode small RNA molecules that sequester the translational regulator RsmA. (**C**) Left, *P. aeruginosa* gene clusters hit (>three-fold in replicate screens; Supplemental Table 1) in this study. Numbers below genes indicate transposon insertion ratio (*B. thai* ΔT6S/wild-type) for screen in **A** followed by the replicate screen (Supplemental Fig. 1B); light toned genes were hit in only one replicate. Asterisk indicate genes for which insertion were undetected in libraries obtained from *B. thai* wild-type competition. Right, Zoom-in of boxed region of **A** with genes colored corresponding to clusters at left. (**D**,**E**) Recovery of *P. aeruginosa* cells with the indicated genotypes following growth competition against *B. thai* wild-type (**D**) or ΔT6S (**E**).

Our screens identified previously uncharacterized genes belonging to three operons under Gac/Rsm control^16^, several genes within two operons of the HSI-I-encoded T6SS (H1-T6SS)^18^, and two genes within an operon not subject to Gac/Rsm regulation (Fig. 1C, Supplemental Fig. 1C, Supplemental Table 1). Given that false positives are less common among multiple gene hits within an operon, we focused on these genes for subsequent validation in pairwise competition assays. Strains bearing in-frame deletions of the genes most strongly hit in each operon showed substantially reduced fitness in competition with *B. thai*, and in each instance this could be partially restored by genetic complementation (Fig. 1D). Further consistent with our screening results, the strains grew similarly to the wild-type in competition with *B. thai* ΔT6S (Fig. 1E).

The *P. aeruginosa* Gac/Rsm pathway has been implicated in a process we termed danger sensing^5, 6^. We showed that the lysis of neighboring kin cells triggers the pathway, leading to T6SS activation. However, the Gac/Rsm pathway posttransciptionally activates hundreds of genes whose function has largely remained cryptic (Fig. 1B). Bioinformatic analyses suggested that each of the uncharacterized Gac/Rsm-regulated operons hit in our screen likely constitute a discrete pathway. Five of the six genes within the PA2536-41 cluster are predicted to encode proteins related to phospholipid biosynthesis or metabolism, including four genes encoding products with significant homology to enzymes required for phosphatidylglycerol synthesis (Supplemental Fig. 2A). Orthologs of core bacterial phospholipid biosynthetic machinery are encoded elsewhere in the genome, indicating likely specialization of this putative pathway. The PA4317-23 genes co-occur across many species within the Xanthomonadaceae and Pseudomonadaceae (Supplemental Fig. 2B). While none of the proteins encoded by this gene cluster have been characterized, they include a predicted MoxR-like AAA+-ATPase and a protein with a Von Willebrand Factor (VWF) domain. Related protein pairs co-occur widely, and are known to cooperate in a chaperone-like function that releases protein inhibition or stimulates metal cofactor insertion^19, 20^. Finally, PA5113 and PA5114 also co-occur widely, and several organisms encode single polypeptides that represent a fusion of the two proteins (Supplemental Fig. 2C).

**Fig. 2.**
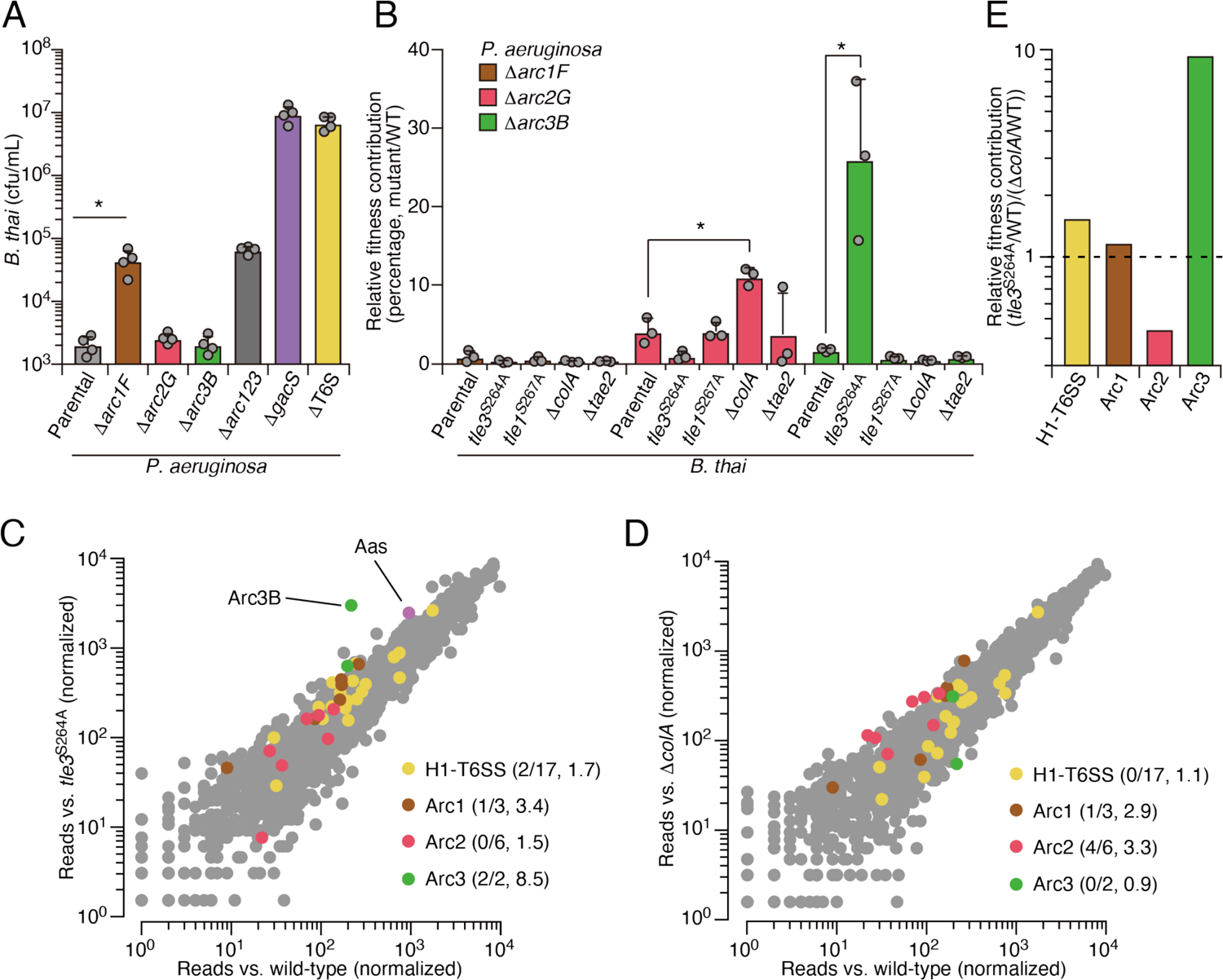
Arc pathways can provide toxin-specific antagonism defense that eclipses that of other cellular factors. (**A**) Recovery of *B. thai* cells following growth competition against a relative abundance of *P. aeruginosa* containing the indicated gene deletions, colored according to Fig. 1C. (**B**) Relative survival of *P. aeruginosa* wild-type versus Arc1-3-inactivated mutants following growth competition against the indicated *B. thai* strains. The loss of ColA or Tle3 activity in *B. thai* increase the relative survival of *P. aeruginosa* lacking Arc2 or Arc3 activity, respectively. **(C,D)** Transposon library sequencing-based comparison of the fitness contribution of individual *P. aeruginosa* genes during growth competition with wild-type *B. thai* versus *B. thai* lacking Tle3 (**C**) or ColA (**D**) activity, colored according to Fig. 1C. Values in parentheses correspond to the number of genes hit (>three-fold change in transposon insertion frequency) within each cluster compared to the number within each cluster hit in the initial screens (*B. thai* wild-type versus *B. thai* ΔT6S) followed by the average transposon insertion frequency ratio (*B. thai* mutant/*B. thai* wild-type) of the genes in the depicted screen. (**E**) Fitness contribution of each *P. aeruginosa* Arc pathway and the H1-T6SS to defense against Tle3 versus ColA.

Next we sought to obtain experimental support for our hypothesis that the Gac/Rsm-regulated operons hit in our screen constitute independent functional units. First, we compared the fitness of our single deletion strains to those bearing in-frame deletions in the two genes most strongly hit within each operon. In each operon, the second deletion did not impact competitiveness with *B. thai*, indicative of an epistatic relationship (Supplemental Fig. 3A). On the contrary, additive effects on interbacterial competitiveness were observed when genes from each of the three operons were inactivated in a single strain. Indeed, the fitness deficiency of this triple mutant strain approaches that of the Δ*gacS* strain, which lacks Gac/Rsm function entirely (Supplemental Fig. 3B). Together, these data reveal that the Gac/Rsm pathway, and discrete functional units under its control, are critical for *P. aeruginosa* survival when antagonized by the T6SS of *B. thai*. Based on these data, we named the genes with the three Gac/Rsm-regulated operons hit in our screen *arc1A-F* (<u>a</u>ntagonism <u>r</u>esistance <u>c</u>luster 1, PA2536-41), *arc2A-G* (PA4317-23) and *arc3A,B* (PA5113-14) (Fig. 1C).

**Fig. 3.**
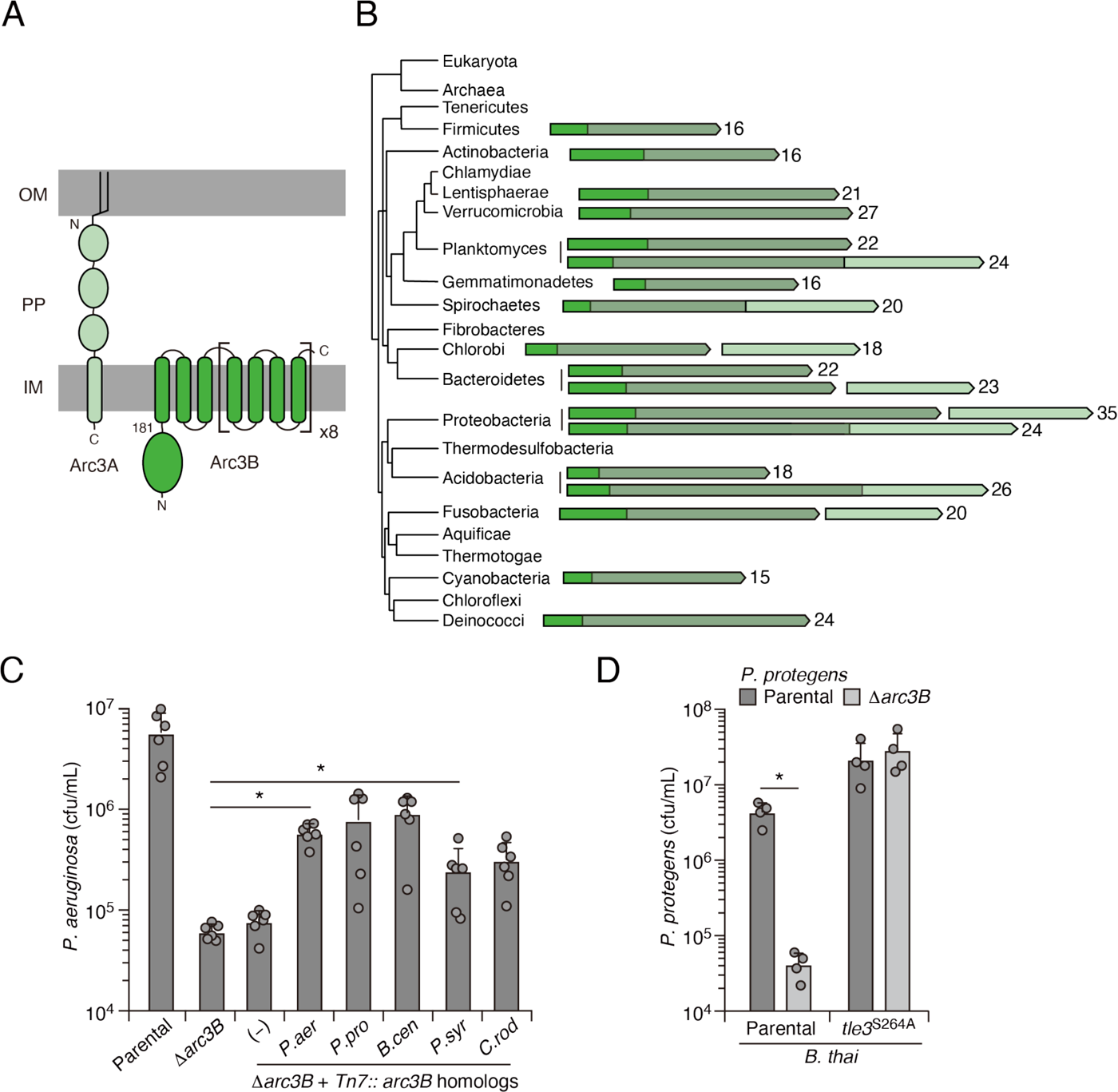
Arc3B is a predicted massive polytopic membrane protein with functionally complementing family members that occur widely in Gram-negative and -positive bacterial phyla. (**A**) Schematic depiction of Arc3A and Arc3B based on bioinformatic predictions. (**B**) Phylogeny of bacterial phyla with representative *arc* genes depicted and colored as in **A**. Membrane associated regions (greyed) and the number of predicted transmembrane segments are indicated. (**C**) Recovery of *P. aeruginosa* strains bearing the indicated mutations and containing a control chromosomal insertion (–) or insertion constructs expressing Arc3B proteins derived from assorted bacteria (*P. aer*, *P. aeruginosa*; *P. pro*, *P. protegens*; *B. cen*, *B. cenocepacia*; *P. syr*, *P. syringae*; *C. rod*, *Citrobacter rodentium*) following growth competition against *B. thai* wild-type. **(D)** Recovery of the indicated *P. protegens* strains following growth competition against *B. thai* wild-type or a strain lacking Tle3 activity.

As a first step toward understanding how Arc1-3 influence the competitiveness of *P. aeruginosa*, we asked whether they serve defensive or offensive roles. In our initial screen, *B. thai* dramatically outnumbered *P. aeruginosa* (∼50:1); therefore, the diminished *P. aeruginosa* viability observed under these conditions suggested each pathway can function defensively.

However, these conditions do not exclusively isolate the defensive contribution of the pathways. For example, we observed that killing competitor cells via the H1-T6SS indirectly contributes to *P. aeruginosa* viability under these conditions, likely by expanding the available growth niche. To probe the capacity of Arc1-3 to antagonize *B. thai*, we initiated co-culture growth competition assays with an excess of *P. aeruginosa.* Under these conditions, Arc2 and Arc3 had no impact on *B. thai* survival, and Arc1 inactivation led to a modest reduction in *B. thai* survival relative to its impact on *P. aeruginosa* defense (Fig. 2A). In contrast, the H1-T6SS reduced *B. thai* survival by approximately four orders of magnitude, eclipsing its impact on *P. aeruginosa* defense by ∼100-fold. Together, these finding indicate that the contributions of Arc1-3 to *P. aeruginosa* competitiveness during antagonism arise predominately from defensive mechanisms.

Our results suggested that the Gac/Rsm system functions analogously to innate immune sensors of eukaryotic organisms, wherein danger sensing activates the expression of independently functioning downstream effectors that defend the cell against specific threats^21^. By extension, this response should be distinct from that provoked by general stressors. To test this, we measured the contribution of Arc1-3 to *P. aeruginosa* survival in conditions known to induce general stress response systems, including heat shock, exposure to H_2_O_2_, and propagation in high salinity media or media containing detergent ^22, 23^. Unlike previously established stress intolerant control strains (Δ*rpoS* and Δ*vacJ*), the survival of strains lacking Arc1-3 function was equivalent to wild-type *P. aeruginosa* in these assays, suggesting that their function is distinct from those involved in general stress resistance (Supplemental Fig. 4). This is also in-line with the regulatory profile of Arc1-3; immunoblotting showed that basal expression each is exceedingly low, yet highly inducible through stimulation of the antagonism responsive Gac/Rsm pathway (Supplemental Fig. 5).

**Fig. 4.**
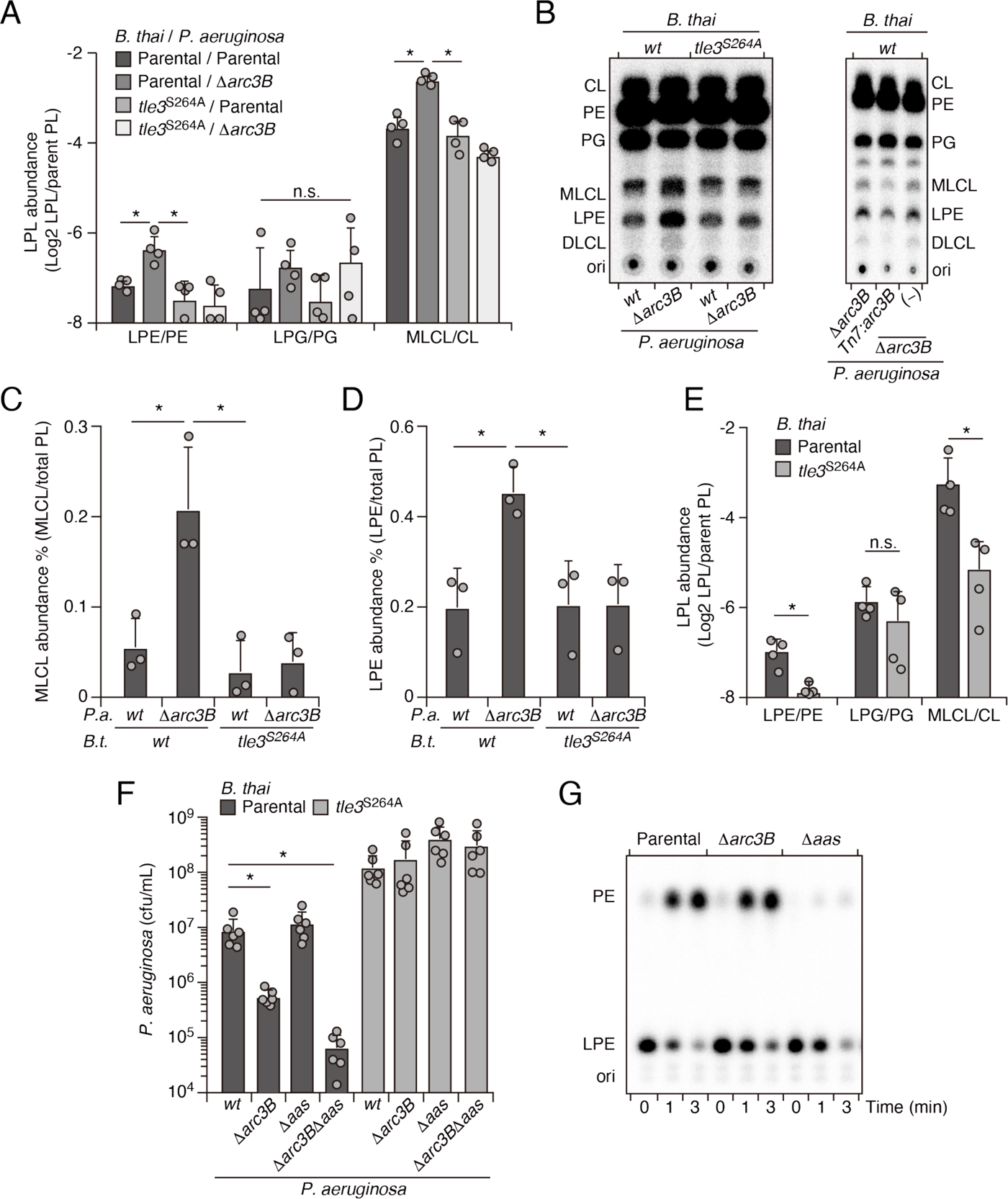
Arc3 prohibits Tle3-catalyzed lysophospholipid accumulation by a mechanism distinct from known pathways. (**A-D**) Mass spectrometric (**A**) and radiographic TLC (**B**) analysis of major phospholipid and lysophospholipid species within lipid extracts derived from the indicated mixtures of *P. aeruginosa* and *B. thai* strains. *P. aeruginosa* were grown in ^32^PO4^2-^ prior to incubation with *B. thai*. Strains were allowed to interact for 1 hr before phospholipids were harvested. Radiolabeled molecules of interest within biological triplicate experiments resolved by TLC were quantified by densitometry (**C**,**D**) (**E**) Mass spectrometric analysis of major phospholipid and lysophospholipid species within lipid extracts derived from mixtures containing *B. thai* lacking Tle3-specific immunity factors competing with the indicated *B. thai* strains. **(F)** Recovery of the indicated *P. aeruginosa* strains following growth competition against *B. thai* wild-type or a strain lacking Tle3 activity. **(G)** Radiographic TLC analysis of products extracted after incubation of *P. aeruginosa*-derived spheroplasts purified radiolabeled LPE.

The specific involvement of Arc1-3 in defense against antagonism led us to hypothesize that the pathways could grant protection against mechanistically distinct threats. Notably, *B. thai* delivers a cocktail of toxins to target cells including two predicted phospholipases (Tle1 and Tle3)^24^, a peptidoglycan-degrading amidase (Tae2) and a colicin-like toxin predicted to form inner membrane pores (ColA)^25, 26^. To determine the role of Arc1-3 in defense against insults caused by toxins with biochemically diverse modes of action, we subjected strains with individual Arc pathways inactivated to pairwise competition against *B. thai* strains lacking single toxins in its arsenal. For Tle1 and Tle3, which are duplicated in the *B. thai* genome and required for basal T6SS function, strains bearing mutated codons corresponding to predicted catalytic residues of the proteins were utilized (Supplemental Fig. 6). Strikingly, we found that Tle3 inactivation abrogated the antagonism defense defect of *P. aeruginosa* lacking Arc3 function (Fig. 2B), whereas its defense defect was maintained in competition experiments against other toxin-inactivated strains of *B. thai*. In contrast, the survival of *P. aeruginosa* lacking Arc2 function was significantly restored by deletion of *colA* in *B. thai*, while none of the single effector mutations within *B. thai* impacted the defense defect of *P. aeruginosa* lacking Arc1 function. These results support the hypothesis that Arc pathways can afford protection in a manner specific to the damage arising from mechanistically distinct threats. *B. thai* delivers toxins beyond those we inactivated in this study, therefore we cannot rule out that Arc1 – like Arc2 and Arc3 – provides effector-specific defense.

We found that Arc2 and Arc3 are major defense factors against insults caused by ColA and Tle3, respectively, suggesting that these toxins may damage cells by a mechanism not efficiently countered by other cellular pathways. To evaluate the relative defensive contribution of Arc2 and Arc3 genome-wide, we performed additional transposon-based screening in which a mutant library of *P. aeruginosa* was grown in competition with *B. thai* strains lacking ColA or Tle3 activity. Remarkably, this experiment defined *arc3B* as the gene most critical for *P. aeruginosa* survival during intoxication by Tle3 (Fig. 2C, Supplemental Table 2). Insertions in the second Arc3 gene, *arc3A*, also crippled *P. aeruginosa* defense against Tle3, though to a lesser degree. Arc2 genes were dispensable for Tle3 defense, and only one gene within Arc1 contributed to Tle3 defense beyond the three-fold insertion frequency ratio cut-off we employed. Inactivation of ColA had a relative minor impact on the fitness of most *P. aeruginosa* transposon mutants (Figure 2D, Supplemental Table 3). However, four of the six Arc2 genes hit in our initial screen provided defense against ColA that exceeded our three-fold cut-off. Additionally, one Arc1 gene met these criteria, whereas Arc3 genes were wholly dispensable for defense against ColA. These results support the model that Arc3 provides specific defense against Tle3, Arc2 provides a degree of specific protection against ColA, and Arc1 serves a broader role in antagonism defense. A comparison of the fitness contributions of each Arc pathway and the H1-T6SS to *P. aeruginosa* undergoing intoxication by Tle3 or ColA highlights the degree of their specificity (Fig. 2E).

The highly specific defense afforded by Arc3 against the predicted phospholipase Tle3 prompted us to further interrogate its function. Arc3B is a large protein containing 35 predicted transmembrane domains and lacking residues strongly indicative of enzymatic activity (Fig. 3A). With predicted outer-membrane anchored N-terminal lipidation and a C-terminal inner-membrane transmembrane helix, Arc3A is expected to span the periplasm. Neither Arc3A nor Arc3B share significant homology with characterized proteins; however, apparent orthologs of each are encoded by neighboring open reading frames across an exceptionally wide distribution of bacteria (Fig. 3B). Arc3B-related proteins are composed of domain of unknown function 2339 (DUF2339), possess large, yet varying numbers of transmembrane segments, and are found in bacteria belonging to most Gram-negative and Gram-positive phyla. Consistent with the predicted N-terminal localization of Arc3A to the outer-membrane, its related proteins are restricted to Gram-negative phyla. Co-immunoprecipitation studies using epitope-tagged variants of the Arc3A and Arc3B encoded at their native chromosomal loci provided evidence that the two proteins stably associate (Supplemental Table 4). The *arc3A* and *arc3B* genes abut a third gene encoded in the same orientation, *estA,* and there is evidence that transcripts containing all three genes are generated by *P. aeruginosa*^27^. Although *estA* encodes an esterase^28^, which could have relevance for phospholipase defense, *estA* was not detected in our co-immunoprecipitation analyses, it did not exhibit differential insertion frequency in our screens, and in-frame deletion of *estA* did not impact *P. aeruginosa* survival during intoxication by Tle3 (Supplemental Tables 1-3, Supplemental Fig. 7A).

The observation that Arc3B proteins are found in bacteria lacking an Arc3A homolog, combined with our finding that Arc3A contributes relatively little to Tle3 defense, led us to speculate that Arc3B plays an intrinsic role in defense, independent of its interaction Arc3A. We tested this by examining the ability of Arc3B-related proteins to complement the defense defect of *P. aeruginosa* Δ*arc3B.* Despite high sequence divergence that would likely preclude specific interactions with Arc3A or other *P. aeruginosa* factors, four of these proteins restored Tle3 defense to a statistically significant degree and two provided protection equivalent to that of *P. aeruginosa arc3B* expressed in the same manner (Fig. 3C). We also examined one of these proteins in its native context. Inactivation of *arc3B* in *P. protegens* conferred a strong interbacterial antagonism defense defect specific to *B. thai* Tle3 (Fig. 3D). In total, these data strongly suggest that proteins in the Arc3B family possess the intrinsic capacity to protect bacteria against interbacterial phospholipase toxins. While we find that Arc3A also participates in defense (Figs. 1A, 2C, Supplemental Table 1, 2), these data suggest that its role is auxiliary.

Enzymes with PLA activity cleave glycerophospholipids at the *sn1* or *sn2* position, generating fatty acid and detergent-like lysophospholipid products^29^. For toxins like Tle3, which are delivered to target cells in exceedingly low quantities^30, 31^, it is likely toxic products, rather than phospholipid depletion *per se*, that most immediately contribute to cell death. Based on the intrinsic capacity of Arc3B to grant protection against Tle3, a predicted PLA, we hypothesized that it directly mitigates damage caused by the toxic products of Tle3. To test this, we measured phospholipid content within extracts derived from competing *P. aeruginosa* and *B. thai* strains at a time point immediately prior to detectable changes in *P. aeruginosa* viability. Strikingly, relative to mixtures containing the wild-type organisms, those containing *P. aeruginosa* Δ*arc3B* possessed highly elevated levels of lysophosphatidylethanolamine (LPE) and monolysocardiolipin (MLCL) in a manner dependent on the activity of Tle3 in *B. thai* (Fig. 4A). Lysophosphatidylglycerol, free fatty acid, and corresponding parent phospholipid levels were not measurably altered by Tle3 (Supplemental Fig. 8A,B).

The accumulated lysophospholipids we observed could derive from *P. aeruginosa* or they could result from retaliatory activity against *B. thai*. To distinguish these possibilities, we subjected *P. aeruginosa* strains with radiolabeled phospholipids to Tle3 intoxication by unlabeled *B. thai* strains. Radiographic TLC analysis of the phospholipids generated in these strain competition mixtures confirmed that *P. aeruginosa* lacking Arc3B accumulates LPE and MLCL during Tle3 intoxication (Fig. 4B-D). Tle3 belongs to a family of effectors for which there remains no definitively characterized member. We were unable to measure the activity of the enzyme in vitro; thus, to determine whether the accumulation of LPE and MLCL were a direct result of Tle3 activity, we quantified phospholipids in extracts derived from *B. thai* strains undergoing Tle3-based self-intoxication (Supplemental Fig. 7B). Samples derived from mixtures composed of a strain with Tle3 and a strain sensitized to Tle3 intoxication through cognate immunity gene inactivation accumulated LPE and MLCL, mirroring our interspecies competition findings (Fig. 4E).

In *E. coli*, lysophospholipids are transported across the inner membrane by the dedicated transporter LplT, after which they are reacylated by Aas^32^. Both LplT and Aas have been shown to play a critical role in defending *E. coli* cells from phospholipase attack^33^. In *P. aeruginosa,* LplT and the acylation domain of Aas are found in a single predicted polypeptide, encoded by PA3267 (*aas*). Interestingly, the *aas* gene was illuminated as a potential *P. aeruginosa* antagonism defense factor in two of the transposon mutant screens we performed and *aas* genes neighbor *arc3* genes in many bacteria (Fig. 2C, Supplemental Fig. 1B, Supplemental Fig. 9, Supplemental Tables 1, 2). In pairwise interbacterial competition assays we were unable to detect the impact of *aas* inactivation on *P. aeruginosa* fitness; however, we found that *P. aeruginosa* strains lacking both *aas* and *arc3B* exhibited a Tle3 defense defect substantially greater than that associated with Arc3B inactivation alone (Fig. 4F). Additionally, we found that Aas is required for PE regeneration from LPE, whereas Arc3B did not contribute to this process (Fig. 4G). These findings indicate that *arc3* encodes a previously undescribed, widespread and independent pathway that prohibits lysophospholipid accumulation by an interbacterial phospholipase toxin.

Evidence suggests that the existence of bacteria is characterized by an onslaught of challenges to their integrity^1, 2^. This is borne out in-part by recent work in the phage arena, which has highlighted a vast number of dedicated bacterial defense systems against these parasites^3, 4^. It is now appreciated that the amalgam of these systems constitutes a bacterial immune system, with both innate and adaptive arms that are functionally analogous, and even evolutionarily related to those of eukaryotic organisms^34^. Our work shows that bacteria also possess a multitude of coordinately regulated pathways dedicated to defense against the threats posed by other bacteria. This further cements the analogy between the immune systems of bacterial and eukaryotic cells, the latter of which also rely on the coordinated production of factors specialized for countering threats of viral or bacterial origin.

We found that Arc3B is the single cellular factor of *P. aeruginosa* most critical for defense against a PLA toxin. Whether Arc3B, perhaps in concert with Arc3A, offer a wider breadth of defensive activity is not known. However, it is notable that phospholipase toxins are ancient and among the most prevalent bacterial toxins known. Even in the absence of direct antagonism by a PLA toxin, environmental lysophospholipids and those generated during normal cellular processes may represent ongoing threats to bacterial viability^35^. We observed low Arc3B expression in the absence of Gac/Rsm activation, arguing that its role in PARA serves an adaptive purpose.

The precise mechanism by which Arc3 proteins defend against intoxication by phospholipase toxins remains unknown. Although we found that Arc3 grants specific protection against Tle3, the relatively low levels of *P. aeruginosa* intoxication achieved by Tle1 does not permit us to rule out that it might also provide broader phospholipase defense. The number and density of transmembrane helices in Arc3B family proteins is unusual and we postulate that these anomalous features of the protein are intimately connected to its mechanism of action. These segments may sequester lysophospholipids and facilitate their turnover or it is conceivable that they directly recruit and interfere with phospholipase enzymes.

We have found that several previously uncharacterized gene clusters under Gac/Rsm control contribute critically to PARA. Our bioinformatic analyses show that the *arc* clusters are widely conserved among Bacteria, yet the Gac/Rsm pathway is restricted to γ-Proteobacteria. If *arc* genes play similar roles in these diverse bacteria, which our data suggest, their expression is also likely subject to induction by antagonism. It will be of interest in future studies to probe the generality of coordinated interbacterial defenses by identifying additional signaling systems responsive to antagonism. Collectively, our findings provide new insight into the diversity of mechanisms bacteria have evolved to defend against interbacterial attack and suggest that many more cryptic specialized defense pathways await discovery.

## Methods

### Bacterial strains and culture conditions

A detailed list of all strains and plasmids used in this study can be found in Supplemental Table 5. Bacterial strains under investigation in this study were derived from *Pseudomonas aeruginosa* PAO1, *Burkholderia thailandensis* E264, and *Pseudomonas protegens* pf-5^36–38^. All strains were grown in Luria-Bertani (LB) broth and incubated either at 37°C (*P. aeruginosa* and *B. thai*) or 30°C (*P. protegens*). For *Pseudomonas* strains the media was supplemented with 25 μg ml^−1^ irgasan, 15 μg ml^−1^ gentamicin, 5% (*w/v*) sucrose, and 0.2% (*w/v*) arabinose as needed.

*B. thai* was cultured with 15 μg ml^−1^ gentamicin and 200 μg ml^−1^ trimethoprim as necessary, and counter-selection for allelic exchange was performed on M9 minimal medium agar plate containing 0.4% (*w/v*) glucose and 0.2% (*w/v*) *p*-chlorophenylalanine. *Escherichia coli* strains used in this study included DH5α for plasmid maintenance, SM10 for conjugal transfer of plasmids into *P. aeruginosa*, *P. protegens*, and *B. thai,* and UE54 for radiolabeled phosphatidylethanolamine (PE) synthesis. *E. coli* strains were grown in LB supplemented with 15 μg ml^−1^ gentamicin, 150 μg ml^−1^ carbenicillin, and 200 μg ml^−1^ trimethoprim as needed.

### Plasmid construction

All primers used in plasmid construction and generation of mutant strains are listed in Supplemental Table 5. In-frame chromosomal deletions and point mutations were created using the suicide vector pEXG2 for *P. aeruginosa* and *P. protegens*, and pJRC115 for *B. thai*^39, 40^. For the generation of chromosomal mutation constructs, 750 bp regions flanking the mutation site were PCR amplified and inserted stitched together into the appropriate vector via Gibson assembly^41^. Site-specific chromosomal insertions in *P. aeruginosa* were generated using pUC18-Tn7t-pBAD-araE as previously described^42^. Our trimethoprim-resistant *B. thai* strain was generated using pUC18T-mini-Tn7-Tp-PS12-mCherry via the mini-Tn7 system^43^.

### Generation of mutant strains

For *P. aeruginosa* and *P. protegens* strain generation: pEXG2 mutation constructs were transformed into *E. coli* SM10, and SM10 donors were subsequently mixed and incubated together with *Pseudomonas* recipients on a nitrocellulose membrane on top of an LB agar plate at a ratio of 10:1 donor–recipient. The cell mixtures were incubated for 6 hours at 37°C to allow for conjugation. These cell mixtures were then scraped up, resuspended into 200 μl LB, and plated on LB agar plates containing irgasan and gentamicin to select for cells containing the mutant construct inserted into the chromosome. *Pseudomonas* merodiploid strains were then grown overnight in non-selective LB media, followed by counter selection on LB no salt agar plates supplemented with sucrose. Gentamicin sensitive, sucrose resistant colonies were screened for allelic replacement by colony PCR and mutations were confirmed via Sanger sequencing of PCR products.

For *B. thai* strain generation, pJRC115 mutation constructs were transformed into *E. coli* SM10. Conjugation to enable plasmid transfer was performed as described above and plated on LB agar containing gentamicin and trimethoprim to select for *B. thai* with the chromosomally integrated plasmid. Merodiploids were grown overnight in LB media at 37°C, then plated on M9 minimal medium containing *p*-chlorophenylalanine for counter selection. Trimethoprim sensitive, p-chlorophenylalanine resistant colonies were screened for allelic replacement by colony PCR and mutations were confirmed via Sanger sequencing of PCR products.

For expression of Arc1F, Arc2G, Arc3B, and Arc3B-homologs in *P. aeruginosa*, pUC18-Tn7t-pBAD-araE containing the gene of interest and the helper plasmid pTNS3 were co-transformed into *P. aeruginosa* strains by electroporation^44^. After 6 hours of outgrowth in LB at 37°C, transformants were plated on a LB agar plate with gentamycin to select for cells with mini-Tn7 integration.

### Bacterial competition assay

For each co-culture competition experiment, donor and recipient strains were first grown for 20 hours in LB medium. Cultures were spun for 1 min at 20,000 g to pellet cells, the supernatant removed, and cells were washed once with fresh LB medium. Cell pellets were resuspended in LB and normalized to OD_600_ = 20. Mixtures of donor and recipient strains were then established at 20:1 (*B. thai* vs. *P. aeruginosa, P. protegens,* or *B. thai*) or 10:1 (*P. aeruginosa* vs. *B. thai*) *v/v* ratios. The initial ratios of donor and recipient strains in these mixtures were measured by performing 10-fold serial dilutions and plating on appropriate selective media to evaluate by colony forming unit (CFU) analysis. The co-culture competitions were initiated by spotting 5 ul of each mixture onto nitrocellulose filters placed on 3% (*w/v*) agar LB no salt plates supplemented with L-arabinose for induction of gene-expression as necessary. Competitions were incubated for 6 hours at 37°C between *B. thai* and *P. aeruginosa*, or 6 hours at 30°C for competition between *B. thai* and *P. protegens.* For *B. thai* self-intoxication, competitions were incubated at 37°C for 3 hours. Cells were harvested by scraping individual spots from excised sections of the nitrocellulose filter into LB medium. Suspensions were serially diluted and plated on selective media for CFU quantification.

### Immunoblotting analysis

To analyze the expression of Arc1F-VSV-G, Arc2G-VSV-G, and Arc3B-VSV-G, *P. aeruginosa* strains were grown in LB broth at 37°C to log phase. Cells were pelleted and resuspended in lysis buffer (20 mM Tris-HCl, pH 7.5, 300 mM NaCl, 10% (*v/v*) glycerol) and then mixed 1:1 with 2X SDS-PAGE sample loading buffer. Samples were then boiled at 100°C for 10 minutes and loaded at equal volumes to resolve using SDS-PAGE, then transferred to nitrocellulose membranes. Membranes were blocked in TBST (10 mM Tris-HCl pH 7.5, 150 mM NaCl, and 0.1% (*w/v*) Tween-20) with 5% (*w/v*) non-fat milk for 30 minutes at room temperature, followed by incubation with anti-VSV-G or with anti-ribosome polymerase β subunit primary antibodies diluted in TBST for 1 hour at room temperature on an orbital shaker. Blots were then washed with TBST, followed by incubation with secondary antibody (Goat anti-Rabbit or anti-Mouse HRP conjugated) diluted in TBST for 30 minutes at room temperature.

Finally, blots were washed with TBST again and developed using Radiance HRP substrate and visualized using iBright imager.

### Lipidomic analysis

To analyze lipid content after intra- and inter-species intoxication, overnight cultures of donor and recipient cells were harvested and mixed at 1 : 1 (*v/v*) ratio in LB medium. These mixtures were spotted onto a 3% (*w/v*) agar LB no salt plate, and incubated for 1 hour at 37°C to allow for intoxication. Cells were then collected, and the lipids were extracted using the Bligh-Dyer method^45^. Briefly, bacterial pellets were resuspended in 0.5 mL of solution containing 0.5 M NaCl in 0.5 N HCl, followed by adding 1.5 mL of chloroform/methanol mixture (1:2, *v/v*).

Suspensions were vortexed at room temperature for 15 minutes, 1.5 mL of 0.5 M NaCl in 0.5 N HCl solution was added to each sample, and vortexed for an additional 5 minutes. Lipid and aqueous layers were separated by centrifugation at 2,000 g for 5 minutes. The lower lipid phase was collected, and these dried lipid samples were analyzed for PE, PG, CL, LPE, LPG, and MLCL content by the Kansas State Lipidomics Research Center.

Lipid samples were analyzed by electrospray ionization triple quadrupole mass spectrometry in direct infusion mode, and data were processed as described by with slight modifications^46^, which include use of a Waters Xevo TQS mass spectrometer (Milford, MA). Internal standards used to normalize data are shown in Supplemental Table 6. For PE and LPE, SPLASH standards were used for normalization (Avanti Polar Lipids, Alabaster, AL). Mass spectrometry global parameters, scan modes, and data processing parameters are shown in Supplemental Table 6.

### Proteomic analysis

*P. aeruginosa* strains containing VSV-G-tagged proteins were grown in 50 mL to mid-log phase, centrifuged at 2,500 g for 15 minutes, and the pellets harvested by resuspension in lysis buffer (20 mM Tris-HCl pH 7.5, 300 mM NaCl, 5% (*w/v*) glycerol, 0.5% (*v/v*) Triton X-100). Cell lysates were prepared by sonication, and tagged proteins were enriched by incubating cell lysates with 30 μl of anti-VSV-G agarose beads at 4°C for 4 hours with constant rotation. Agarose beads were then pelleted by centrifugation at 100 g for 2 minutes, and washed 3 times with 20 mM ammonium bicarbonate. VSV-G agarose beads and bound proteins were then treated with 10 μl of 10 ng/μl (100 ng total per sample) sequencing grade trypsin for 16 hours at 37°C. After digestion, 40 μl of 20 mM ammonium bicarbonate was added to the agarose beads and peptide mixture and lightly mixed. Beads were centrifuged at 300 g for 3 minutes, and the supernatant collected as the peptide fraction. This peptide mixture was reduced with 5 mM Tris(2-carboxyethyl) phosphine hydrochloride for 1 hour, followed by alkylation using 14 mM iodoacetamide for 30 minutes in the dark at room temperature. Alkylation reactions were quenched using 5 mM 1,4-Dithiothreitol. Samples were then diluted with 100% acetonitrile (ACN) and 10% (*w/v*) trifluoroacetic acid (TFA) to a final concentration of 5% ACN (*v/v*) and 0.5% TFA (*w/v*) and applied to MacroSpin C18 columns (30 μg capacity) that had been charged with 100% acetonitrile and ddH_2_O. Bound peptides were then washed twice in 5% (*v/v*) ACN and 0.5% (*w/v*) TFA, before elution with 70% (*v/v*) ACN and 0.1% (*v/v*) formic acid (FA).

Peptides were analyzed by LC-MS/MS using a Dionex UltiMate 3000 Rapid Separation nanoLC and a Q Exactive™ HF Hybrid Quadrupole-Orbitrap™ Mass Spectrometer (Thermo Fisher Scientific Inc, San Jose, CA). Approximately 1 μg of peptide samples was loaded onto the trap column, which was 150 μm × 3 cm in-house packed with 3 um C18 beads. The analytical column was a 75 μm × 10.5 cm PicoChip column packed with 3um C18 beads (New Objective, Inc. Woburn, MA). The flow rate was kept at 300 nL/minute. Solvent A was 0.1% FA in water and Solvent B was 0.1% FA in ACN. The peptide was separated on a 120-minute analytical gradient from 5% ACN/0.1% FA to 40% ACN/0.1% FA. The mass spectrometer was operated in data-dependent mode. The source voltage was 2.10 kV and the capillary temperature was 320 degrees C. MS 1 scans were acquired from 300–2000 m/z at 60,000 resolving power and automatic gain control (AGC) set to 3×10^6^. The top 15 most abundant precursor ions in each MS 1 scan were selected for fragmentation. Precursors were selected with an isolation width of 2 Da and fragmented by Higher-energy collisional dissociation (HCD) at 30% normalized collision energy in the HCD cell. Previously selected ions were dynamically excluded from re-selection for 20 seconds. The MS 2 AGC was set to 1×10^5^.

Proteins were identified from the tandem mass spectra extracted by Xcalibur version 4.0. MS/MS spectra were searched against the Uniprot *P. aeruginosa* PAO1 strain database using Mascot search engine (Matrix Science, London, UK; version 2.5.1). The MS 1 precursor mass tolerance was set to 10 ppm and the MS 2 tolerance was set to 0.05 Da. A 1% false discovery rate cutoff was applied at the peptide level. Only proteins with a minimum of two unique peptides above the cutoff were considered for further study. The search result was visualized by Scaffold (version 4.8.3. Proteome Software, INC., Portland, OR).

### Transposon mutant library construction

A *P. aeruginosa* PAO1 transposon mutant library of ∼80,000 unique Himar1 insertions was prepared using established protocols^47^. Briefly, *P. aeruginosa* were mutagenized by delivery of the transposon- and transposase-bearing suicide vector pBT20 from *E. coli* SM10 λpir^48^.

Insertion mutants were selected on LB agar containing irgasan and gentamycin and pooled. The library was harvested and transposon insertion sites of the complete pool were defined by transposon insertion sequencing as described below.

### Transposon mutant library screen in bacterial competition

The *P. aeruginosa* transposon mutant library was grown at 37°C to mid-log phase. Cell suspensions were then normalized to OD_600_ = 1 and incubated with 100 OD_600_ of *B. thai* donor strains (wild type, Δ*icmF*, *tle3*^S264A^, or Δ*colA*). Competitions were performed on 3% (*w/v*) agar LB no salt plates at 37°C for 7 hours. Cells were washed once with PBS, diluted, and plated on LB agar plates supplemented with irgsan to select for viable *P. aeruginosa* cells. After overnight incubation, cells were harvested, and genomic DNA was directly extracted from pellets using Qiagen Blood and Tissue Midi gDNA prep kit. Transposon insertion sequencing libraries were generated from 3 µg gDNA per sample using the C-tailing method as described^49^ with the transposon-specific primers listed in Supplemental Table 5. For PCR round 1, PCR_1A and PCR_1B were used. For PCR round 2, PCR_2A, PCR_2B, and PCR_2C were used. For sequencing, Seq_primer was used. Libraries were pooled and sequenced in multiplex using an Illumina MiSeq with a 5% PhiX spike-in.

### Tn-Seq data analysis

Seqmagick was used to trim the first six bases from each read (https://github.com/fhcrc/seqmagick). Reads were then aligned, and counts were enumerated using TRANSIT TPP (https://transit.readthedocs.io/en/latest/tpp.html). An annotation GFF was created from the original *P. aeruginosa* PAO1 GFF file (GCF_000006765.1) and was translated to TRANSIT portable format using the TRANSIT ‘convert gff_to_prot_table’ command (https://transit.readthedocs.io/en/latest/transit_running.html#prot-tables-annotations). Finally, the TRANSIT ‘export combined_wig’ command was used to combine and annotate the counts.

Supplementary Table S7 provides summary data of the sequencing runs and their processing by TRANSIT.

### Bioinformatic analysis of Arc3 genes distribution

The phylogenetic profiler tool in IMG^50^ was used to determine the distribution of Arc3B homologs (identified as proteins classified under the Pfam 10101) across bacterial phyla. The number and location of predicted transmembrane domains present in representative Arc3B homologs from each phylum was calculated using CCTOP and TMpred (, https://embnet.vital-it.ch/software/TMPRED_form.html)^51^. Homologs of Arc3A encoded adjacent to Arc3B-like genes were identified on the basis of their classification in the Pfam 13163.

### Stress resistance assay

*P. aeruginosa* cultures were assayed for the ability to survive oxidative stress (50 mM H_2_O_2_), heat stress (55°C), osmotic stress (3 M NaCl), and detergent stress (0.5% (*w/v*) SDS and 0.25 mM EDTA) in LB media as previously described with modifications^22^. After incubation with the oxidative, heat and osmotic stress conditions, cultures were serially diluted 10-fold in LB medium and plated on LB agar for subsequent analysis of remaining viable cells by CFU determination. For oxidative stress, stationary-phase cultures were diluted to OD_600_ = 0.1 and incubated in LB media containing 50 mM H_2_O_2_ for 30 minutes at 37°C with continuous shaking. For heat stress, stationary-phase cultures were diluted in pre-warmed LB media to OD_600_ = 0.1 and incubated at 55°C for 30 minutes with continuous shaking. For osmotic stress, overnight cultures were diluted in LB media containing 3 M NaCl to OD_600_ = 0.1. Cultures were incubated at 37°C for 20 hours with shaking. For detergent stress, cultures were serially diluted in LB media and directly plated on LB agar plates containing 0.5% (*w/v*) SDS and 0.25 mM EDTA for CFU quantification.

### Thin Layer Chromatography - Phospholipid analysis

To prepare [^32^P]-labeled bacteria, *P. aeruginosa* strains were grown in LB media supplemented with 5 μCi/ml [^32^P]-orthophosphoric acid. Tle3 intoxication was performed by mixing 200 OD_600_ of *B. thai* and [^32^P]-*P. aeruginosa* at a 10 : 1 ratio (*v/v*) and incubating this mixture at 37°C for 1 hour on 3% (*w/v*) agar LB no salt plates containing L-arabinose as necessary. After 1 hour of intoxication, cells were collected, and the lipids were extracted using the Bligh-Dyer method as described above. Purified lipid samples were loaded onto a silica gel thin-layer plate and developed with chloroform/methanol/acetic acid/ddH_2_O (85:15:10:3.5, *v/v/v/v*) solvent system^52^. The air-dried plate was exposed to a storage phosphor screen.

Individual phospholipid was visualized and quantified using phosphorimaging to calculate phospholipid content expressed as mol % of the total phospholipid pool.

### PE regeneration assay

To generate [^32^P]-labeled PE, *E. coli* UE54 strain was grown in LB media supplemented with 5 μCi/ml [^32^P]-orthophosphate overnight at 37°C^53^. Cells were collected and the lipids were extracted using the Bligh-Dyer method as described above. The extracted lipids were dried and resuspended in buffer containing 100 mM HEPES-NaOH, pH=7.5, 100 mM KCl, 10 mM CaCl_2_, and 1% (*w/v*) n-Dodecyl-β-D-Maltoside. 10 units of pancreas Phospholipase A2 were added to digest PE to LPE at 37°C overnight with shaking. After incubation, lipids were extracted and loaded onto a silica gel thin-layer plate and developed with chloroform/methanol/acetic acid/ddH_2_O (85:15:10:3.5, *v/v/v/v*) solvent system. The dried plate was exposed to an x-ray film for 2 hours. The phospholipid bands were visualized by developing the film, and bands corresponding [^32^P]-LPE on the TLC plate were scraped, extracted, and resuspended in 100% ethanol.

*P. aeruginosa* spheroplasts were generated by resuspending log-phase culture in 25 mM Tris-HCl, pH 8, 450 mM sucrose, and 1.4 mM EDTA. After addition of 20 μg ml^-1^ lysozyme, cells were incubated on ice for 30 minutes. Intact spheroplasts were collected by centrifugation (3,000 g for 5 minutes) at 4°C and gently resuspended in 25 mM Tris-HCl, pH 8, and 450 mM sucrose. To examine PE regeneration, [^32^P]-LPE was added into spheroplast solutions and incubated at 37°C. The reactions were terminated at the indicated time by adding chloroform-methanol mixture (1:2, *v/v*). The lipids were extracted, separated by TLC, and analyzed using phosphorimaging as described above^54^.

## Data availability

Sequence data associated with this study is available from the Sequence Read Archive at BioProject PRJNA754428 (http://www.ncbi.nlm.nih.gov/bioproject/754428).

## Acknowledgements

We thank Simon Dove, Josh Woodward, Lei Zheng and Mougous laboratory members for helpful discussions, Mikhail Bogdanov and Lei Zheng for sharing reagents, and Colin Manoil and Jason Smith for sharing equipment. This work was supported by the grants from NIH (AI080609 to J.D.M., DK089507 to S.J.S., and R01AI136979 to L.X.) and the Cystic Fibrosis Foundation (SINGH19R0). Equipment utilized was supported by the Office of the Director, National Institutes of Health under award number S10OD026741. JDM is an HHMI Investigator.

## Author contributions

S.-Y.T., K.D.L., H.E.L., S.B.P. and J.D.M. designed the study, S.-Y.T., K.D.L., H.E.L., R.Z., H.D.K., R.S., S.K.B., L.A.G., J.K., K.M.P., and S.J.S. performed experiments, S.-Y.T., K.D.L., H.E.L., M.C.R., L.X., S.B.P., and J.D.M analyzed data, and S.-Y.T., S.B.P., and J.D.M. wrote the manuscript with input from the other authors.

## Author Information

Correspondence and requests for materials should be addressed to J.D.M.

---

**Supplemental Fig. 1.**
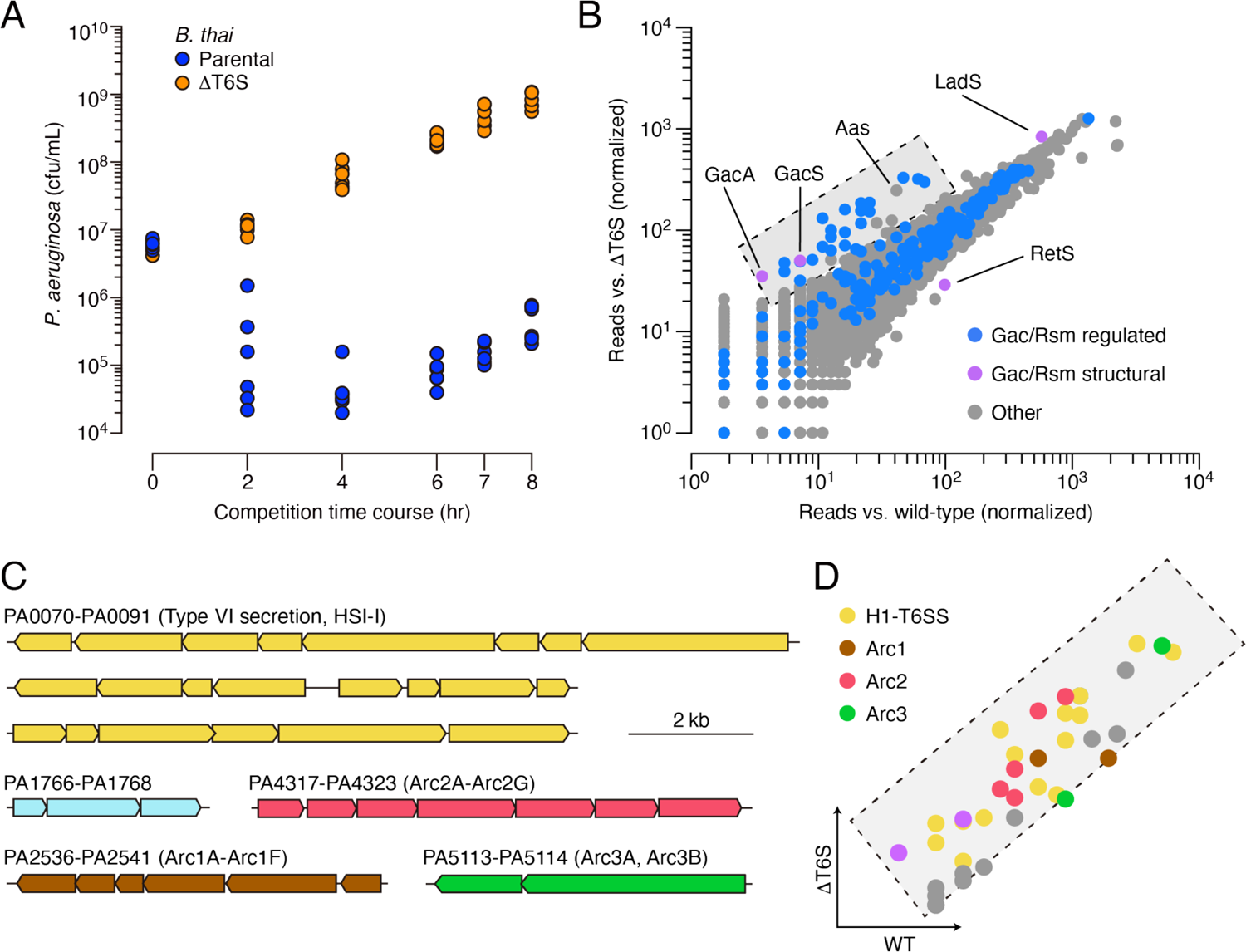
Multiple pathways under Gac/Rsm control contribute to *P. aeruginosa* defense against antagonism. **(A)** Growth of wild-type *P. aeruginosa* in co-culture with the indicated *B. thai* strains. **(B)** Transposon library sequencing-based comparison of the fitness contribution of individual *P. aeruginosa* genes during growth competition with wild-type *B. thai* versus *B. thai* ΔT6S. Genes under Gac/Rsm control (blue) and those encoding core Gac/Rsm regulatory factors (purple, labeled) are indicated. This experiment is a biological replicate of that in Fig. 1A. **(C)** *P. aeruginosa* gene clusters hit (Supplemental Table 1) in this study. **(D)** Zoom-in of boxed region of **B** with genes colored corresponding to clusters at left.

**Supplemental Fig. 2.**
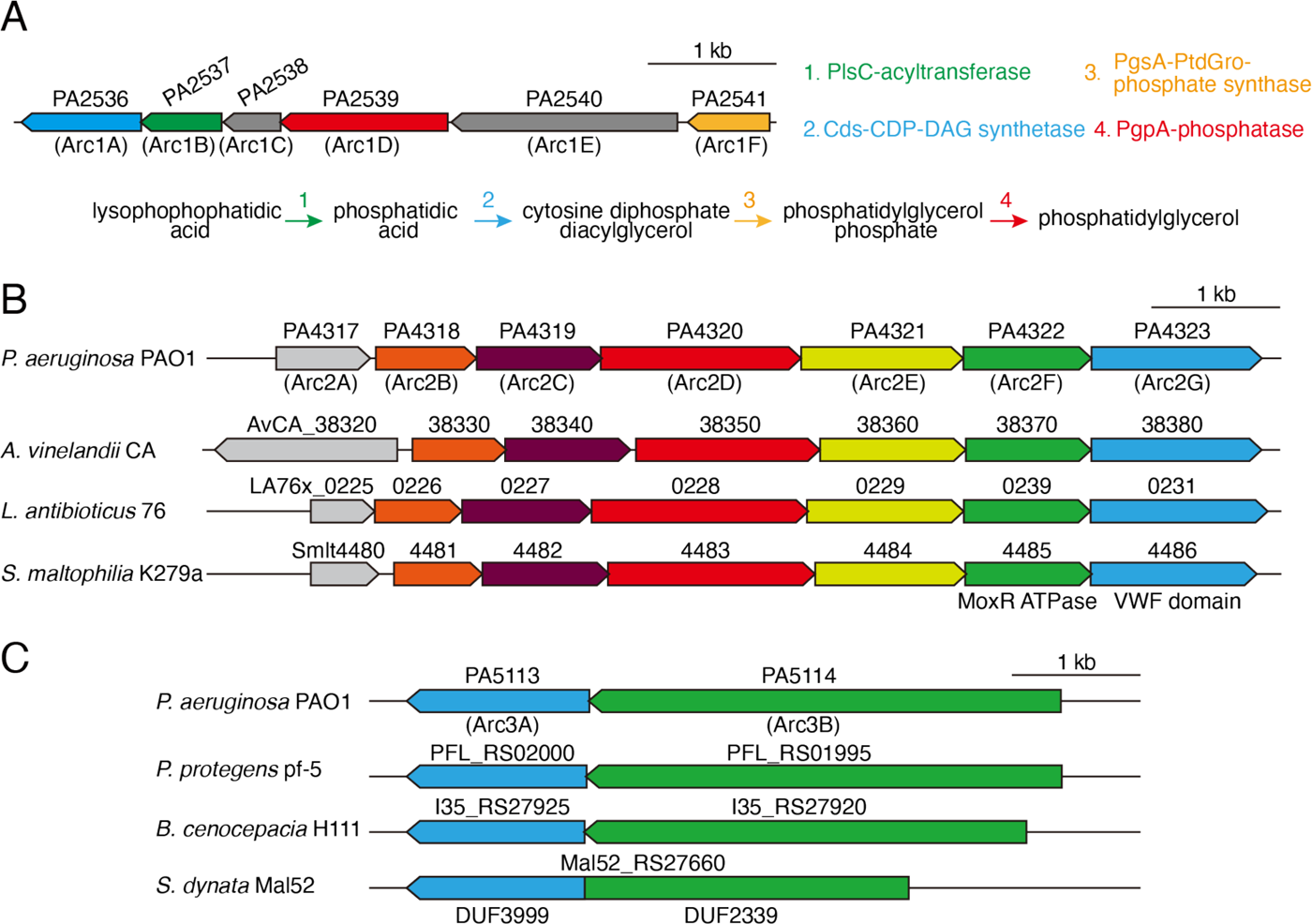
Antagonism resistance clusters possess functionally related genes and are in diverse bacteria. **(A)** Schematic depicting the *arc1* gene cluster of *P. aeruginosa*. Predicted enzymatic activity of Arc1A, B, D, F are indicated (right) and placed onto a phospholipid biosynthesis pathway (bottom). **(B)** Depiction of *arc2* genes from the indicated species. Genes encoding the predicted MoxR-like ATPase and Von Willebrand Factor (VWF) domain protein are indicated below the respective genes. Orthologs are colored to highlight synteny. **(C)** Depiction of *arc2* genes from the indicated species. *S. dynata* exemplifies an instance of translational fusion of *arc3A,B*. Domains of unknown function constituting the two proteins are indicated below.

**Supplemental Fig. 3.**
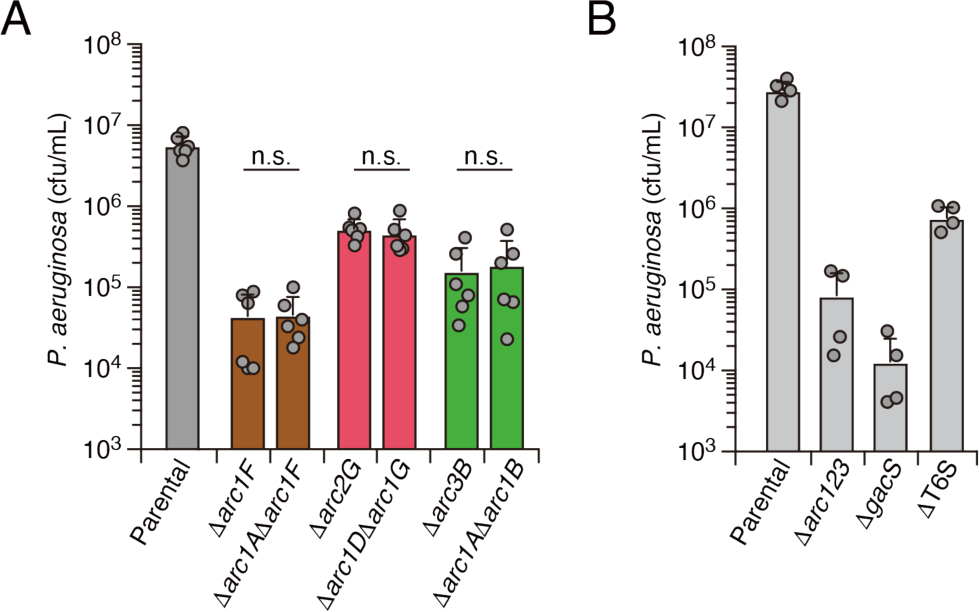
Interbacterial competition assays measure the contribution of *arc1-3* to *P. aeruginosa* fitness during antagonism by *B. thai*. (A) Recovery of *P. aeruginosa* strains bearing the indicated single or double deletions within *arc1-3* following growth competition against wild-type *B. thai*. **(B)** Recovery of *P. aeruginosa* cells with the indicated genotypes following growth competition against wild-type *B. thai*.

**Supplemental Fig. 4.**
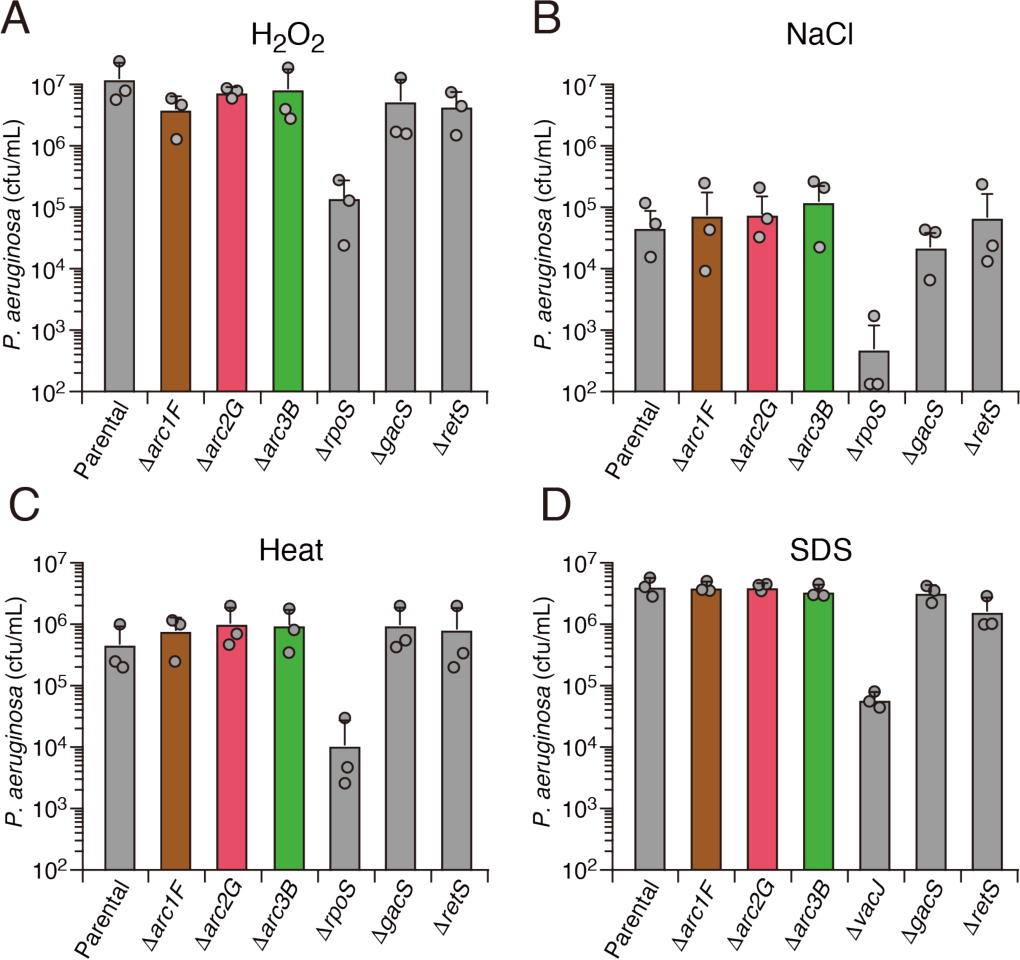
Arc1-3 do not contribute to the survival of *P. aeruginosa* exposed to common environmental stresses. Survival of the indicated *P. aeruginosa* strains during exposure to hydrogen peroxide (50 mM, 30 min) **(A),** high salinity (3M NaCl, 20 hours) **(B),** high temperature (55°C, 30 min) **(C)**, or detergent (0.5% (*w/v*) SDS) **(D)**.

**Supplemental Fig. 5.**
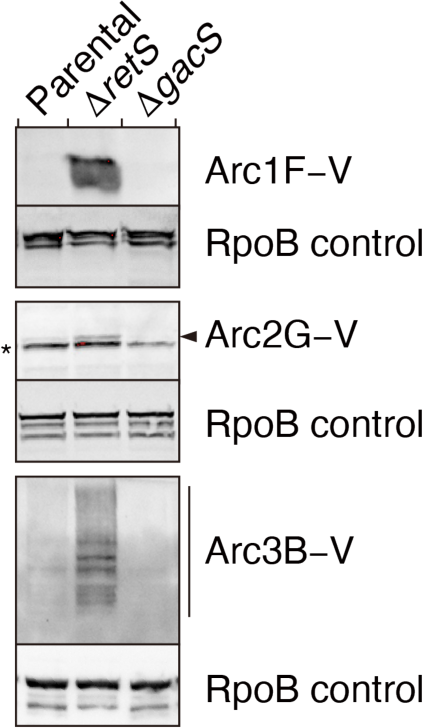
Arc1-3 are subject to tight regulation by the Gac/Rsm signaling pathway. Anti-VSV-G immunoblot analysis was used to probe levels of the indicated VSV–G-tagged (–V) Arc proteins encoded at their respective native genomic loci in *P. aeruginosa* wild-type, Δ*retS*, or Δ*gacS* backgrounds. The asterisk denotes a non-specific VSV-G antibody-reactive band. The arrow highlights the position of Arc2G–V.

**Supplemental Fig. 6.**
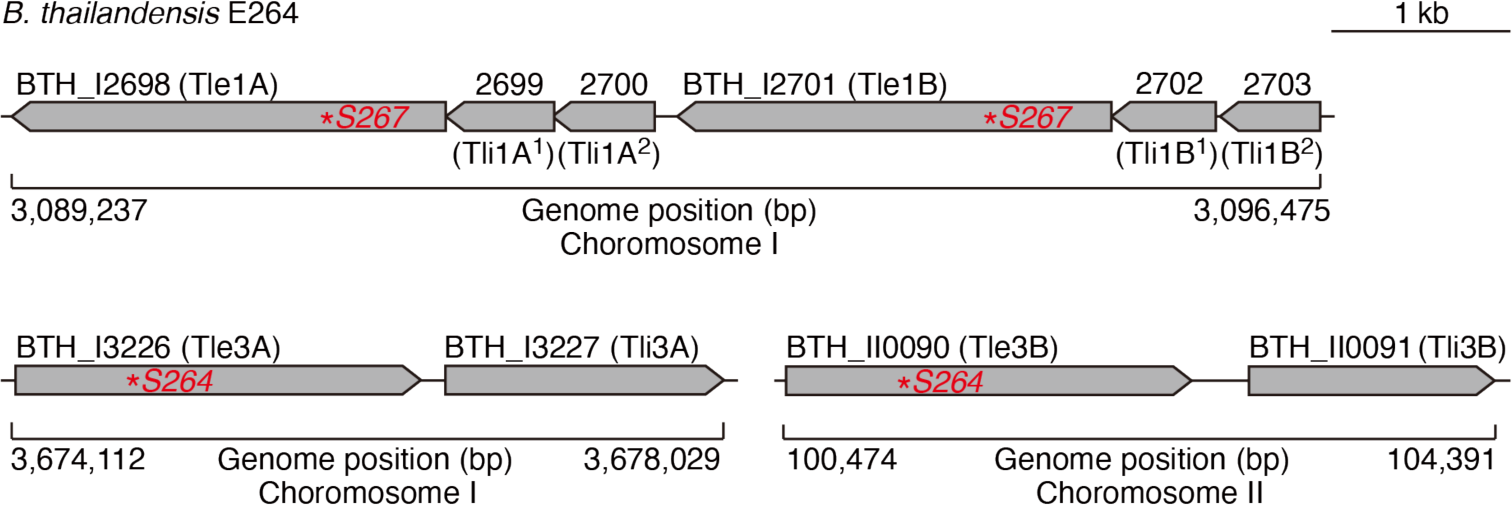
The loci encoding Tle1 and Tle3 are duplicated in *B. thai*. Schematic depicting the *tle1*/*tli1* (top) *tle3*/*tli3* (bottom) loci in the genome *B. thai* E264 strain. Predicted catalytic serine residues mutagenized in this study are indicated by red asterisks.

**Supplemental Fig. 7.**
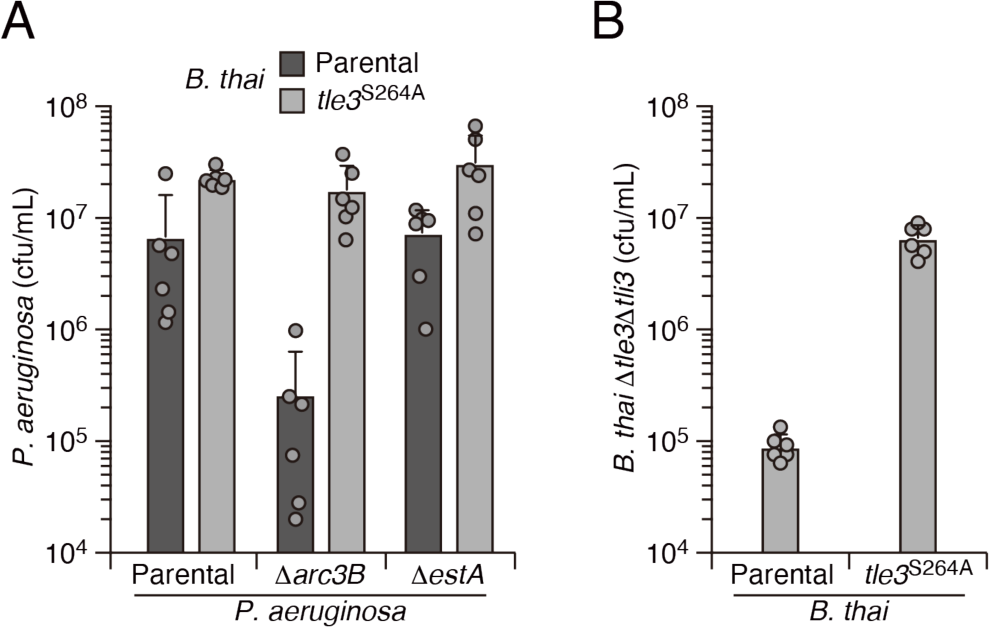
EstA is not involved in defense against Tle3 and Tle3 induces potent self-intoxication within *B. thai*. **(A)** Recovery of the indicated *P. aeruginosa* strains following growth competition against *B. thai* wild-type or a strain lacking Tle3 activity. **(B)** Recovery of *B. thai* strain sensitized to Tle3 intoxication (Δ*tle3*Δ*tli3*) following growth competition against *B. thai* wild-type or a strain lacking Tle3 activity.

**Supplemental Fig. 8.**
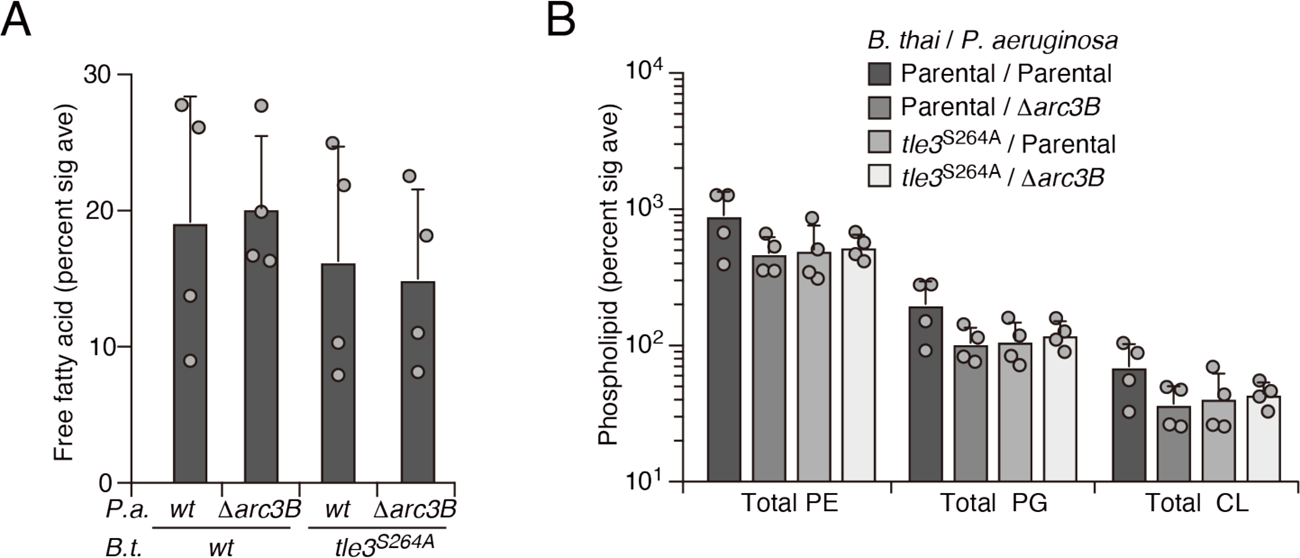
Tle3 intoxication does not impact free fatty acids nor intact parent phospholipids in *P. aeruginosa*. Mass spectrometric analysis of free fatty acid **(A)** and parent phospholipid species **(B)** within lipid extracts derived from the indicated mixtures.

**Supplemental Fig. 9.**
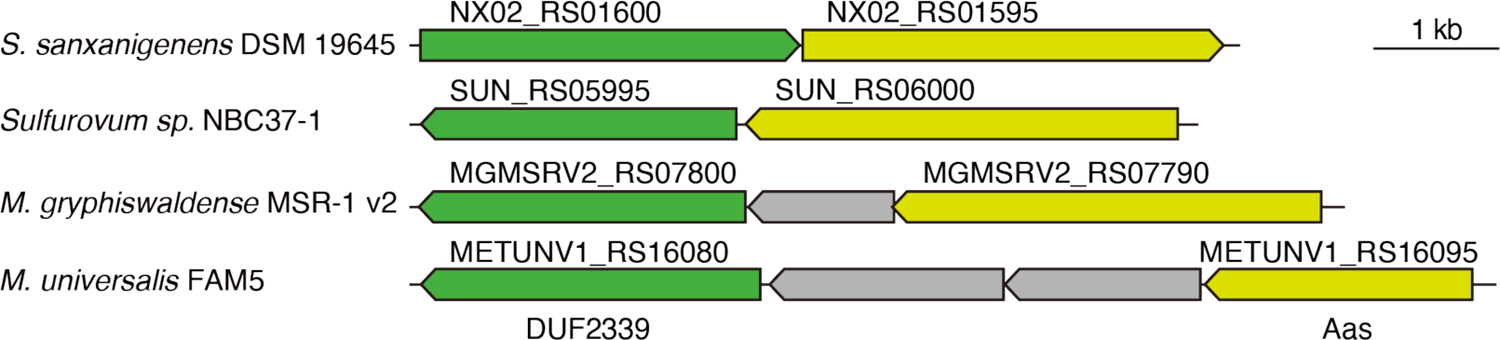
*Arc3B* genes are encoded adjacent to *aas* genes in diverse bacteria. Schematic depicting *arc3B* and *aas* loci in the indicated bacterial species.

**Supplemental Table 1.**
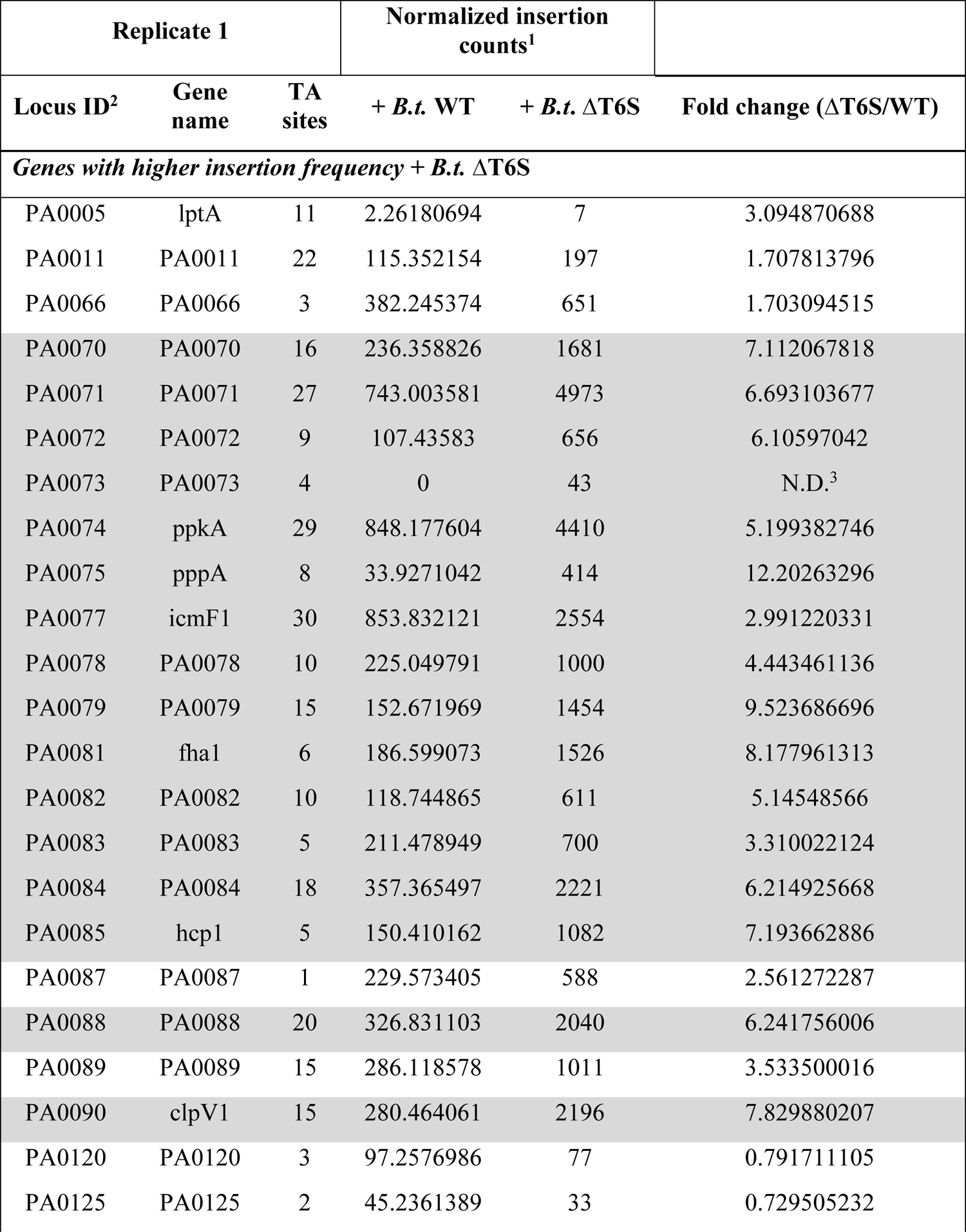
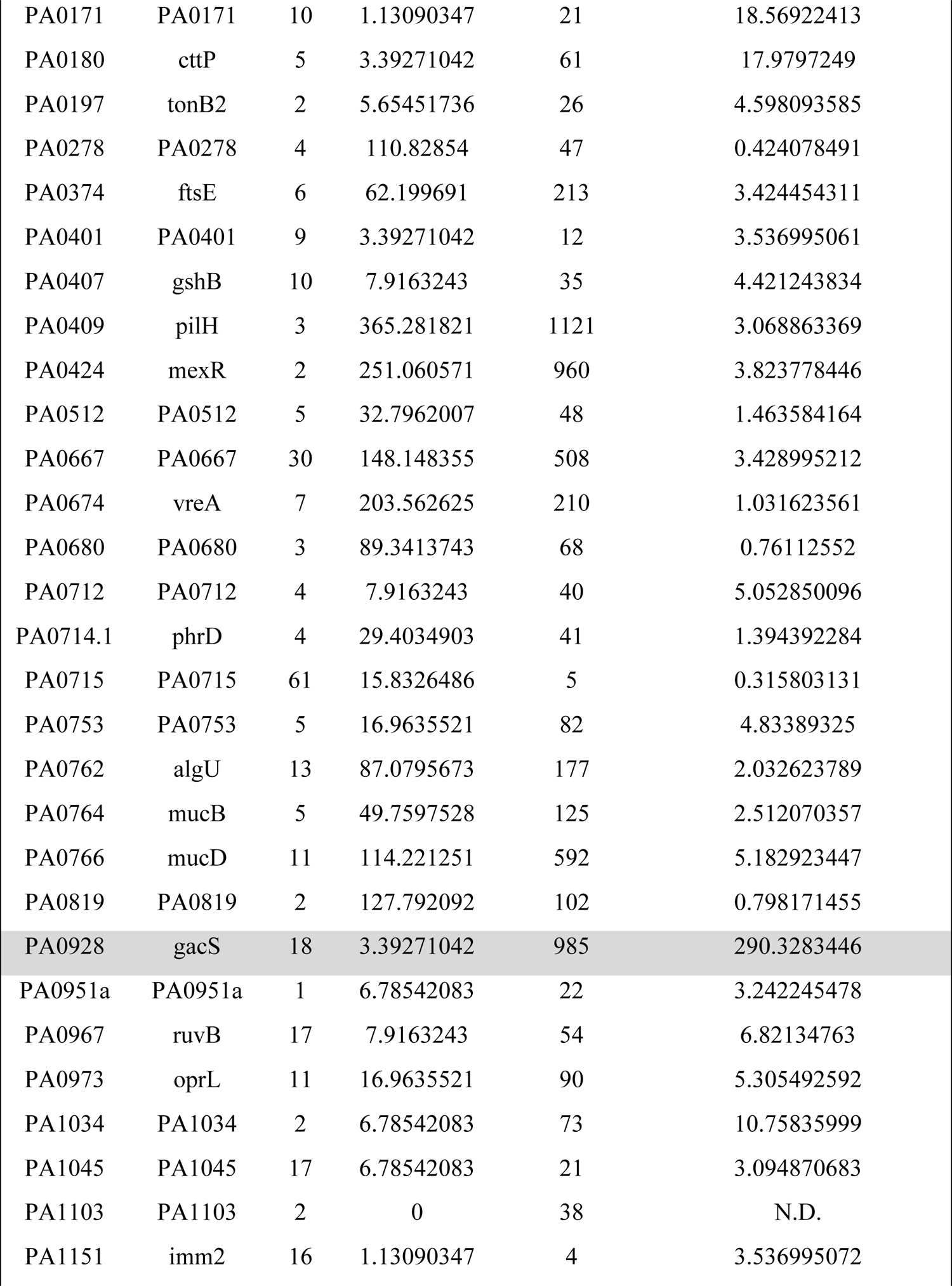
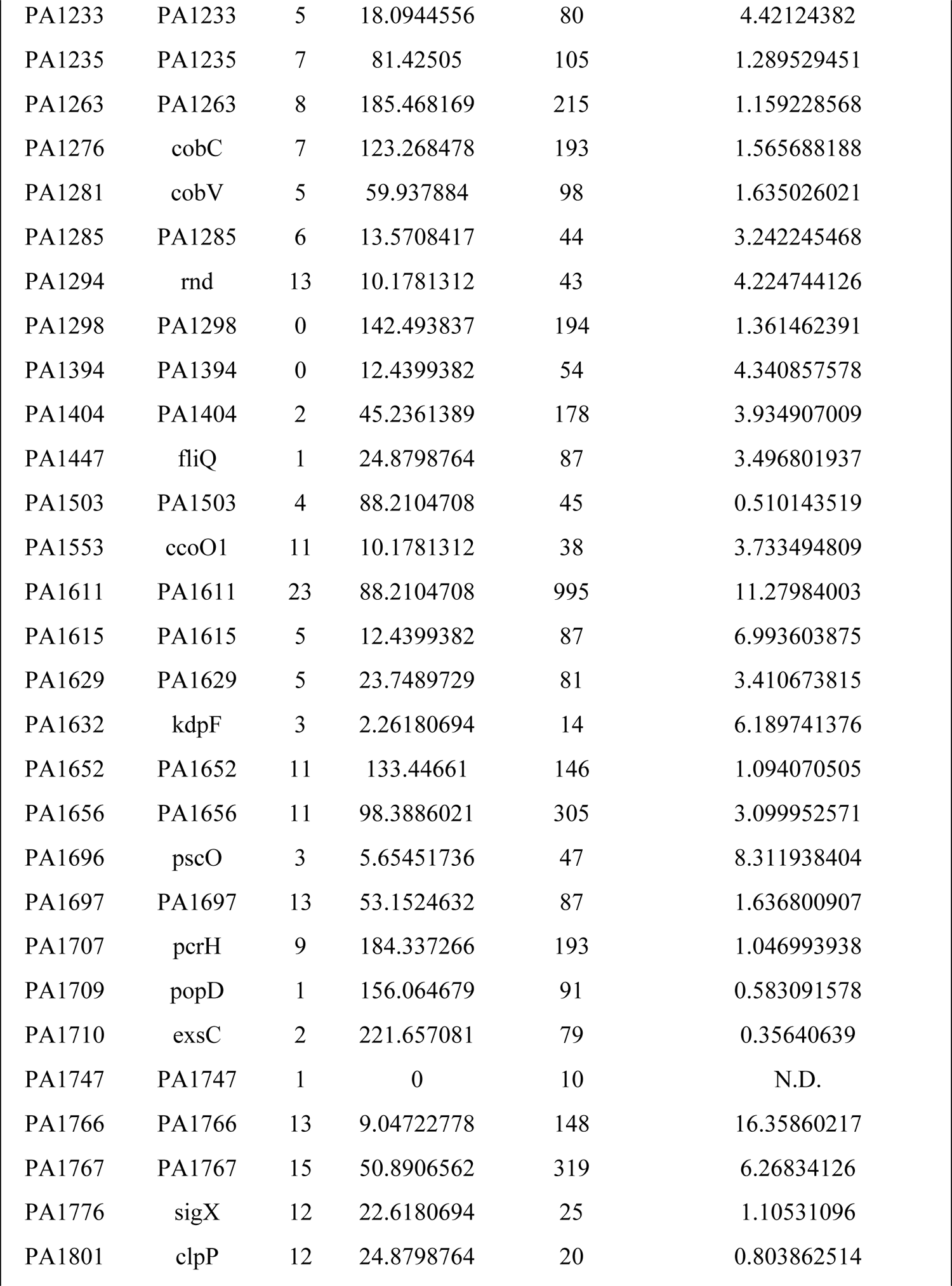
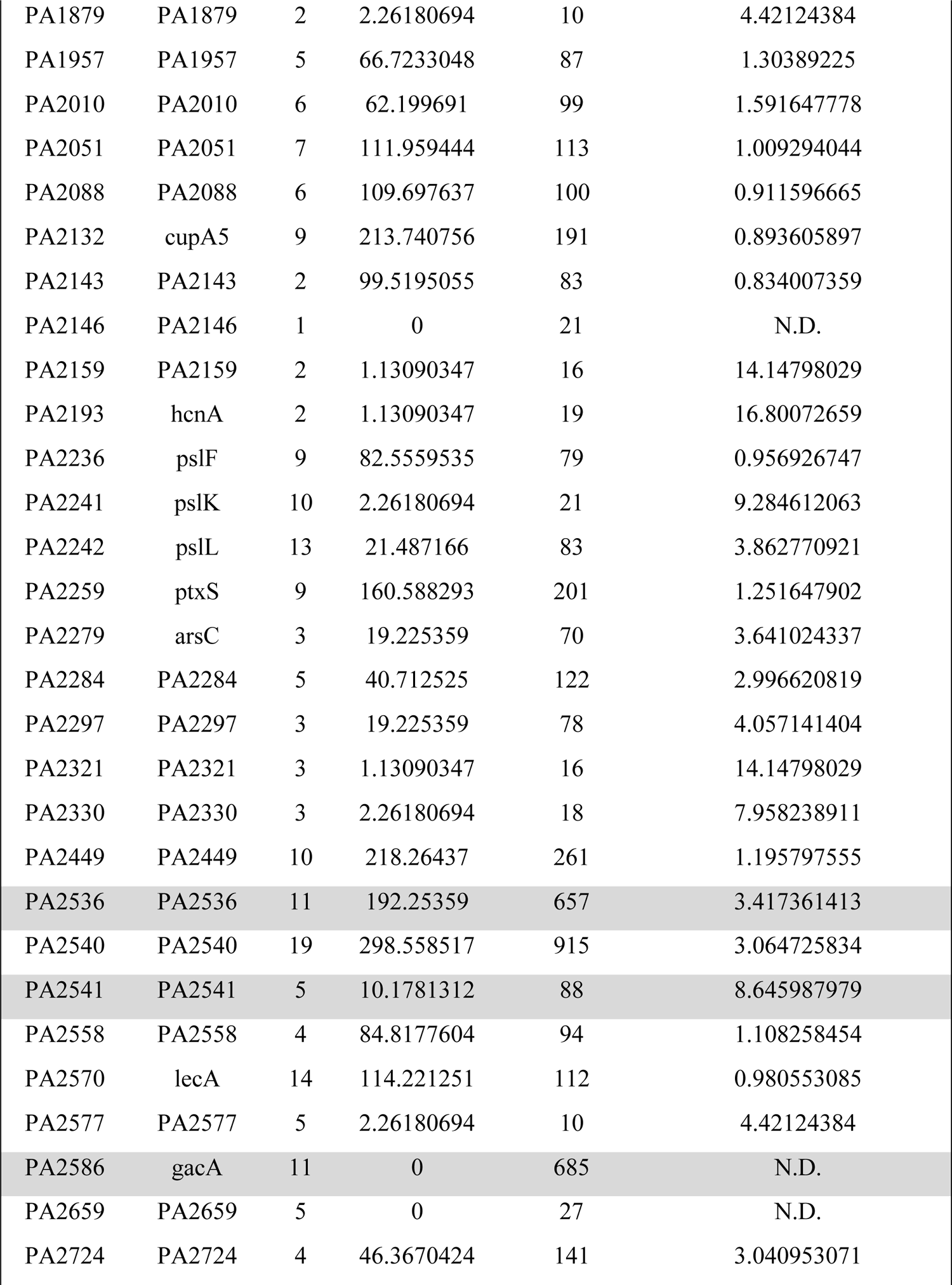
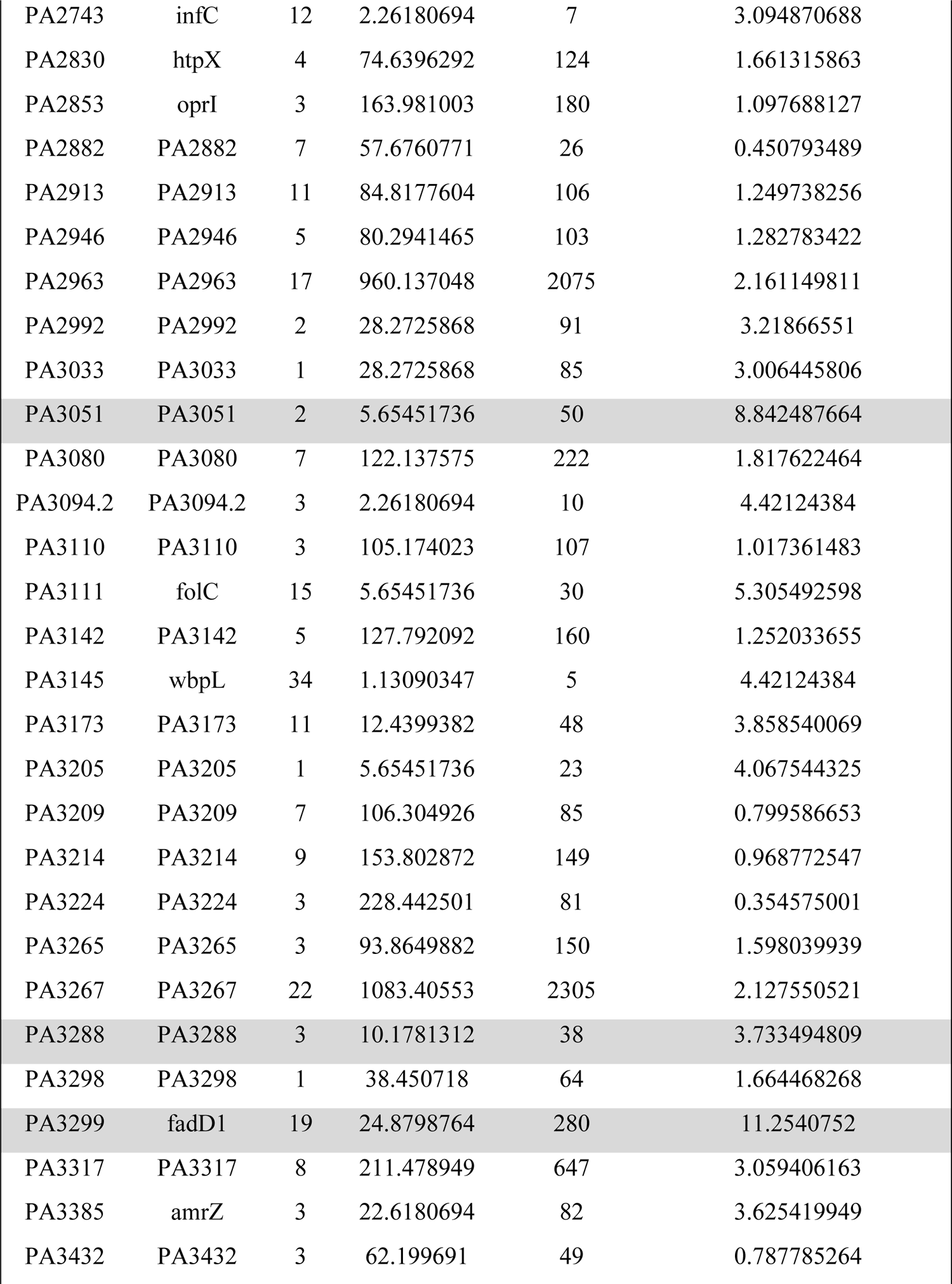
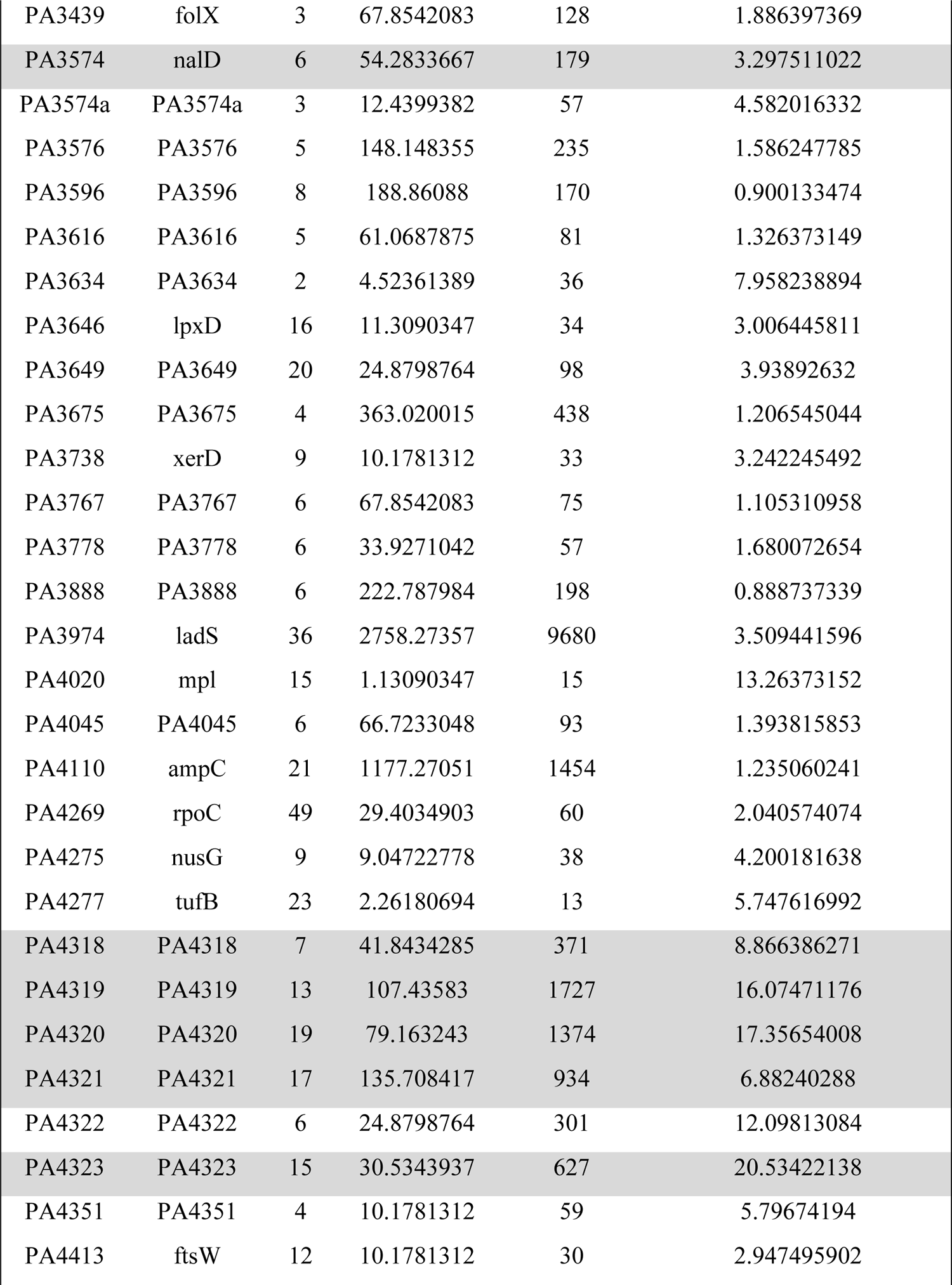
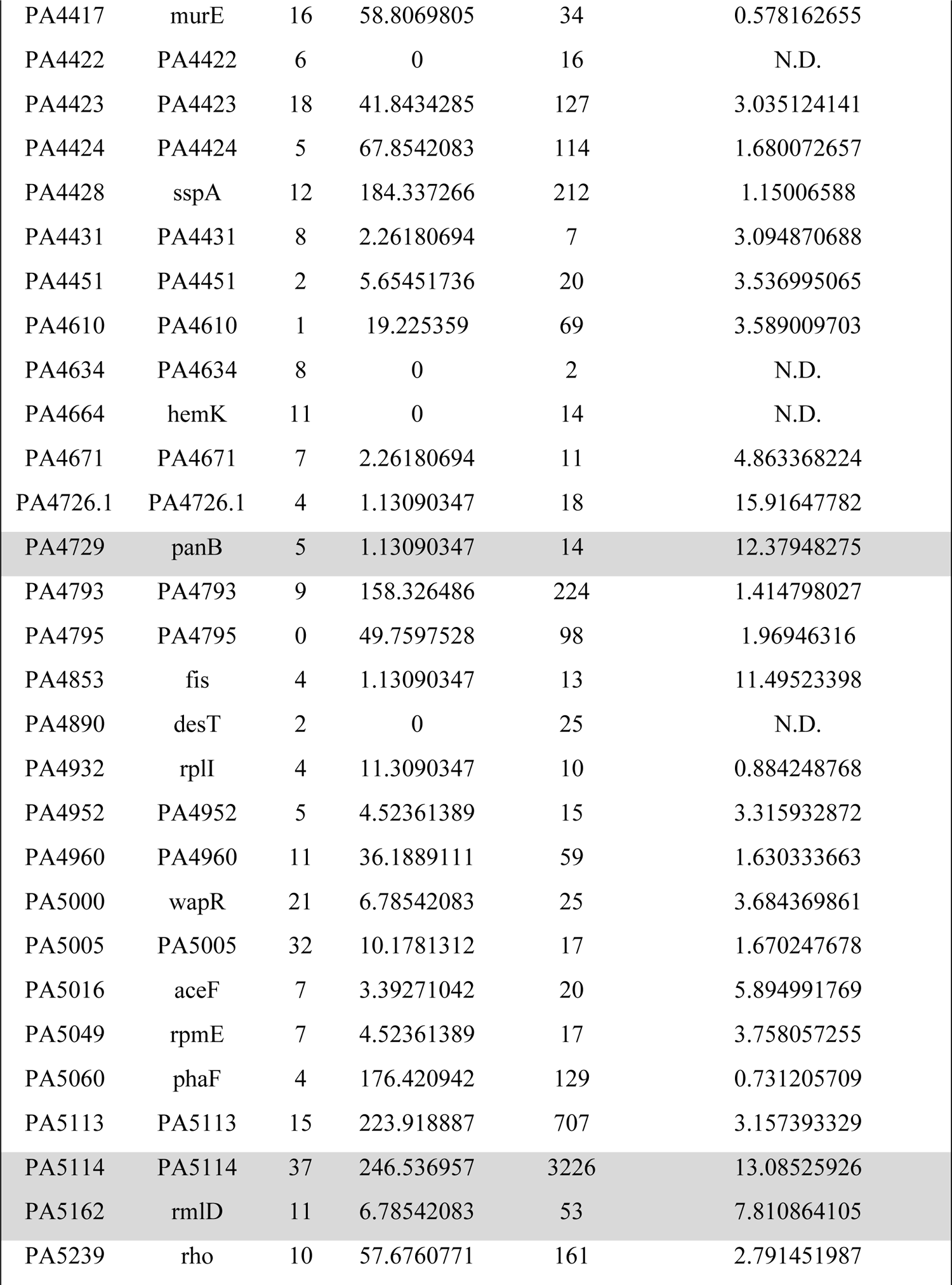
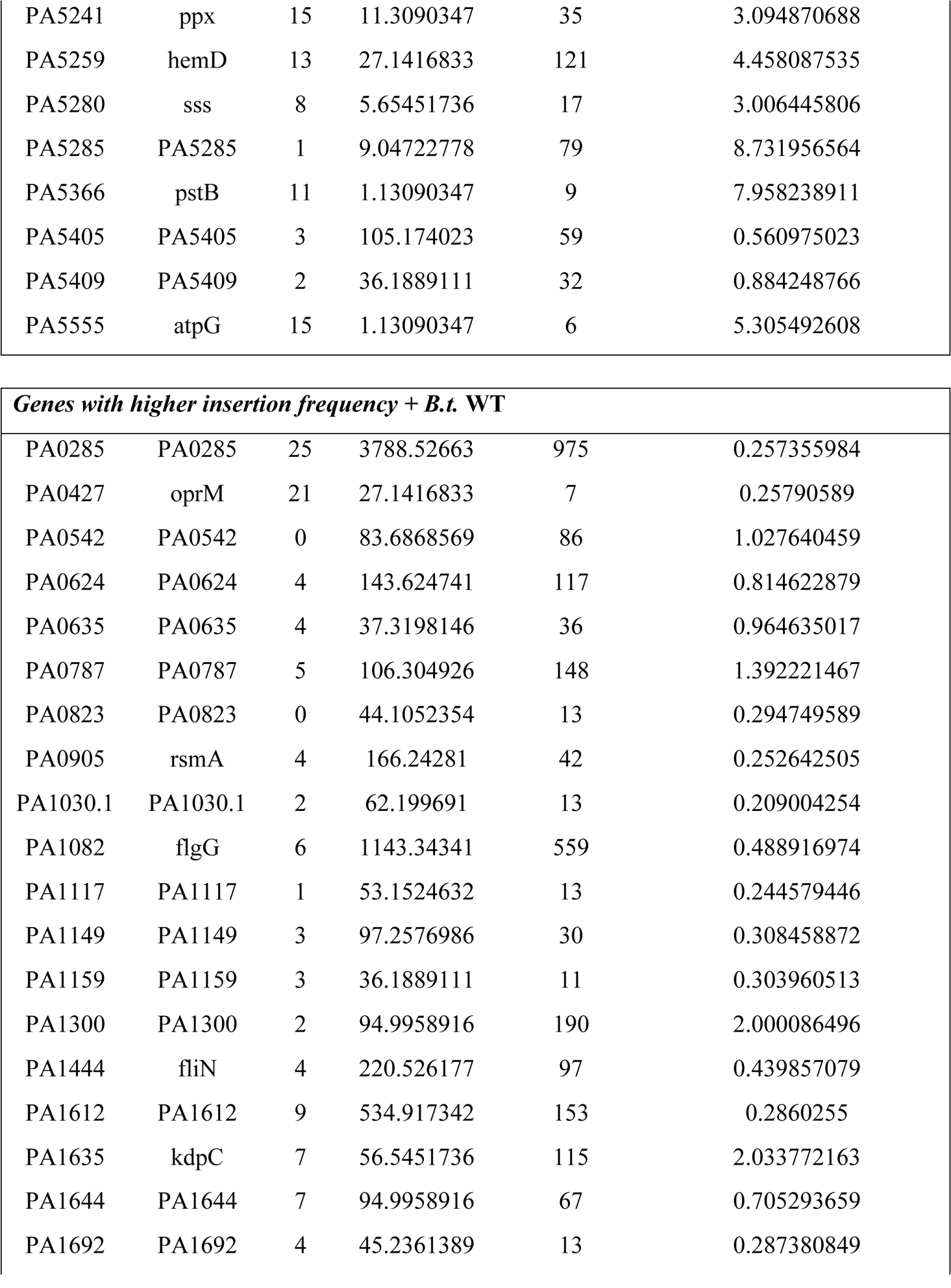
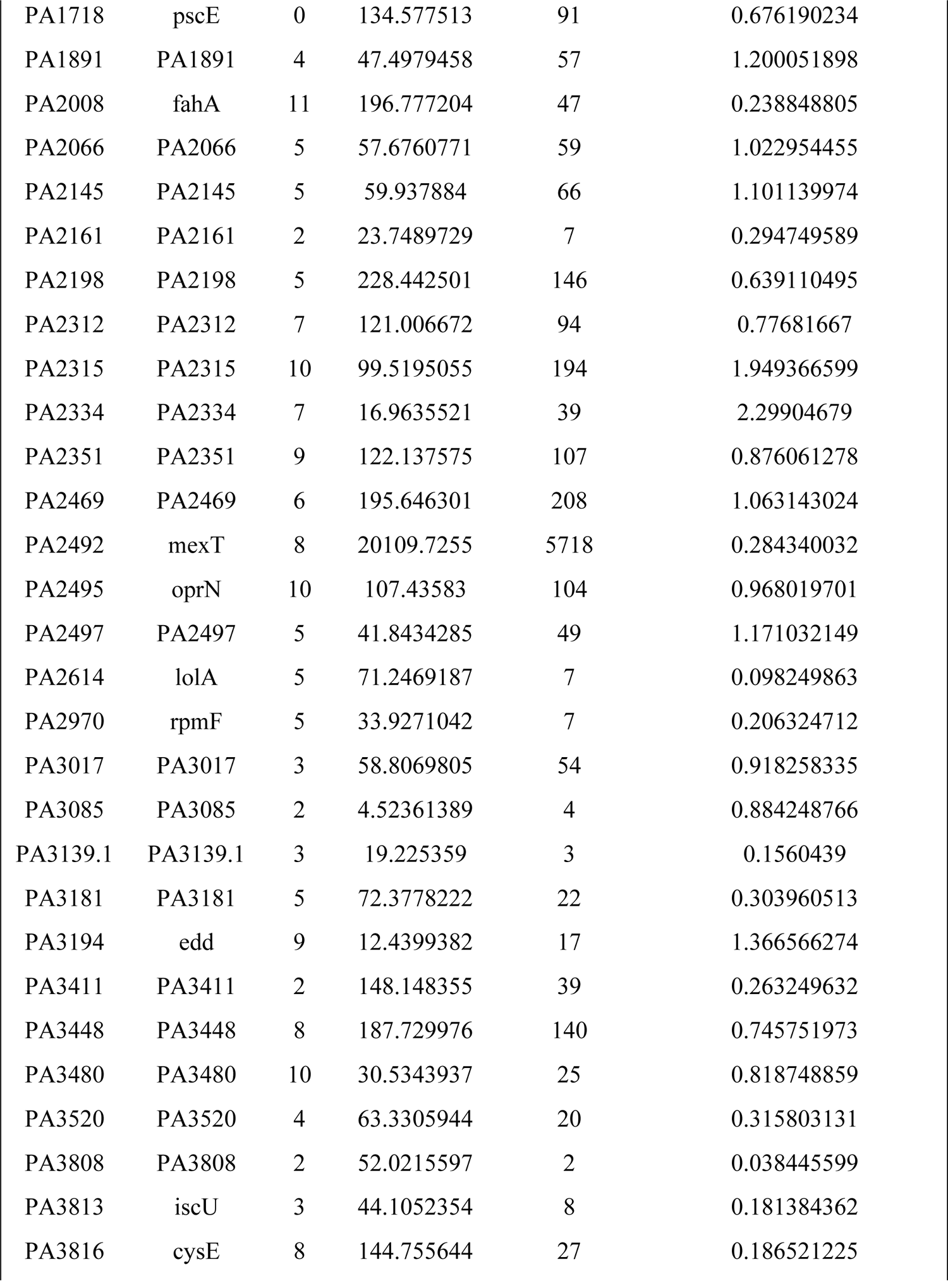
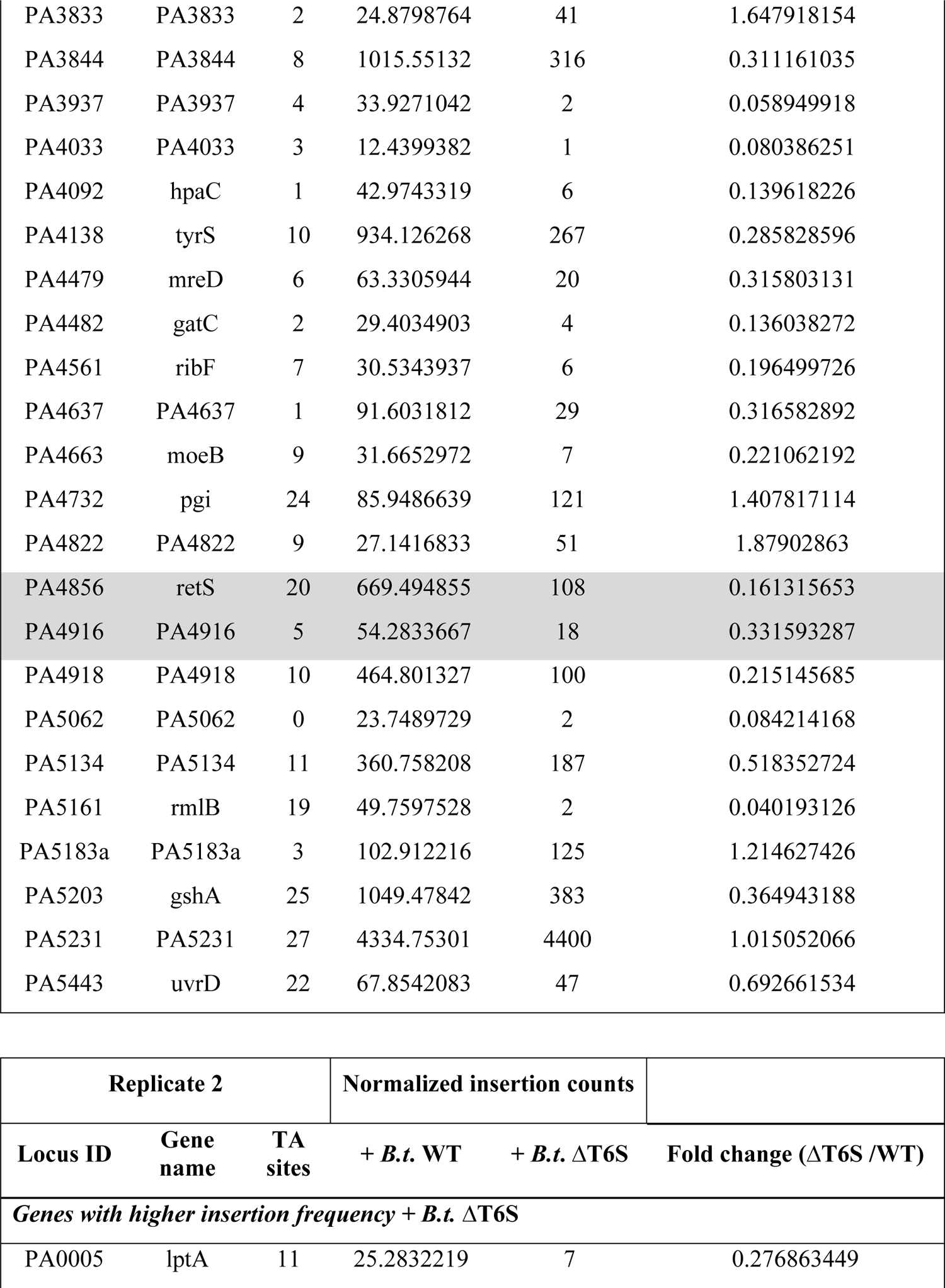
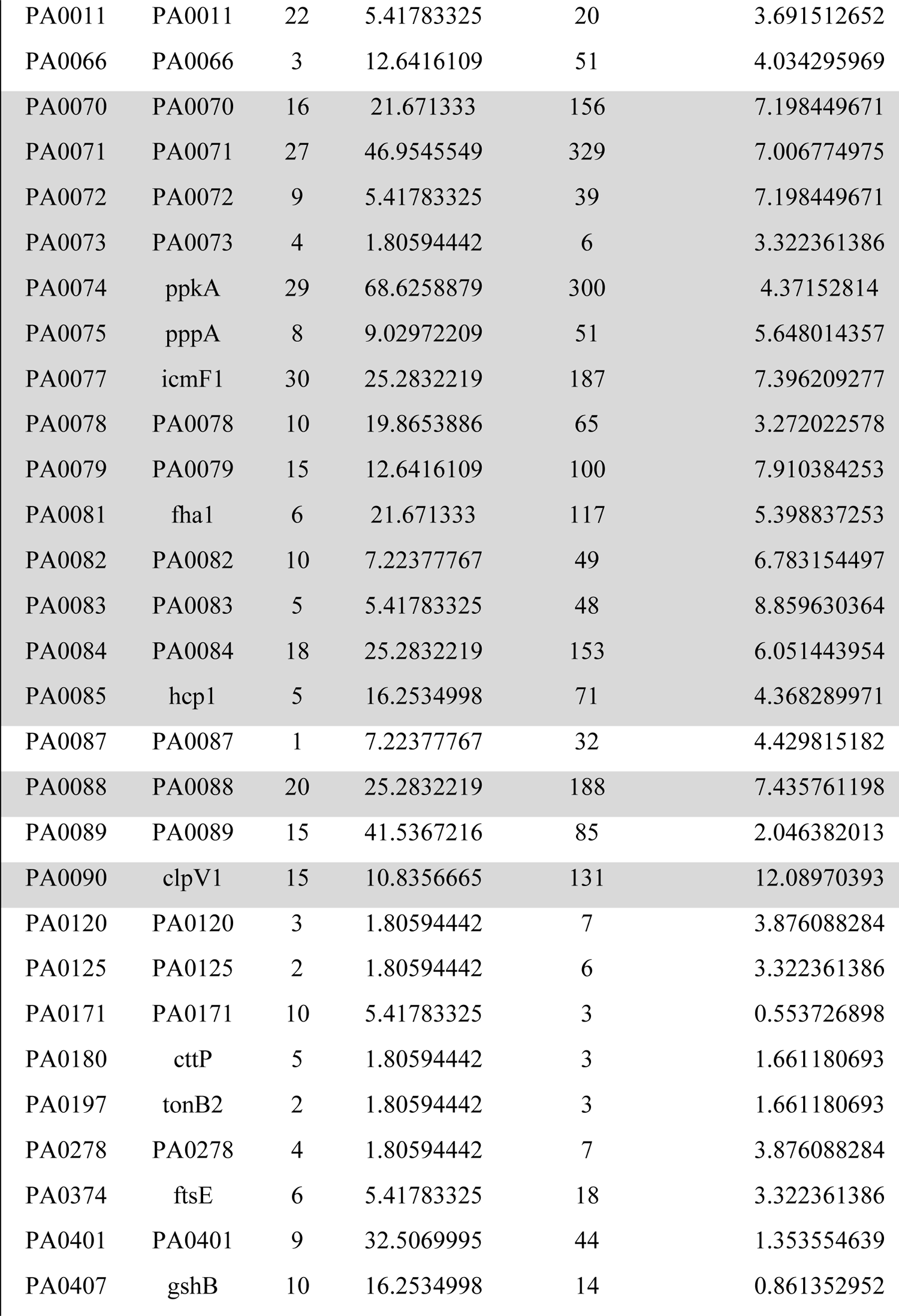
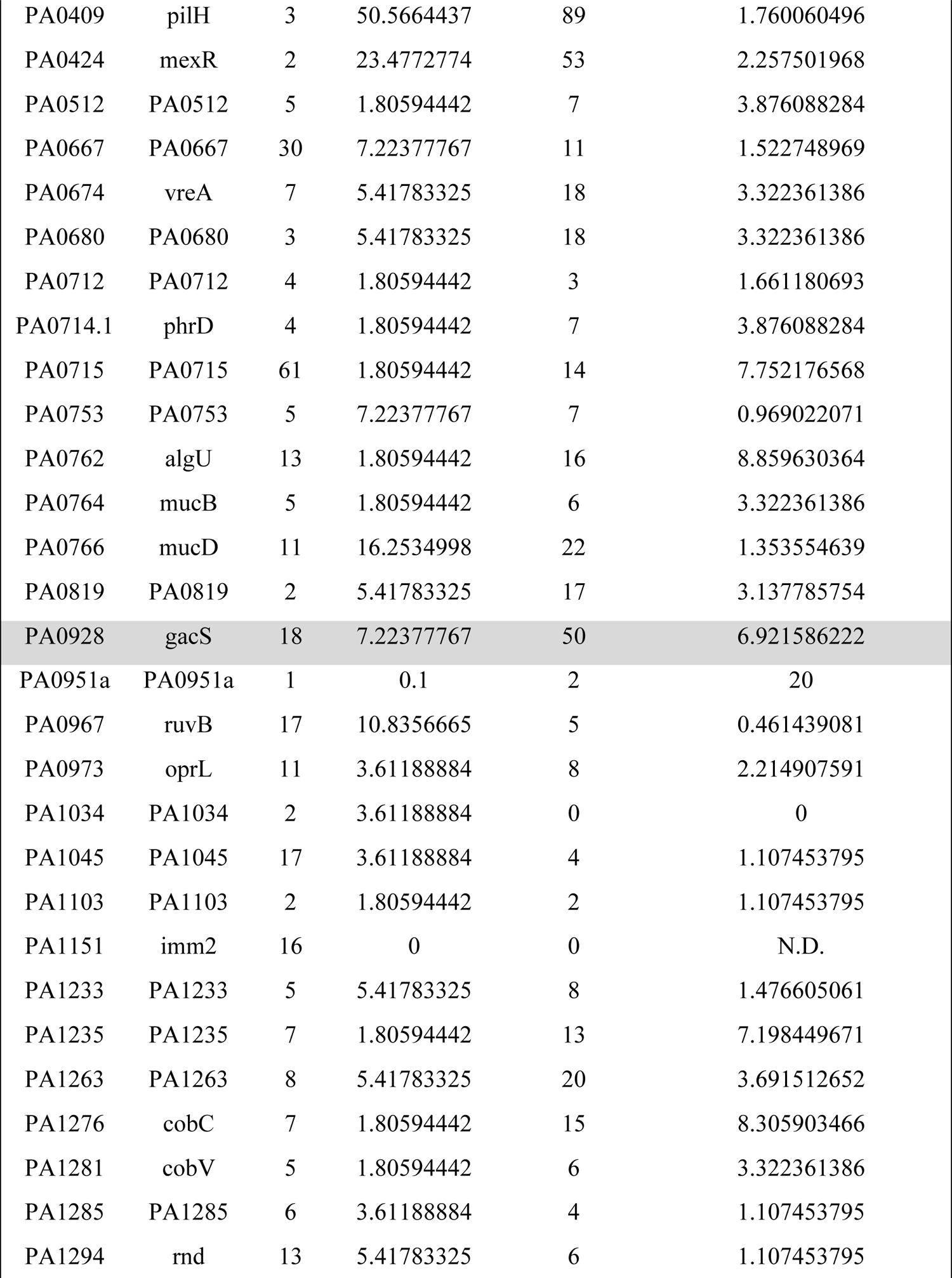
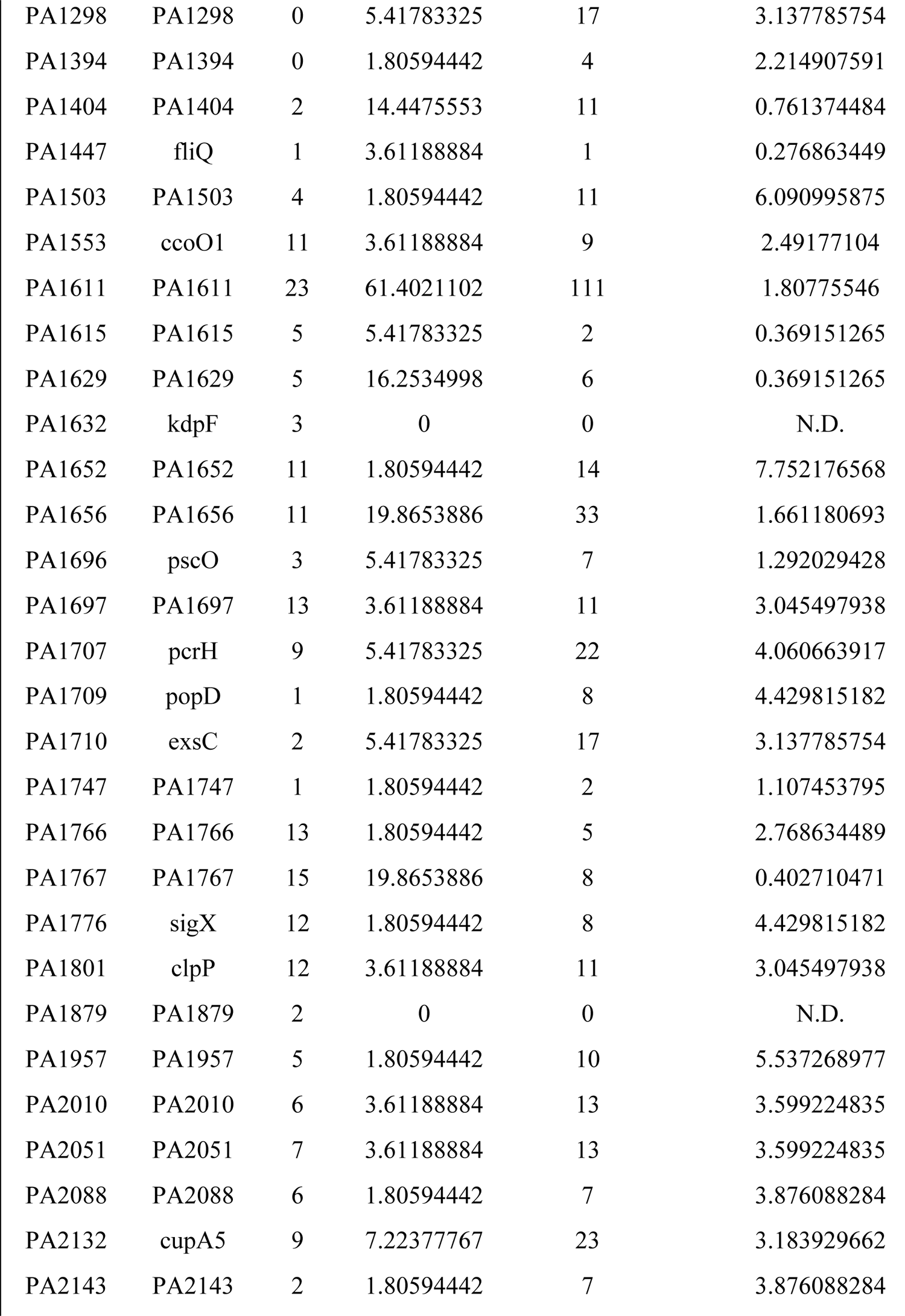
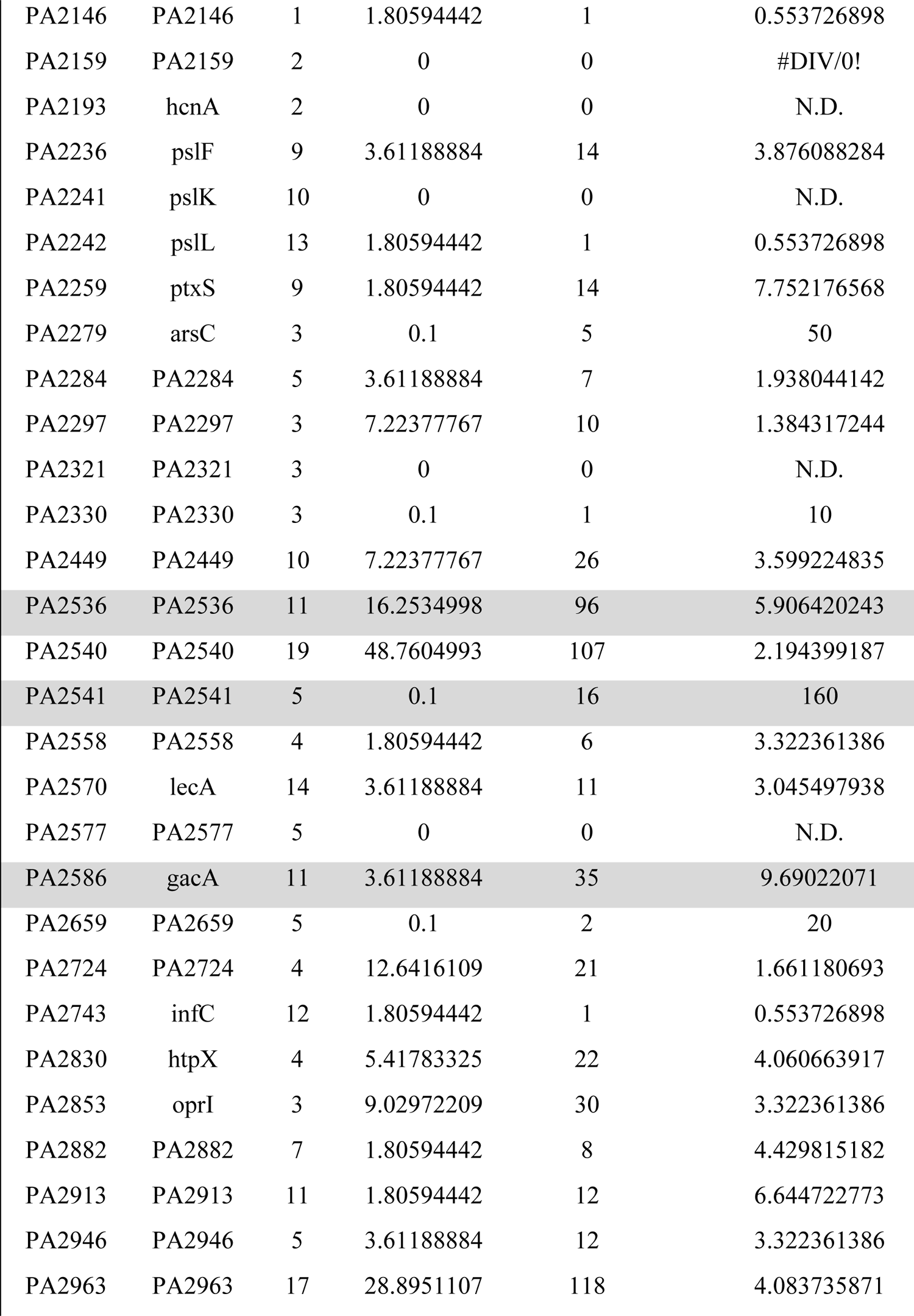
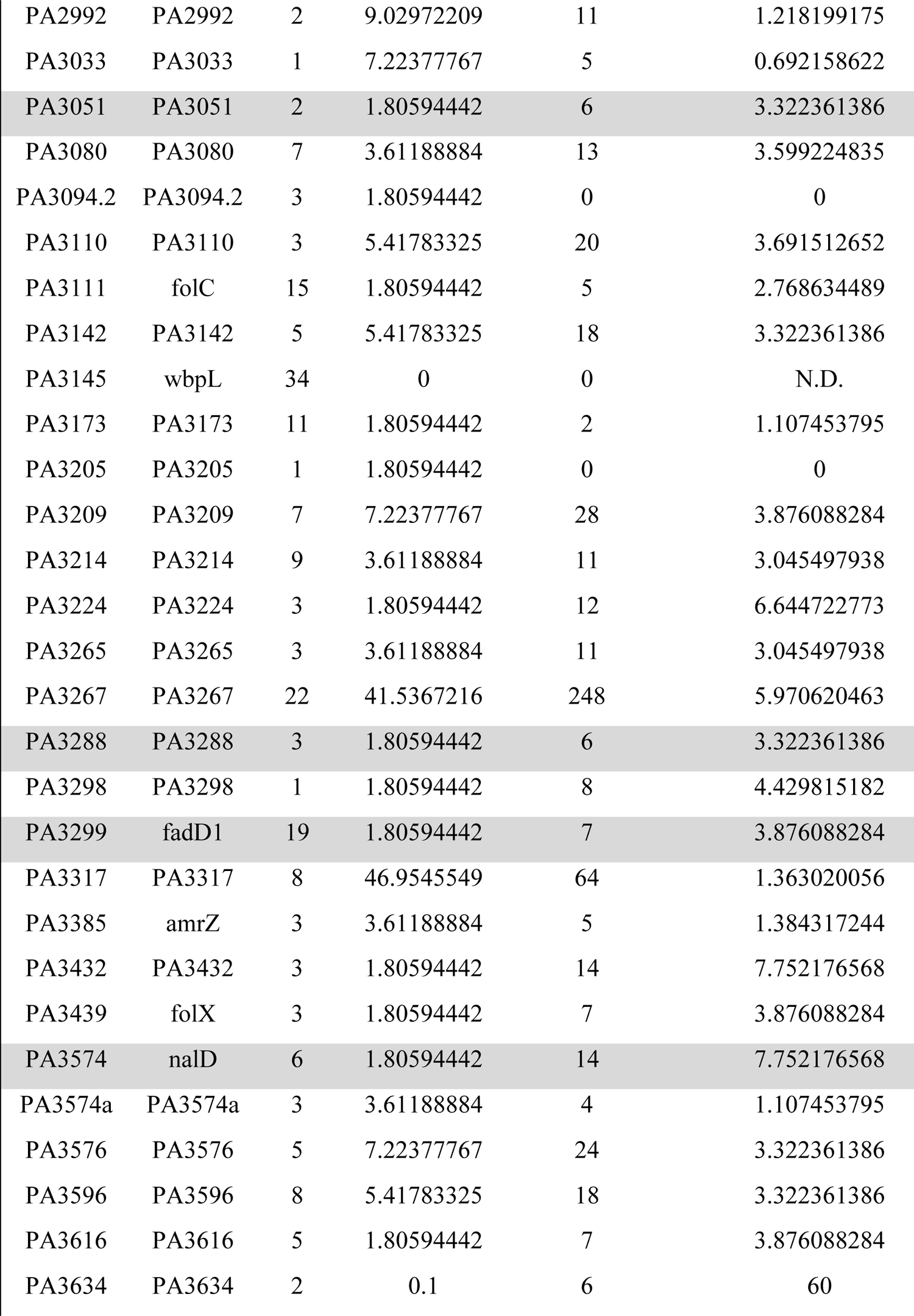
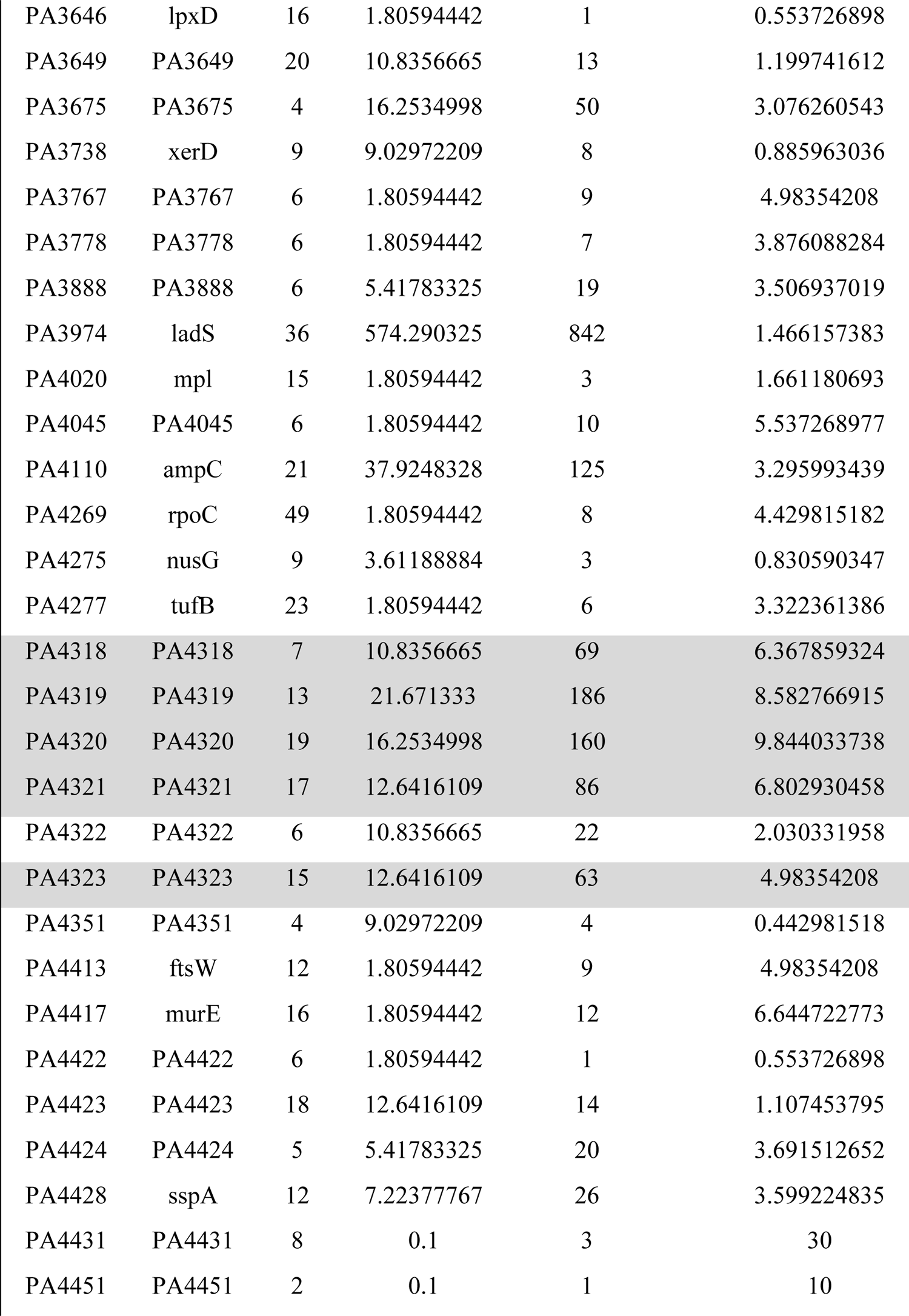
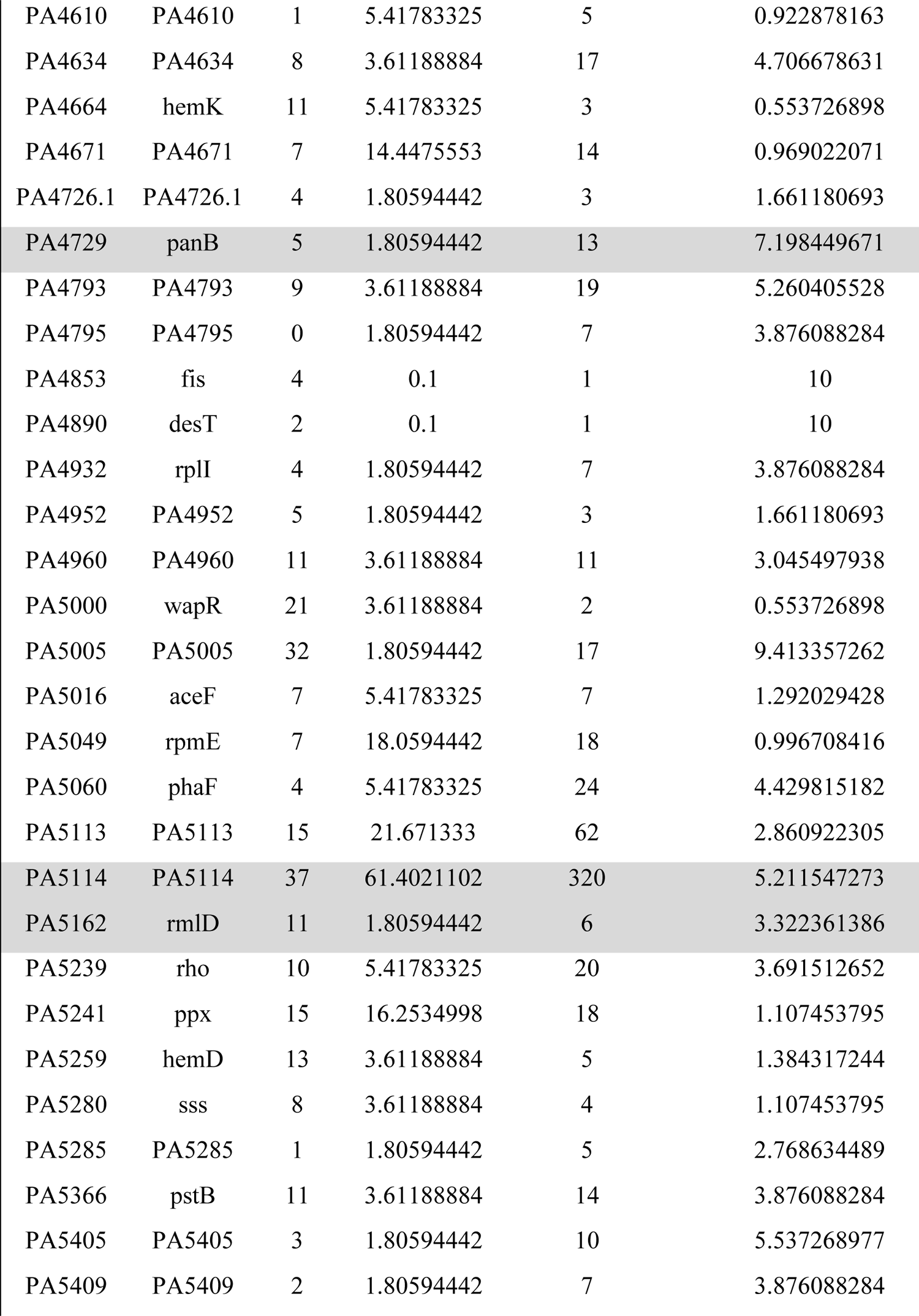
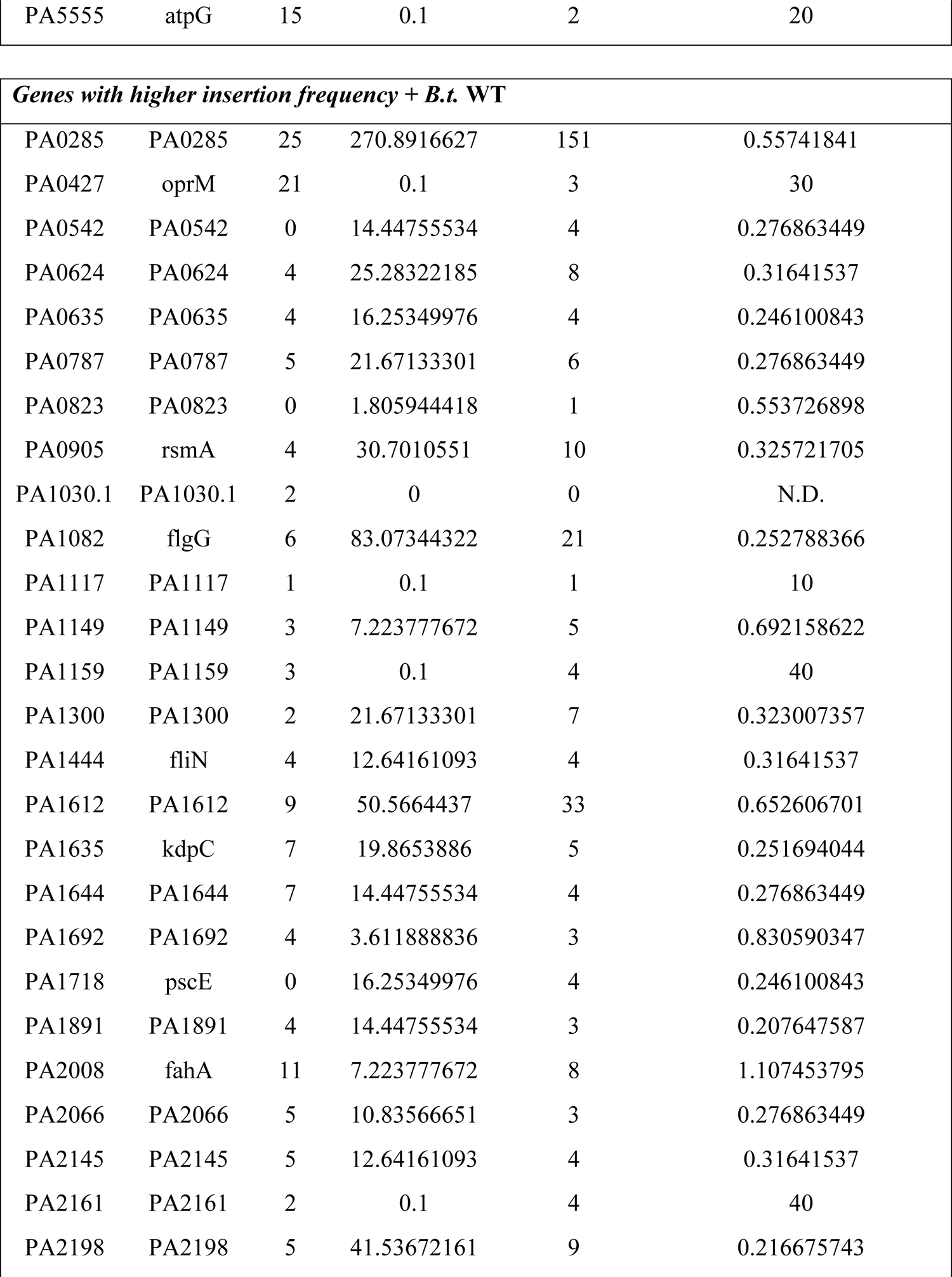
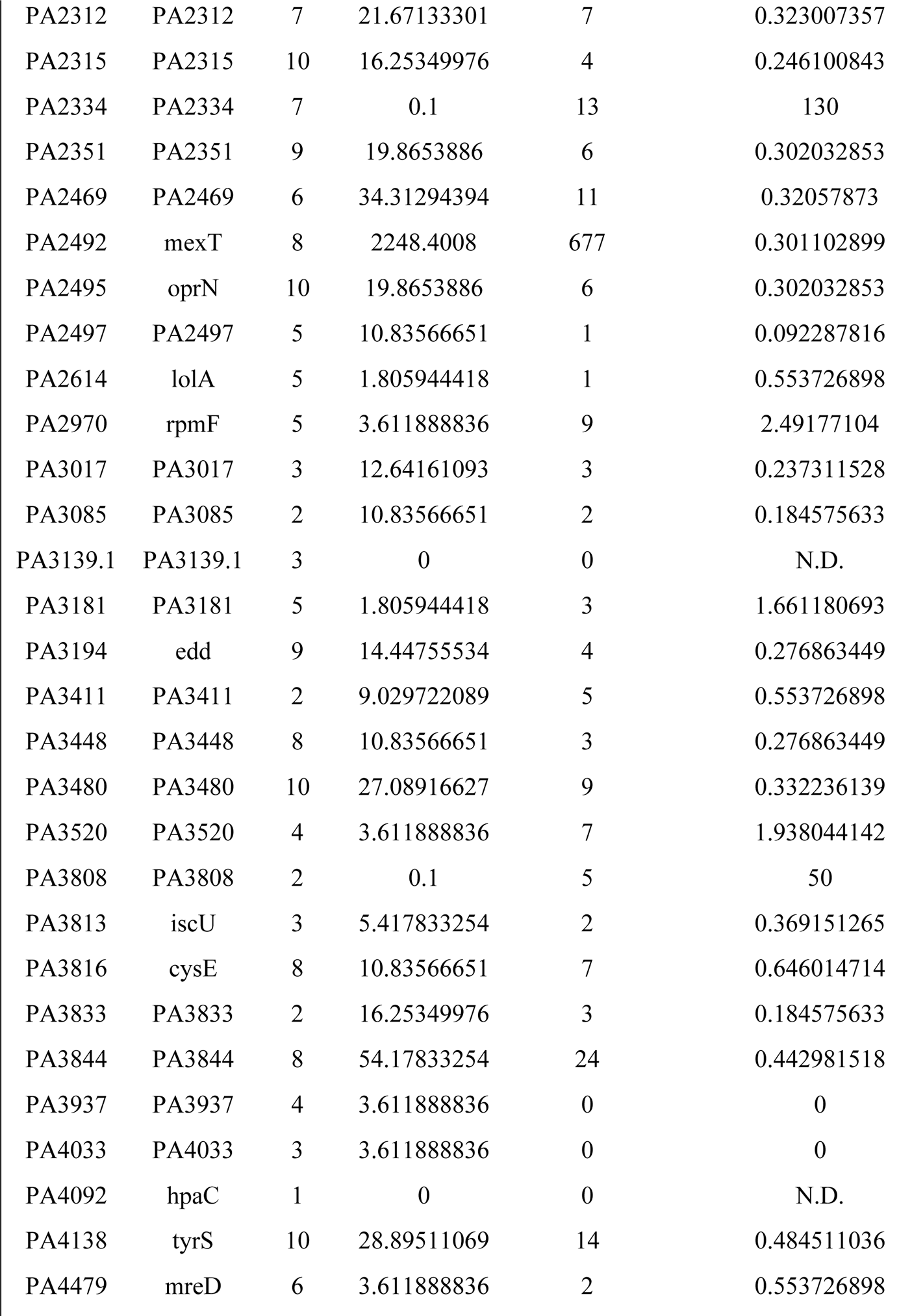
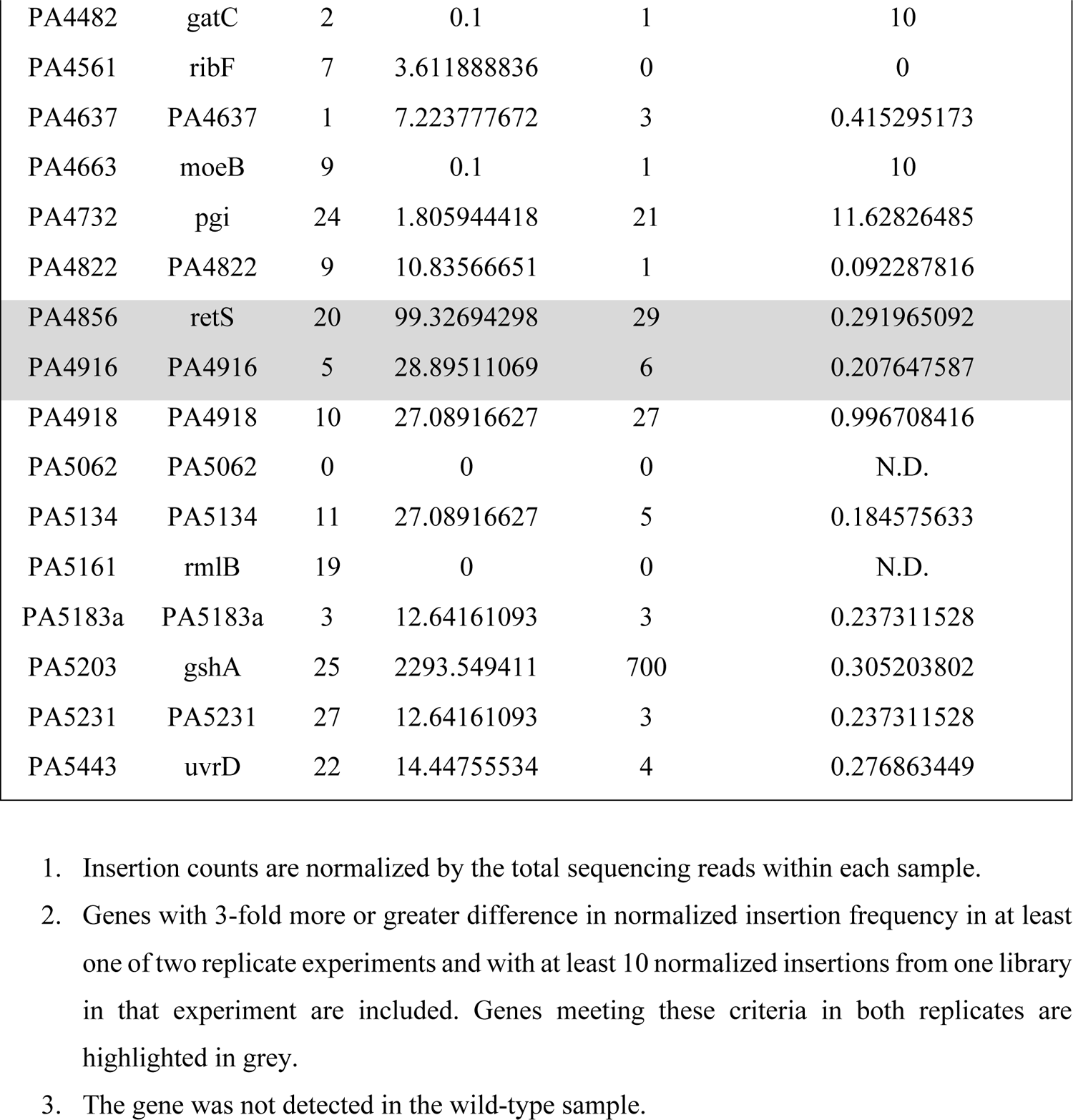
Transposon sequencing-based analysis of *P. aeruginosa* fitness determinants during antagonism by *B. thai*.

**Supplemental Table 2.** Transposon sequencing-based analysis of *P. aeruginosa* fitness determinants during antagonism by *B. thai*.

|  |  |  | Normalized insertion counts |  |  |
| --- | --- | --- | --- | --- | --- |
| Locus ID | Gene name | TA sites | + *B.t.* WT | + *B.t. tle3*<sup>S264A</sup> | Fold change (*tle3*<sup>S264A</sup>/WT) |
| ***Genes with higher insertion frequency + *B.t. tle3*<sup>S264A</sup>*** |  |  |  |  |  |
| PA0005 | lptA | 11 | 2 | 17.00958 | 8.504789999 |
| PA0073 | PA0073 | 4 | 0.1 | 34.01916 | 340.1916 |
| PA0075 | pppA | 8 | 30 | 100.5111545 | 3.350371818 |
| PA0079 | PA0079 | 15 | 135 | 415.9615472 | 3.081196646 |
| PA0171 | PA0171 | 10 | 1 | 7.731627272 | 7.731627272 |
| PA0180 | cttP | 5 | 3 | 52.5750654 | 17.5250218 |
| PA0197 | tonB2 | 2 | 5 | 24.74120727 | 4.948241454 |
| PA0307 | PA0307 | 7 | 65 | 231.9488182 | 3.568443356 |
| PA0316 | serA | 13 | 21 | 85.04789999 | 4.0499 |
| PA0402 | pyrB | 9 | 1 | 12.37060363 | 12.37060363 |
| PA0407 | gshB | 10 | 7 | 26.28753272 | 3.755361818 |
| PA0466 | PA0466 | 1 | 0.1 | 27.83385818 | 278.3385818 |
| PA0526 | PA0526 | 1 | 38 | 160.8178473 | 4.232048612 |
| PA0553 | PA0553 | 2 | 0.1 | 12.37060363 | 123.7060363 |
| PA0712 | PA0712 | 4 | 7 | 24.74120727 | 3.534458181 |
| PA0721 | PA0721 | 4 | 0.1 | 12.37060363 | 123.7060363 |
| PA0766 | mucD | 11 | 101 | 400.4982927 | 3.965329631 |
| PA0909 | PA0909 | 5 | 34 | 108.2427818 | 3.18361123 |
| PA0928 | gacS | 18 | 3 | 9.277952726 | 3.092650909 |
| PA0945 | purM | 13 | 32 | 123.7060363 | 3.865813636 |
| PA0951a | PA0951a | 1 | 6 | 57.21404181 | 9.535673635 |
| PA0960 | PA0960 | 0 | 10 | 34.01916 | 3.401916 |
| PA0973 | oprL | 11 | 15 | 46.38976363 | 3.092650909 |
| PA0974 | PA0974 | 16 | 70 | 228.8561672 | 3.269373818 |
| PA1013 | purC | 3 | 2 | 15.46325454 | 7.731627272 |
| PA1034 | PA1034 | 2 | 6 | 40.20446181 | 6.700743636 |
| PA1101 | fliF | 16 | 421 | 1456.638578 | 3.459949116 |
| PA1103 | PA1103 | 2 | 0.1 | 35.56548545 | 355.6548545 |
| PA1121 | yfiR | 7 | 21 | 83.50157454 | 3.976265454 |
| PA1285 | PA1285 | 6 | 12 | 43.29711272 | 3.608092727 |
| PA1394 | PA1394 | 0 | 11 | 37.1118109 | 3.373800991 |
| PA1443 | fliM | 7 | 80 | 366.4791327 | 4.580989159 |
| PA1547 | PA1547 | 5 | 0.1 | 10.82427818 | 108.2427818 |
| PA1548 | PA1548 | 3 | 0.1 | 30.92650909 | 309.2650909 |
| PA1549 | PA1549 | 19 | 6 | 20.10223091 | 3.350371818 |
| PA1550 | PA1550 | 6 | 3 | 27.83385818 | 9.277952726 |
| PA1554 | ccoN1 | 25 | 211 | 638.6324127 | 3.026693899 |
| PA1615 | PA1615 | 5 | 11 | 35.56548545 | 3.23322595 |
| PA1629 | PA1629 | 5 | 21 | 80.40892363 | 3.828996363 |
| PA1632 | kdpF | 3 | 2 | 10.82427818 | 5.41213909 |
| PA1696 | pscO | 3 | 5 | 18.55590545 | 3.71118109 |
| PA1715 | pscB | 2 | 2 | 6.185301817 | 3.092650909 |
| PA1766 | PA1766 | 13 | 8 | 54.1213909 | 6.765173863 |
| PA1796.2 | PA1796.2 | 2 | 0.1 | 13.91692909 | 139.1692909 |
| PA1879 | PA1879 | 2 | 2 | 10.82427818 | 5.41213909 |
| PA2146 | PA2146 | 1 | 0.1 | 24.74120727 | 247.4120727 |
| PA2173 | PA2173 | 0 | 0.1 | 34.01916 | 340.1916 |
| PA2193 | hcnA | 2 | 1 | 12.37060363 | 12.37060363 |
| PA2297 | PA2297 | 3 | 17 | 58.76036727 | 3.456492192 |
| PA2330 | PA2330 | 3 | 2 | 21.64855636 | 10.82427818 |
| PA2425 | pvdG | 13 | 8 | 24.74120727 | 3.092650909 |
| PA2541 | PA2541 | 5 | 9 | 46.38976363 | 5.154418181 |
| PA2816 | PA2816 | 1 | 31 | 114.4280836 | 3.691228504 |
| PA2852 | PA2852 | 9 | 6 | 20.10223091 | 3.350371818 |
| PA2898 | PA2898 | 1 | 4 | 17.00958 | 4.252395 |
| PA2947 | PA2947 | 2 | 18 | 69.58464545 | 3.865813636 |
| PA2964 | pabC | 5 | 5 | 27.83385818 | 5.566771636 |
| PA2992 | PA2992 | 2 | 25 | 119.06706 | 4.762682399 |
| PA3094.2 | PA3094.2 | 3 | 2 | 12.37060363 | 6.185301817 |
| PA3099 | xcpV | 1 | 8 | 29.38018363 | 3.672522954 |
| PA3111 | folC | 15 | 5 | 20.10223091 | 4.020446181 |
| PA3173 | PA3173 | 11 | 11 | 44.84343818 | 4.076676198 |
| PA3202 | PA3202 | 3 | 10 | 34.01916 | 3.401916 |
| PA3299 | fadD1 | 19 | 22 | 98.96482908 | 4.498401322 |
| PA3382 | phnE | 4 | 32 | 128.3450127 | 4.010781647 |
| PA3509 | PA3509 | 6 | 0.1 | 27.83385818 | 278.3385818 |
| PA3574a | PA3574a | 3 | 11 | 58.76036727 | 5.34185157 |
| PA3738 | xerD | 9 | 9 | 34.01916 | 3.779906666 |
| PA3815 | iscR | 7 | 13 | 51.02873999 | 3.925287692 |
| PA4232 | ssb | 6 | 0.1 | 12.37060363 | 123.7060363 |
| PA4275 | nusG | 9 | 8 | 61.85301817 | 7.731627272 |
| PA4351 | PA4351 | 4 | 9 | 41.75078727 | 4.638976363 |
| PA4357 | PA4357 | 1 | 21 | 80.40892363 | 3.828996363 |
| PA4431 | PA4431 | 8 | 2 | 17.00958 | 8.504789999 |
| PA4451 | PA4451 | 2 | 5 | 29.38018363 | 5.876036727 |
| PA4726 | cbrB | 14 | 181 | 889.1371363 | 4.912359869 |
| PA4726.1 | PA4726.1 | 4 | 1 | 12.37060363 | 12.37060363 |
| PA4752 | ftsJ | 9 | 9 | 32.47283454 | 3.608092727 |
| PA4782 | PA4782 | 2 | 39 | 119.06706 | 3.053001538 |
| PA4853 | fis | 4 | 1 | 40.20446181 | 40.20446181 |
| PA4890 | desT | 2 | 0.1 | 12.37060363 | 123.7060363 |
| PA4894 | PA4894 | 3 | 8 | 34.01916 | 4.252395 |
| PA5000 | wapR | 21 | 6 | 24.74120727 | 4.123534545 |
| PA5015 | aceE | 38 | 58 | 270.6069545 | 4.665637147 |
| PA5016 | aceF | 7 | 3 | 18.55590545 | 6.185301817 |
| PA5113 | PA5113 | 15 | 198 | 632.4471108 | 3.194177327 |
| PA5114 | PA5114 | 37 | 218 | 3002.964032 | 13.77506437 |
| PA5117 | typA | 13 | 13 | 66.49199454 | 5.114768811 |
| PA5148 | PA5148 | 5 | 33 | 149.9935691 | 4.545259669 |
| PA5162 | rmlD | 11 | 6 | 40.20446181 | 6.700743636 |
| PA5198 | PA5198 | 9 | 4 | 12.37060363 | 3.092650909 |
| PA5260 | hemC | 8 | 0.1 | 12.37060363 | 123.7060363 |
| PA5285 | PA5285 | 1 | 8 | 57.21404181 | 7.151755226 |
| PA5366 | pstB | 11 | 1 | 6.185301817 | 6.185301817 |

| *Genes with higher insertion frequency + B.t. WT* |  |  |  |  |  |
| --- | --- | --- | --- | --- | --- |
| PA0006 | PA0006 | 13 | 156 | 34.01916 | 0.218071538 |
| PA0125 | PA0125 | 2 | 40 | 12.37060363 | 0.309265091 |
| PA0871 | phhB | 4 | 225 | 41.75078727 | 0.185559055 |
| PA1045 | PA1045 | 17 | 6 | 1.546325454 | 0.257720909 |
| PA1112a | PA1112a | 1 | 55 | 12.37060363 | 0.224920066 |
| PA1117 | PA1117 | 1 | 47 | 12.37060363 | 0.263204333 |
| PA1149 | PA1149 | 3 | 86 | 20.10223091 | 0.233746871 |
| PA1446 | fliP | 3 | 161 | 46.38976363 | 0.288135178 |
| PA1503 | PA1503 | 4 | 78 | 23.19488182 | 0.29737028 |
| PA1571 | PA1571 | 2 | 48 | 7.731627272 | 0.161075568 |
| PA1664 | PA1664 | 2 | 17 | 0.1 | 0.005882353 |
| PA1685 | masA | 10 | 30 | 9.277952726 | 0.309265091 |
| PA1742 | PA1742 | 12 | 129 | 32.47283454 | 0.2517274 |
| PA1743 | PA1743 | 3 | 42 | 6.185301817 | 0.147269091 |
| PA1802 | clpX | 20 | 19 | 6.185301817 | 0.325542201 |
| PA1842 | PA1842 | 2 | 72 | 23.19488182 | 0.322151136 |
| PA1995 | PA1995 | 3 | 33 | 10.82427818 | 0.32800843 |
| PA2003 | bdhA | 3 | 112 | 35.56548545 | 0.317548977 |
| PA2007 | maiA | 8 | 455 | 106.6964564 | 0.234497706 |
| PA2008 | fahA | 11 | 174 | 18.55590545 | 0.106643135 |
| PA2143 | PA2143 | 2 | 88 | 10.82427818 | 0.123003161 |
| PA2183 | PA2183 | 3 | 58 | 18.55590545 | 0.319929404 |
| PA2187 | PA2187 | 6 | 116 | 24.74120727 | 0.21328627 |
| PA2357 | msuE | 7 | 112 | 37.1118109 | 0.331355455 |
| PA2492 | mexT | 8 | 17782 | 4741.033843 | 0.266619831 |
| PA2614 | lolA | 5 | 63 | 20.10223091 | 0.31908303 |
| PA2967 | fabG | 8 | 11 | 1.546325454 | 0.140575041 |
| PA3139.1 | PA3139.1 | 3 | 17 | 0.1 | 0.005882353 |
| PA3165 | hisC2 | 15 | 32 | 0.1 | 0.003125 |
| PA3368.1 | PA3368.1 | 3 | 35 | 6.185301817 | 0.176722909 |
| PA3520 | PA3520 | 4 | 56 | 13.91692909 | 0.248516591 |
| PA3557 | arnE | 1 | 18 | 3.092650909 | 0.171813939 |
| PA3721 | nalC | 8 | 6 | 1.546325454 | 0.257720909 |
| PA3808 | PA3808 | 2 | 46 | 12.37060363 | 0.268926166 |
| PA3813 | iscU | 3 | 39 | 1.546325454 | 0.039649371 |
| PA3821 | secD | 16 | 10 | 0.1 | 0.01 |
| PA3822 | PA3822 | 3 | 12 | 0.1 | 0.008333333 |
| PA4092 | hpaC | 1 | 38 | 6.185301817 | 0.1627711 |
| PA4236 | katA | 23 | 2815 | 896.8687635 | 0.318603468 |
| PA4561 | ribF | 7 | 27 | 4.638976363 | 0.171813939 |
| PA4662 | murI | 15 | 30 | 3.092650909 | 0.103088364 |
| PA4758 | carA | 17 | 5 | 1.546325454 | 0.309265091 |
| PA5062 | PA5062 | 0 | 21 | 3.092650909 | 0.147269091 |
| PA5134 | PA5134 | 11 | 319 | 94.32585272 | 0.295692328 |

**Supplemental Table 3.** Transposon sequencing-based analysis of *P. aeruginosa* fitness determinants during antagonism by *B. thai*.

|  |  |  | Normalized insertion counts |  |  |
| --- | --- | --- | --- | --- | --- |
| Locus ID | Gene name | TA sites | + *B.t.* WT | + *B.t.* $\Delta colA$ | Fold change ($\Delta colA$/WT) |
| ***Genes with higher insertion frequency + *B.t.* $\Delta colA$*** |  |  |  |  |  |
| PA0005 | lptA | 11 | 2 | 7.928575216 | 3.964287608 |
| PA0037 | trpI | 8 | 237 | 767.4860809 | 3.238337894 |
| PA0180 | cttP | 5 | 3 | 17.44286548 | 5.814288492 |
| PA0197 | tonB2 | 2 | 5 | 25.37144069 | 5.074288138 |
| PA0407 | gshB | 10 | 7 | 25.37144069 | 3.624491527 |
| PA0466 | PA0466 | 1 | 0 | 11.1000053 | N.D. |
| PA0526 | PA0526 | 1 | 38 | 185.5286601 | 4.88233316 |
| PA0712 | PA0712 | 4 | 7 | 33.30001591 | 4.75714513 |
| PA0753 | PA0753 | 5 | 15 | 63.42860173 | 4.228573449 |
| PA0928 | gacS | 18 | 3 | 9.51429026 | 3.171430087 |
| PA0947 | PA0947 | 7 | 7 | 22.20001061 | 3.171430087 |
| PA0960 | PA0960 | 0 | 10 | 44.40002121 | 4.440002121 |
| PA0973 | oprL | 11 | 15 | 57.08574156 | 3.805716104 |
| PA1034 | PA1034 | 2 | 6 | 49.15716634 | 8.192861057 |
| PA1045 | PA1045 | 17 | 6 | 30.12858582 | 5.02143097 |
| PA1121 | yfiR | 7 | 21 | 77.70003712 | 3.700001768 |
| PA1233 | PA1233 | 5 | 16 | 49.15716634 | 3.072322896 |
| PA1394 | PA1394 | 0 | 11 | 33.30001591 | 3.027274174 |
| PA1548 | PA1548 | 3 | 0 | 14.27143539 | N.D. |
| PA1550 | PA1550 | 6 | 3 | 9.51429026 | 3.171430087 |
| PA1552.1 | ccoQ1 | 2 | 0 | 26.95715574 | N.D. |
| PA1564 | PA1564 | 2 | 0 | 12.68572035 | N.D. |
| PA1645 | PA1645 | 5 | 9 | 36.471446 | 4.052382888 |
| PA1696 | pscO | 3 | 5 | 42.81430617 | 8.562861234 |
| PA1747 | PA1747 | 1 | 0 | 15.85715043 | N.D. |
| PA1796.2 | PA1796.2 | 2 | 0 | 12.68572035 | N.D. |
| PA1816 | dnaQ | 8 | 0 | 15.85715043 | N.D. |
| PA1879 | PA1879 | 2 | 2 | 19.02858052 | 9.51429026 |
| PA2176 | PA2176 | 4 | 17 | 52.32859643 | 3.078152731 |
| PA2193 | hcnA | 2 | 1 | 20.61429556 | 20.61429556 |
| PA2321 | PA2321 | 3 | 1 | 26.95715574 | 26.95715574 |
| PA2330 | PA2330 | 3 | 2 | 11.1000053 | 5.550002651 |
| PA2425 | pvdG | 13 | 8 | 39.64287608 | 4.95535951 |
| PA2541 | PA2541 | 5 | 9 | 30.12858582 | 3.347620647 |
| PA2554 | PA2554 | 3 | 11 | 47.5714513 | 4.324677391 |
| PA2724 | PA2724 | 4 | 41 | 161.7429344 | 3.94494962 |
| PA2738 | himA | 3 | 4 | 12.68572035 | 3.171430087 |
| PA2743 | infC | 12 | 2 | 6.342860173 | 3.171430087 |
| PA2805 | PA2805 | 0 | 6 | 20.61429556 | 3.435715927 |
| PA2855 | PA2855 | 3 | 15 | 45.98573625 | 3.06571575 |
| PA2895 | PA2895 | 1 | 9 | 28.54287078 | 3.171430087 |
| PA2898 | PA2898 | 1 | 4 | 15.85715043 | 3.964287608 |
| PA2914 | PA2914 | 5 | 33 | 115.7571982 | 3.507793884 |
| PA2947 | PA2947 | 2 | 18 | 63.42860173 | 3.523811207 |
| PA2964 | pabC | 5 | 5 | 19.02858052 | 3.805716104 |
| PA2991 | sth | 18 | 0 | 14.27143539 | N.D. |
| PA2992 | PA2992 | 2 | 25 | 98.31433268 | 3.932573307 |
| PA3051 | PA3051 | 2 | 5 | 17.44286548 | 3.488573095 |
| PA3067 | PA3067 | 1 | 0 | 25.37144069 | N.D. |
| PA3094.2 | PA3094.2 | 3 | 2 | 23.78572565 | 11.89286282 |
| PA3099 | xcpV | 1 | 8 | 34.88573095 | 4.360716369 |
| PA3111 | folC | 15 | 5 | 42.81430617 | 8.562861234 |
| PA3133.4 | PA3133.4 | 3 | 0 | 14.27143539 | N.D. |
| PA3145 | wbpL | 34 | 1 | 9.51429026 | 9.51429026 |
| PA3205 | PA3205 | 1 | 5 | 20.61429556 | 4.122859113 |
| PA3382 | phnE | 4 | 32 | 115.7571982 | 3.617412442 |
| PA3385 | amrZ | 3 | 20 | 61.84288669 | 3.092144334 |
| PA3574a | PA3574a | 3 | 11 | 53.91431147 | 4.901301043 |
| PA3634 | PA3634 | 2 | 4 | 14.27143539 | 3.567858847 |
| PA3640 | dnaE | 46 | 2 | 15.85715043 | 7.928575216 |
| PA3649 | PA3649 | 20 | 22 | 74.52860703 | 3.387663956 |
| PA3815 | iscR | 7 | 13 | 49.15716634 | 3.781320488 |
| PA3879 | narL | 2 | 31 | 123.6857734 | 3.989863657 |
| PA4020 | mpl | 15 | 1 | 3.171430087 | 3.171430087 |
| PA4275 | nusG | 9 | 8 | 71.35717695 | 8.919647118 |
| PA4277 | tufB | 23 | 2 | 6.342860173 | 3.171430087 |
| PA4319 | PA4319 | 13 | 95 | 307.6287184 | 3.238197036 |
| PA4320 | PA4320 | 19 | 70 | 274.3287025 | 3.918981464 |
| PA4322 | PA4322 | 6 | 22 | 115.7571982 | 5.261690825 |
| PA4323 | PA4323 | 15 | 27 | 107.8286229 | 3.993652702 |
| PA4351 | PA4351 | 4 | 9 | 47.5714513 | 5.285716811 |
| PA4413 | ftsW | 12 | 9 | 50.74288138 | 5.638097932 |
| PA4422 | PA4422 | 6 | 0 | 14.27143539 | N.D. |
| PA4431 | PA4431 | 8 | 2 | 6.342860173 | 3.171430087 |
| PA4452 | PA4452 | 2 | 11 | 49.15716634 | 4.468833304 |
| PA4459 | PA4459 | 13 | 9 | 31.71430087 | 3.523811207 |
| PA4610 | PA4610 | 1 | 17 | 53.91431147 | 3.171430087 |
| PA4671 | PA4671 | 7 | 2 | 11.1000053 | 5.550002651 |
| PA4672 | PA4672 | 5 | 3 | 12.68572035 | 4.228573449 |
| PA4726.1 | PA4726.1 | 4 | 1 | 17.44286548 | 17.44286548 |
| PA4820 | PA4820 | 5 | 9 | 26.95715574 | 2.995239526 |
| PA4853 | fis | 4 | 1 | 19.02858052 | 19.02858052 |
| PA4890 | desT | 2 | 0 | 20.61429556 | N.D. |
| PA4894 | PA4894 | 3 | 8 | 52.32859643 | 6.541074553 |
| PA4933 | PA4933 | 10 | 7 | 22.20001061 | 3.171430087 |
| PA4952 | PA4952 | 5 | 4 | 26.95715574 | 6.739288934 |
| PA5000 | wapR | 21 | 6 | 25.37144069 | 4.228573449 |
| PA5016 | aceF | 7 | 3 | 22.20001061 | 7.400003535 |
| PA5049 | rpmE | 7 | 4 | 22.20001061 | 5.550002651 |
| PA5148 | PA5148 | 5 | 33 | 111.000053 | 3.363637971 |
| PA5162 | rmlD | 11 | 6 | 45.98573625 | 7.664289376 |
| PA5198 | PA5198 | 9 | 4 | 17.44286548 | 4.360716369 |
| PA5199 | amgS | 9 | 3 | 23.78572565 | 7.928575216 |
| PA5285 | PA5285 | 1 | 8 | 34.88573095 | 4.360716369 |

| *Genes with higher insertion frequency + B.t. WT* |  |  |  |  |  |
| --- | --- | --- | --- | --- | --- |
| PA0006 | PA0006 | 13 | 156 | 41.22859113 | 0.264285841 |
| PA0125 | PA0125 | 2 | 40 | 7.928575216 | 0.19821438 |
| PA0420 | bioA | 19 | 45 | 12.68572035 | 0.281904897 |
| PA0666 | PA0666 | 10 | 127 | 36.471446 | 0.28717674 |
| PA0715 | PA0715 | 61 | 14 | 3.171430087 | 0.22653072 |
| PA0763 | mucA | 9 | 52 | 15.85715043 | 0.304945201 |
| PA0905.3 | PA0905.3 | 4 | 10 | 1.585715043 | 0.158571504 |
| PA0913 | mgtE | 10 | 96 | 23.78572565 | 0.247767976 |
| PA1030.1 | PA1030.1 | 2 | 55 | 15.85715043 | 0.288311826 |
| PA1117 | PA1117 | 1 | 47 | 1.585715043 | 0.033738618 |
| PA1149 | PA1149 | 3 | 86 | 12.68572035 | 0.147508376 |
| PA1159 | PA1159 | 3 | 32 | 9.51429026 | 0.297321571 |
| PA1801 | clpP | 12 | 22 | 6.342860173 | 0.288311826 |
| PA1845 | PA1845 | 4 | 184 | 58.6714566 | 0.318866612 |
| PA1985 | pqqA | 1 | 12 | 0.1 | 0.008333333 |
| PA2142a | PA2142a | 2 | 50 | 9.51429026 | 0.190285805 |
| PA2187 | PA2187 | 6 | 116 | 36.471446 | 0.314409017 |
| PA2375 | PA2375 | 1 | 32 | 9.51429026 | 0.297321571 |
| PA2421 | PA2421 | 16 | 302 | 99.90004773 | 0.33079486 |
| PA2645 | nuoJ | 8 | 145 | 45.98573625 | 0.317143009 |
| PA2778 | PA2778 | 7 | 325 | 104.6571929 | 0.322022132 |
| PA2967 | fabG | 8 | 11 | 0.1 | 0.009090909 |
| PA2970 | rpmF | 5 | 30 | 7.928575216 | 0.264285841 |
| PA3014 | faoA | 19 | 17 | 0.1 | 0.005882353 |
| PA3139.1 | PA3139.1 | 3 | 17 | 4.75714513 | 0.279832066 |
| PA3151 | hisF2 | 27 | 19 | 3.171430087 | 0.166917373 |
| PA3165 | hisC2 | 15 | 32 | 9.51429026 | 0.297321571 |
| PA3482 | metG | 25 | 71 | 20.61429556 | 0.290342191 |
| PA3496 | PA3496 | 1 | 71 | 12.68572035 | 0.178672118 |
| PA3808 | PA3808 | 2 | 46 | 6.342860173 | 0.137888265 |
| PA3821 | secD | 16 | 10 | 3.171430087 | 0.317143009 |
| PA3844 | PA3844 | 8 | 898 | 247.3715468 | 0.275469428 |
| PA4092 | hpaC | 1 | 38 | 0 | 0 |
| PA4236 | katA | 23 | 2815 | 829.3289676 | 0.294610646 |
| PA4479 | mreD | 6 | 56 | 17.44286548 | 0.311479741 |
| PA4581.1 | PA4581.1 | 2 | 21 | 6.342860173 | 0.302040961 |
| PA4663 | moeB | 9 | 28 | 3.171430087 | 0.11326536 |
| PA5114 | PA5114 | 37 | 218 | 55.50002651 | 0.254587278 |
| PA5134 | PA5134 | 11 | 319 | 71.35717695 | 0.22369021 |
| PA5223 | ubiH | 14 | 36 | 7.928575216 | 0.2202382 |
| PA5241 | ppx | 15 | 10 | 3.171430087 | 0.317143009 |

**Supplemental Table 4.**
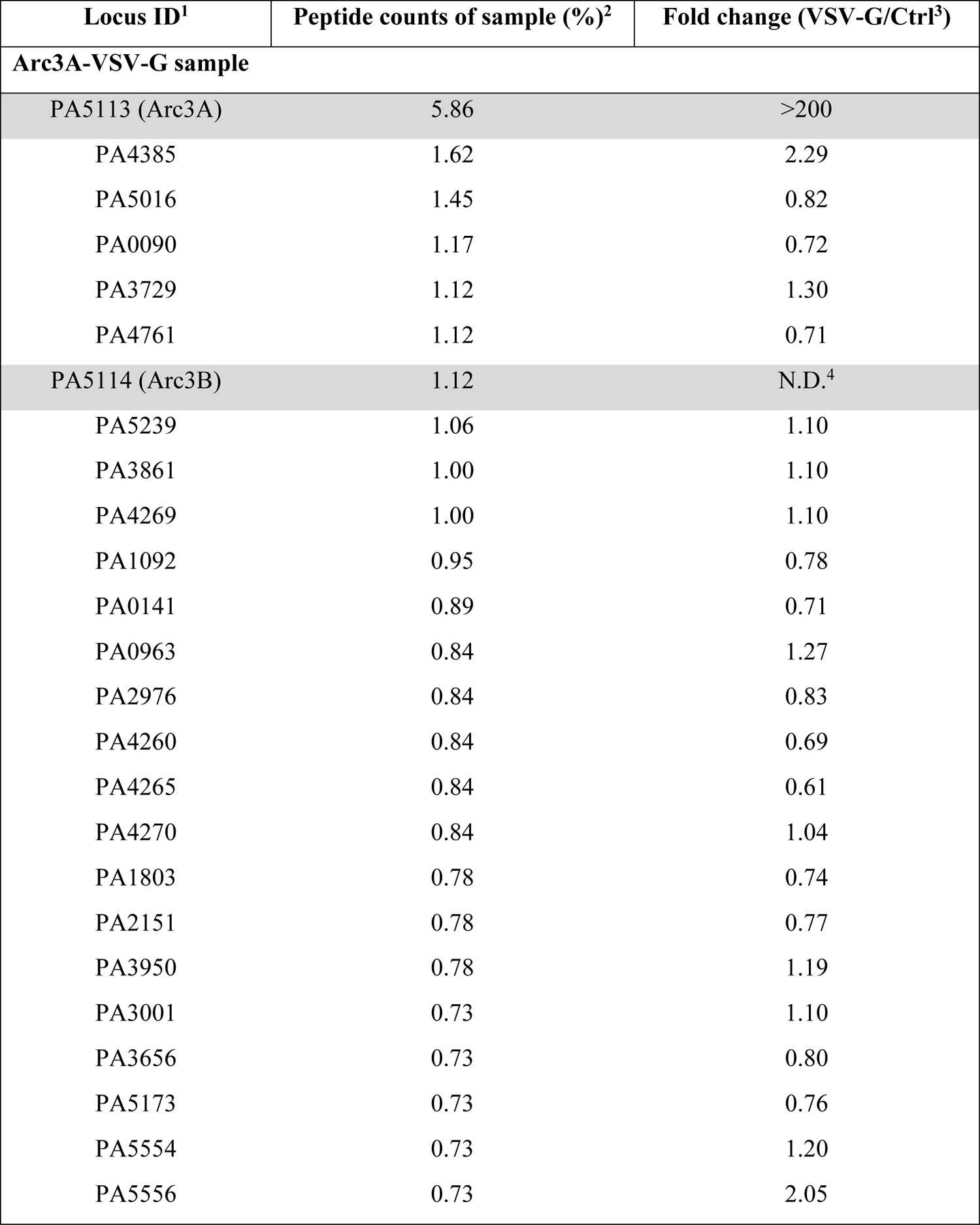
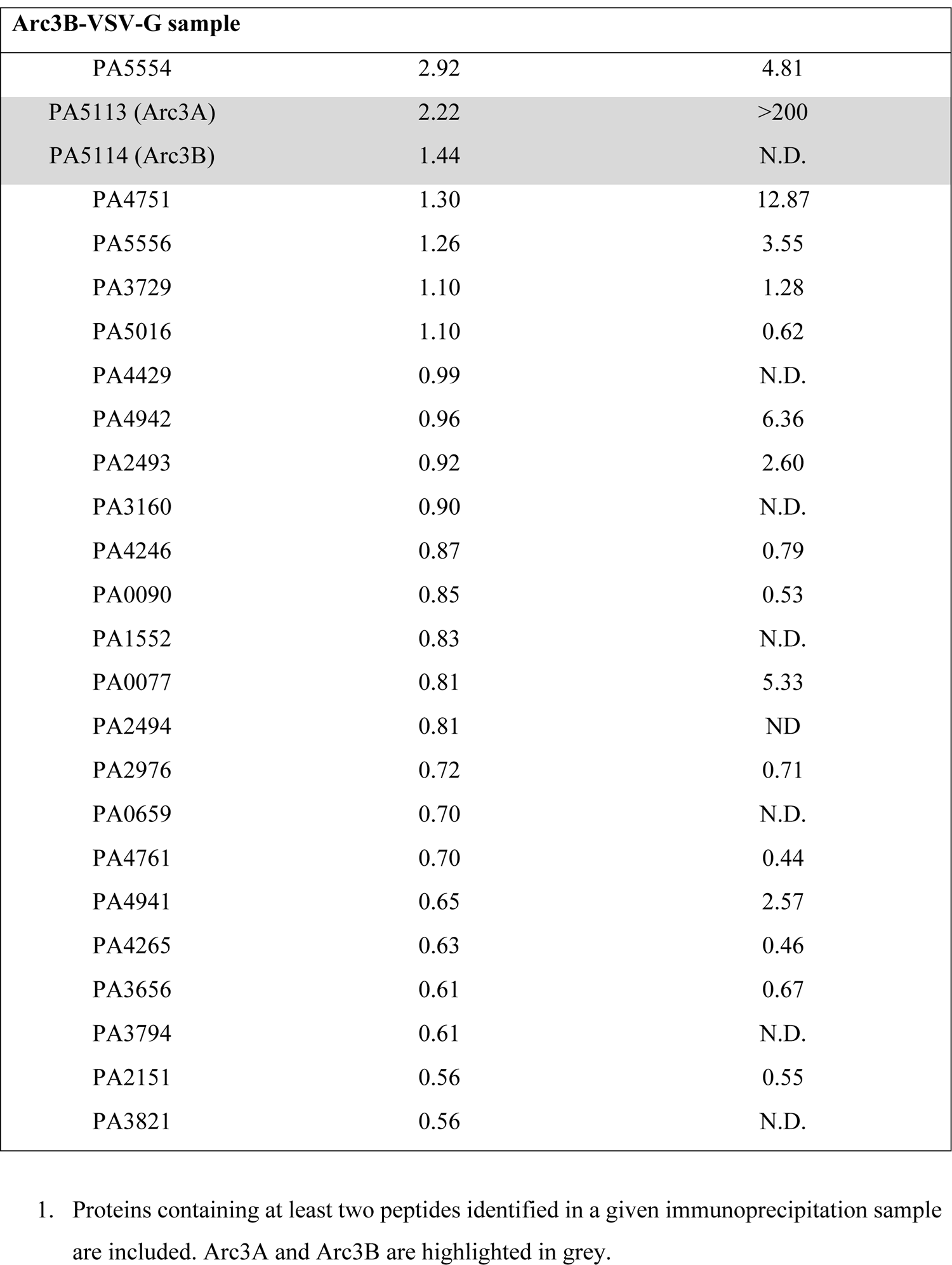
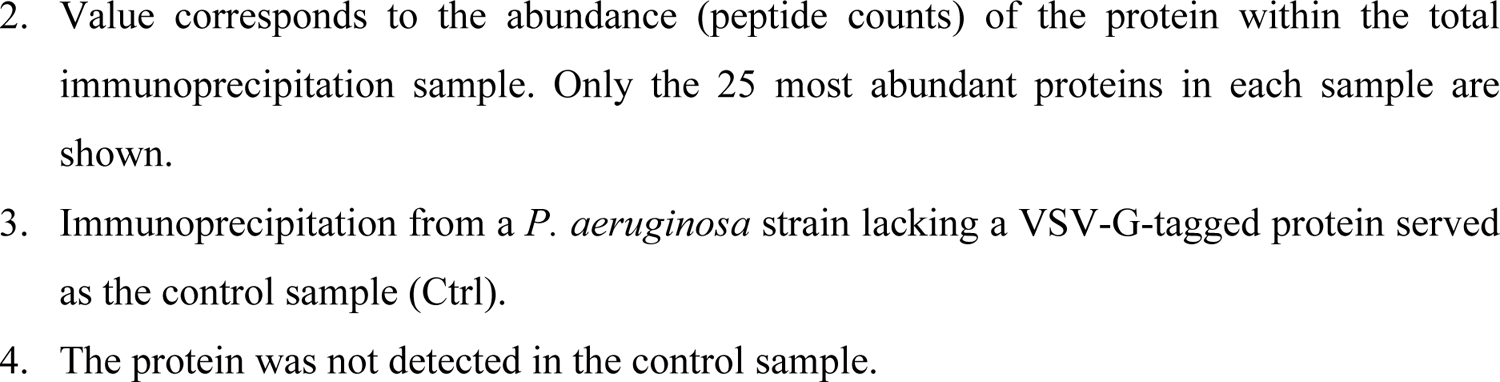
Mass spectrometric analysis of *P. aeruginosa* Arc3A and Arc3B immunoprecipitation samples.

**Supplemental Table 5.**
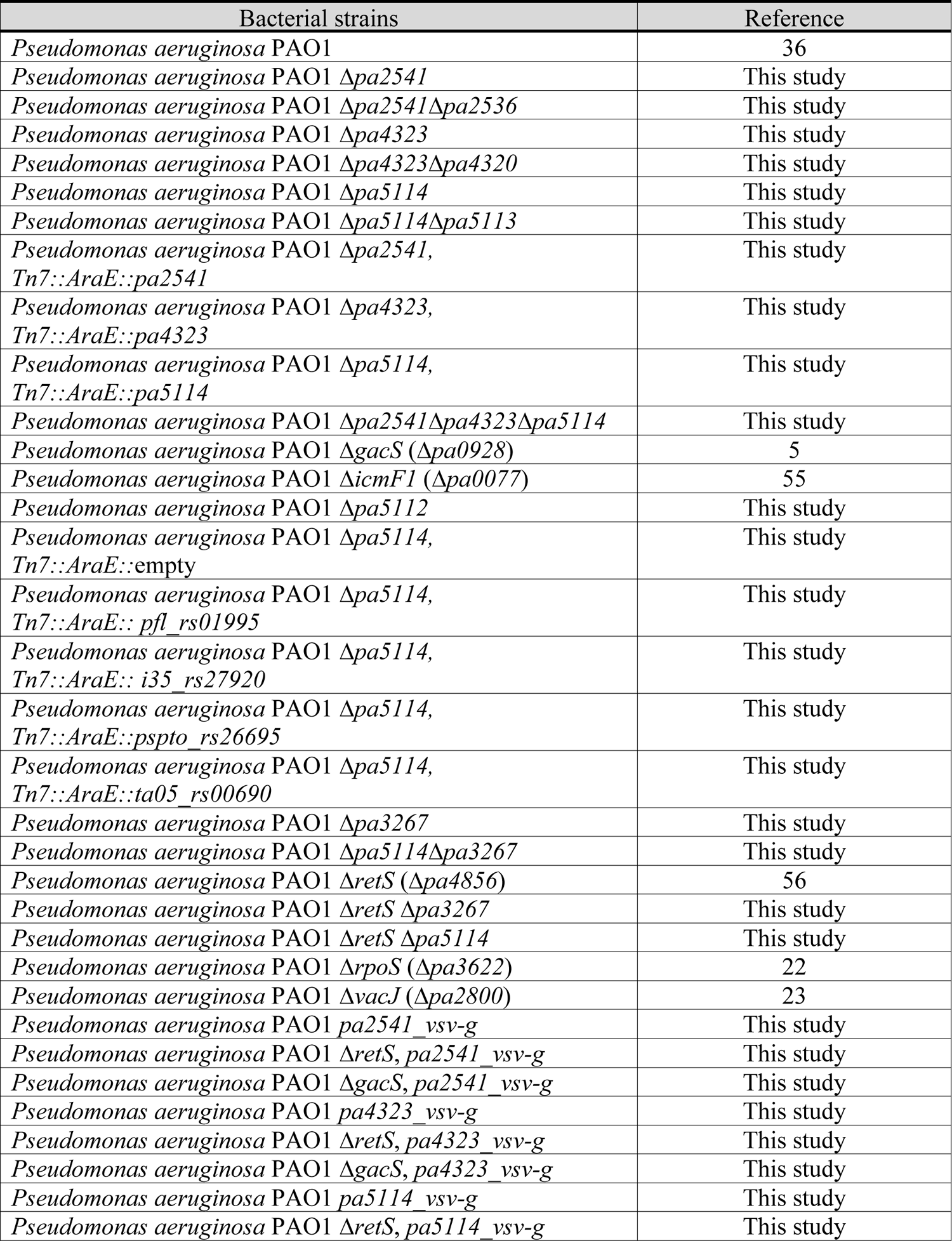
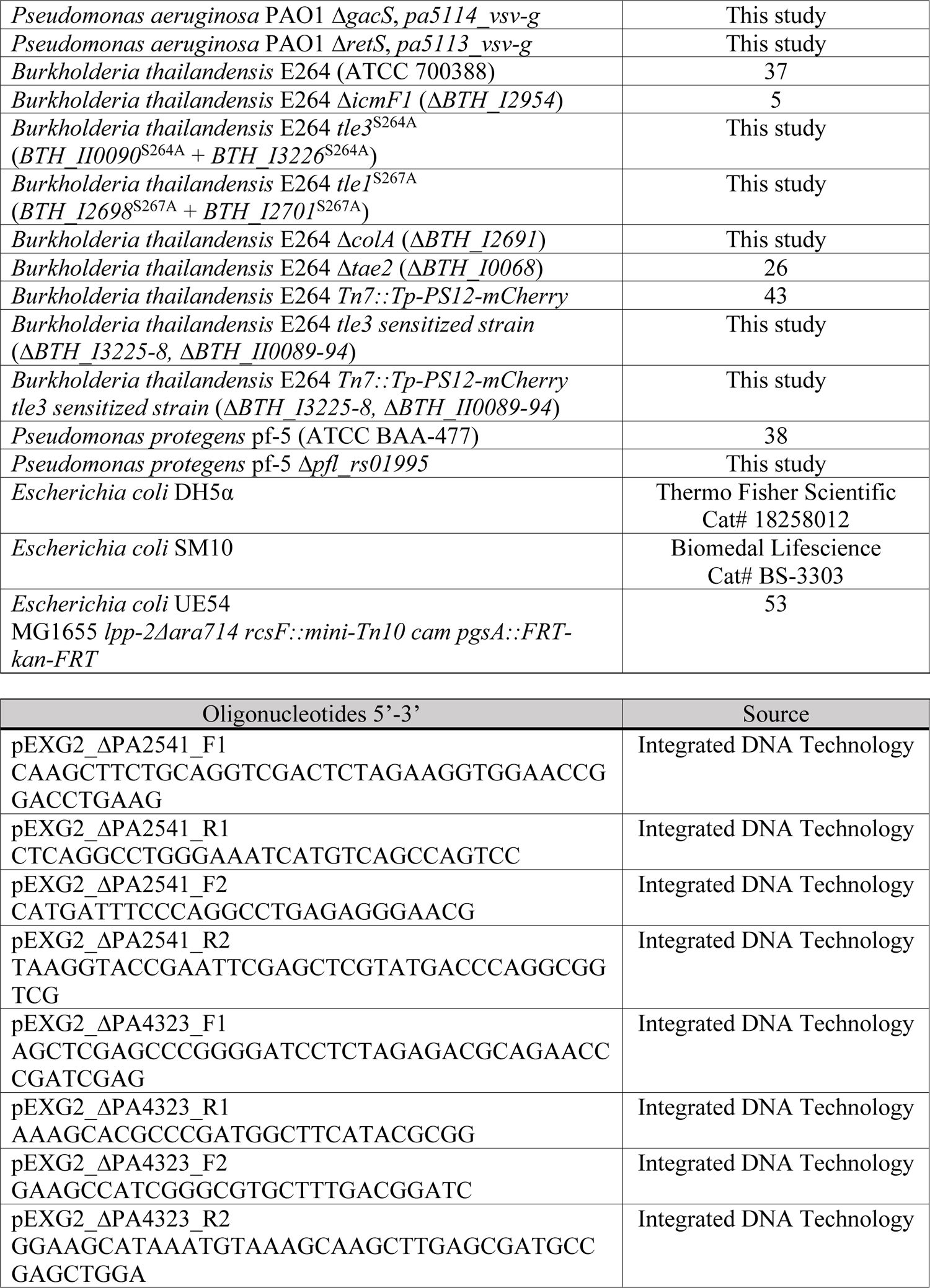
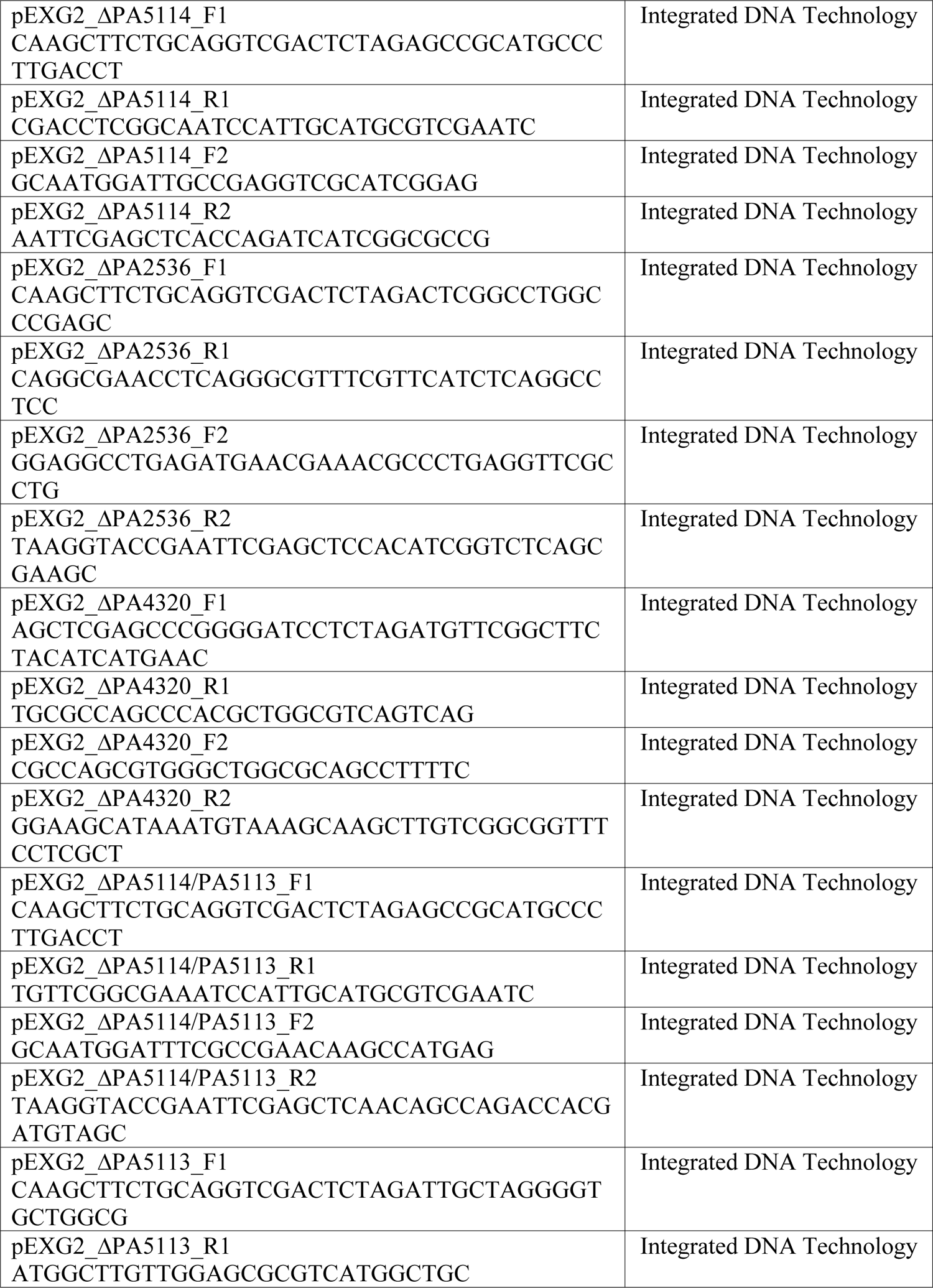
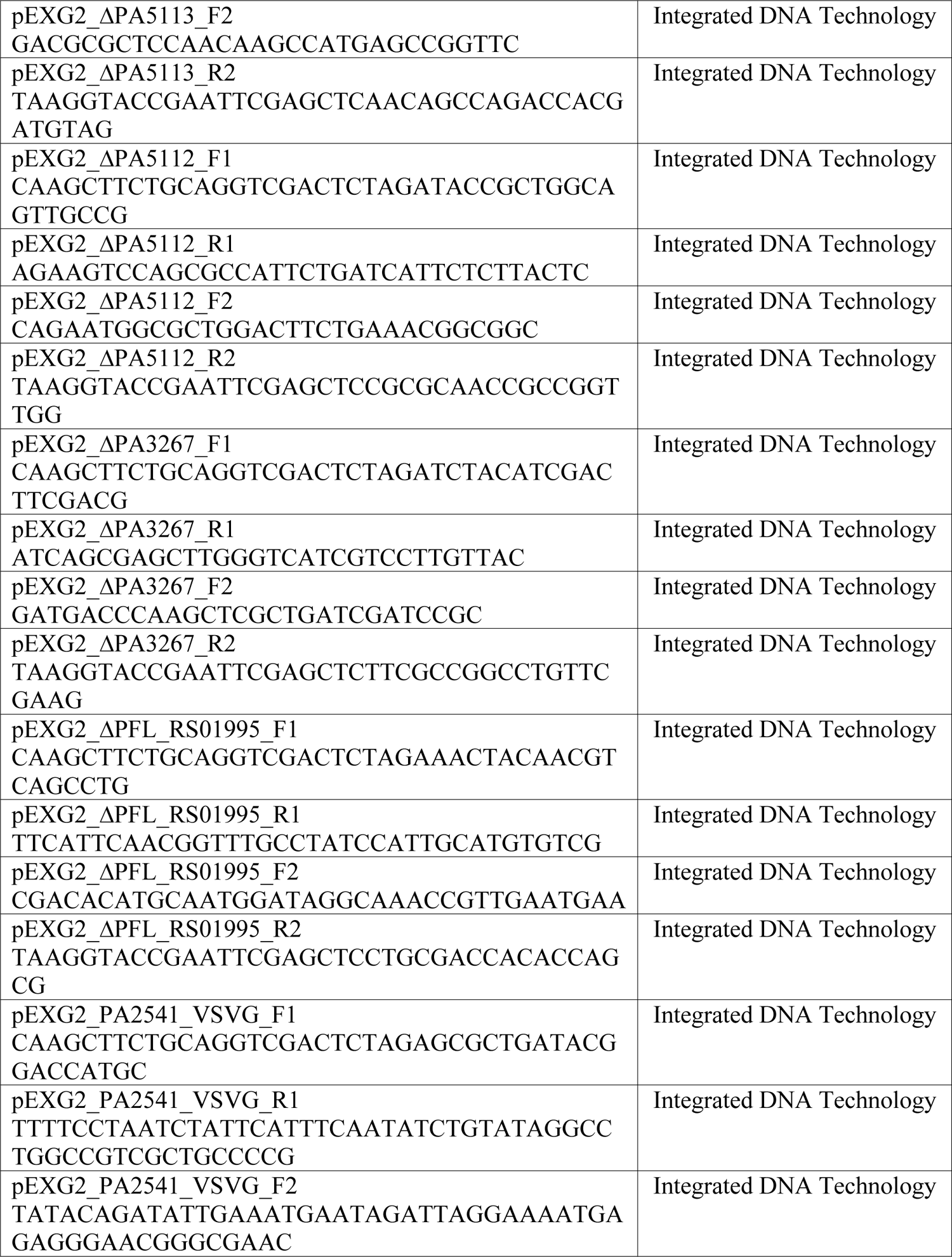
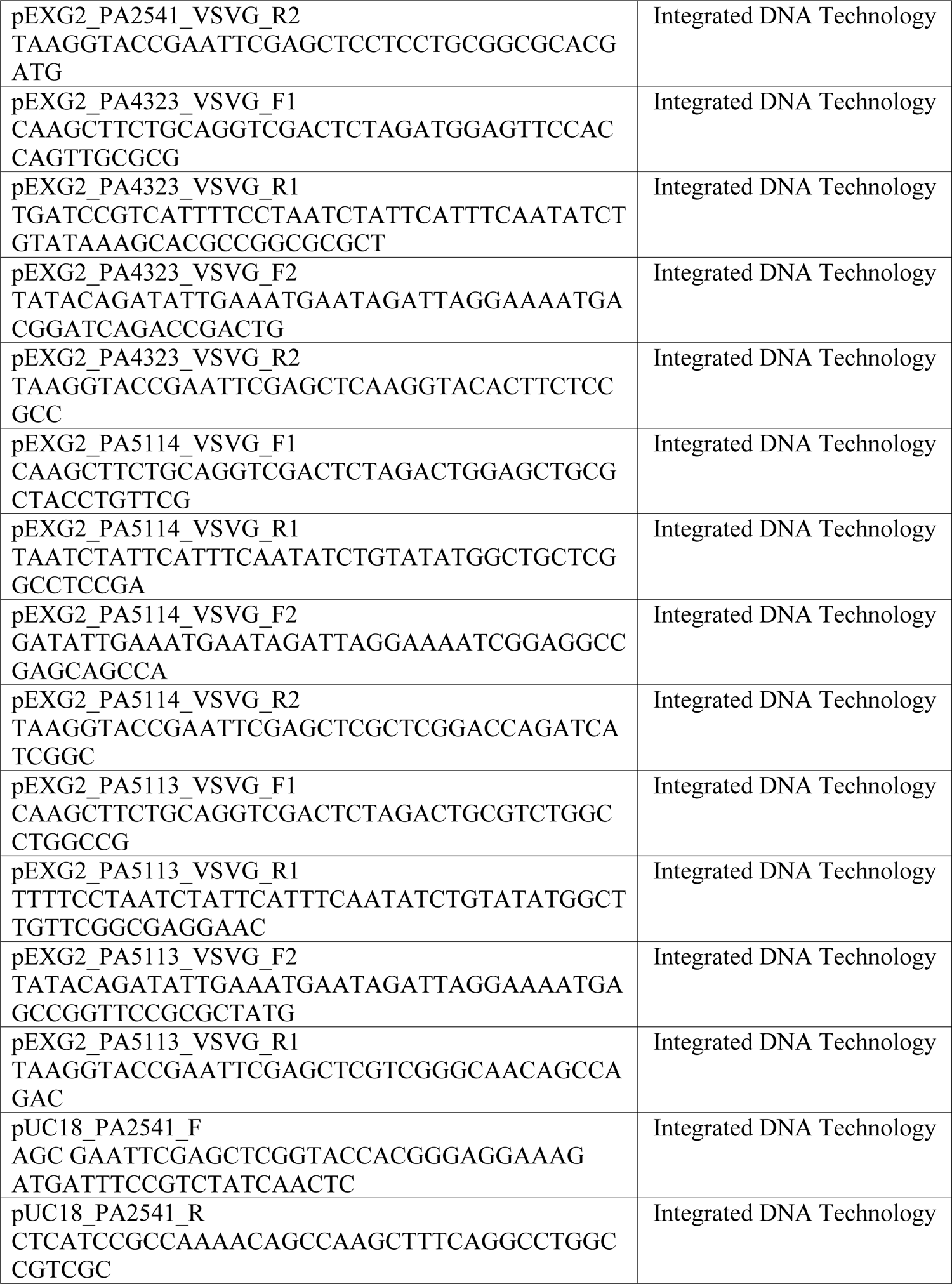
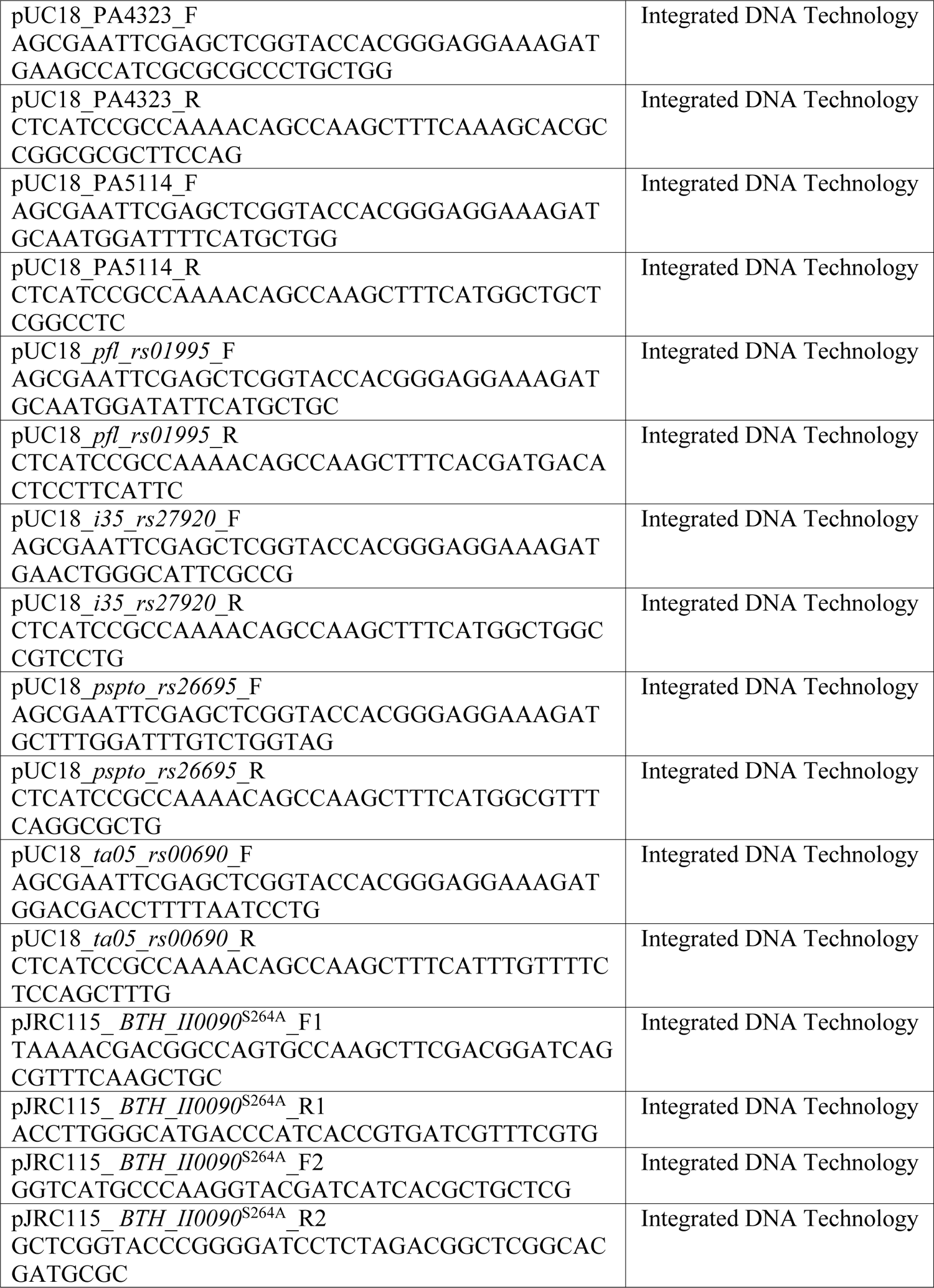
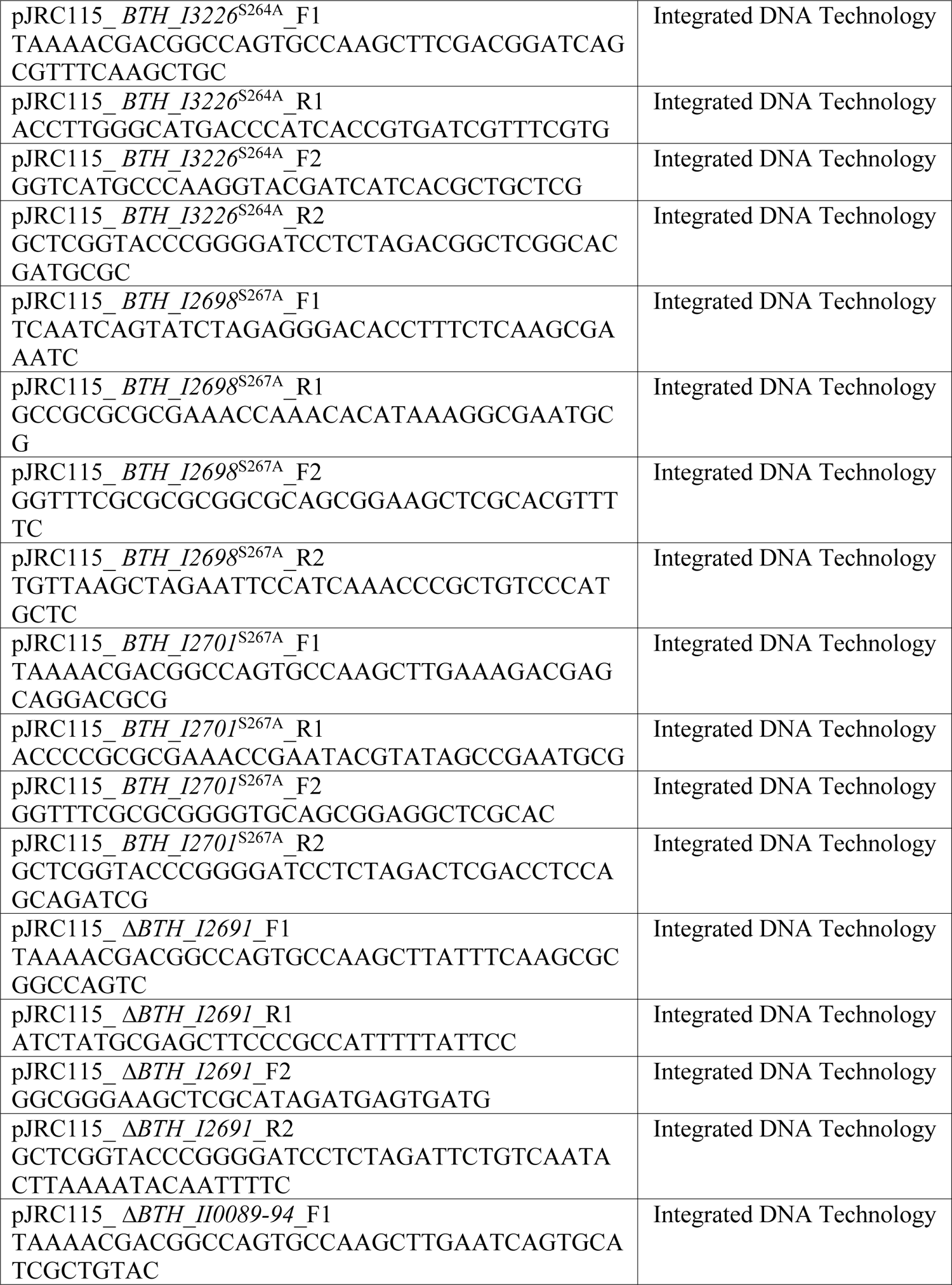
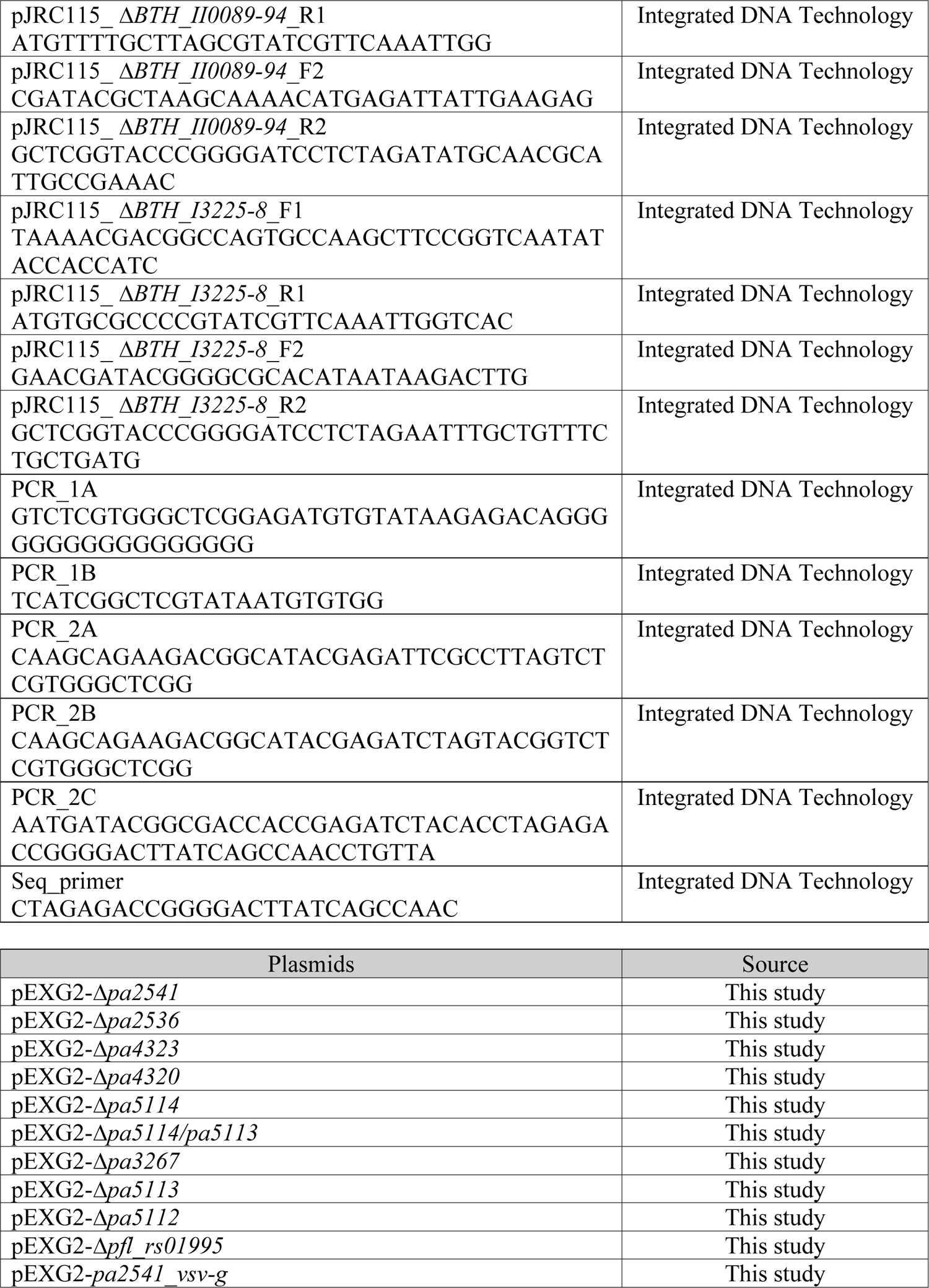
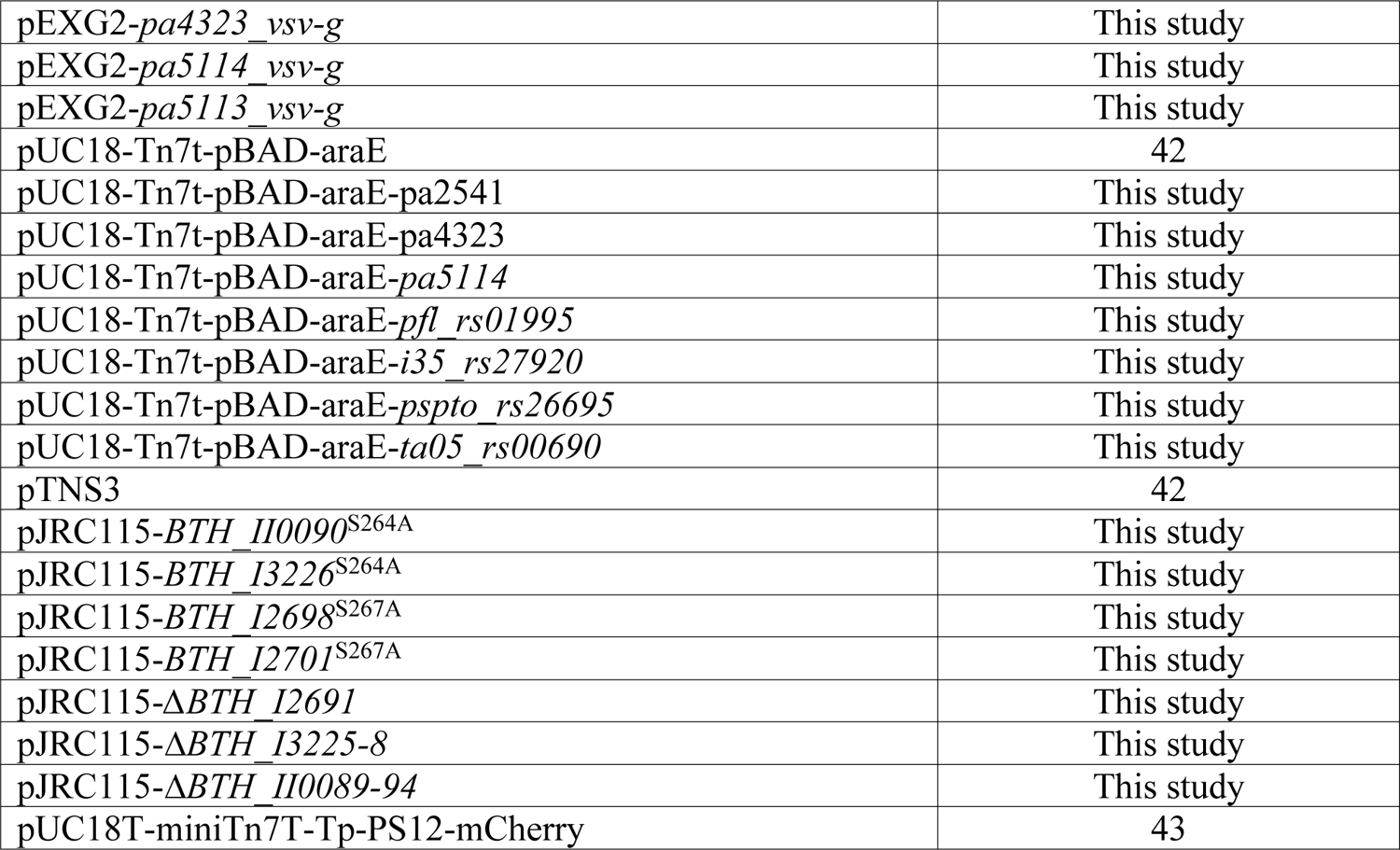
Bacterial strains, Oligonucleotides, Plasmids used in the study.

**Supplemental Table 6.**
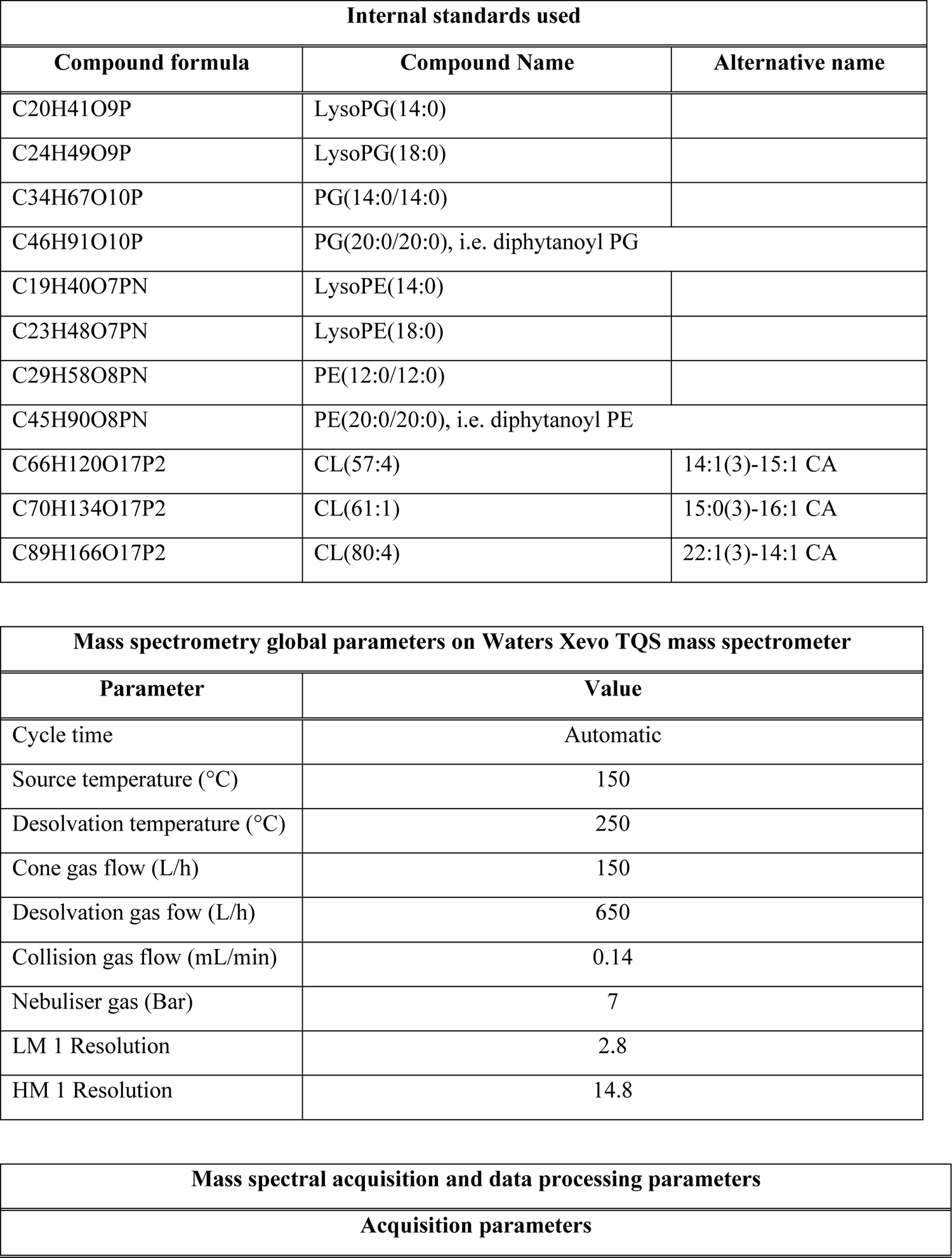
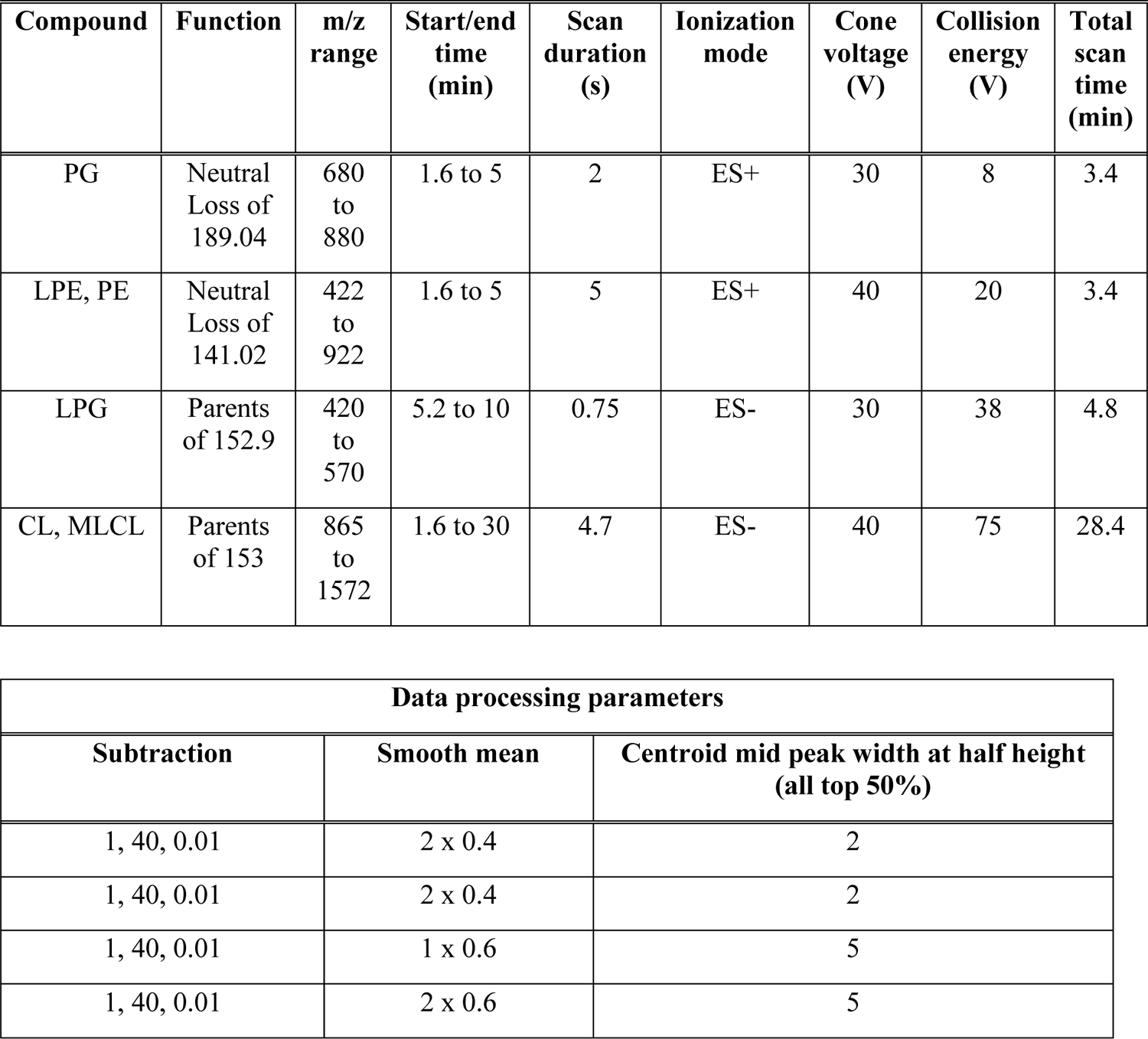
Internal standards and parameters used in lipidomic analysis

| Internal standards used |  |  |
| --- | --- | --- |
| Compound formula | Compound Name | Alternative name |
| C20H41O9P | LysoPG(14:0) |  |
| C24H49O9P | LysoPG(18:0) |  |
| C34H67O10P | PG(14:0/14:0) |  |
| C46H91O10P | PG(20:0/20:0), i.e. diphytanoyl PG |  |
| C19H40O7PN | LysoPE(14:0) |  |
| C23H48O7PN | LysoPE(18:0) |  |
| C29H58O8PN | PE(12:0/12:0) |  |
| C45H90O8PN | PE(20:0/20:0), i.e. diphytanoyl PE |  |
| C66H120O17P2 | CL(57:4) | 14:1(3)-15:1 CA |
| C70H134O17P2 | CL(61:1) | 15:0(3)-16:1 CA |
| C89H166O17P2 | CL(80:4) | 22:1(3)-14:1 CA |

| Mass spectrometry global parameters on Waters Xevo TQS mass spectrometer |  |
| --- | --- |
| Parameter | Value |
| Cycle time | Automatic |
| Source temperature (°C) | 150 |
| Desolvation temperature (°C) | 250 |
| Cone gas flow (L/h) | 150 |
| Desolvation gas flow (L/h) | 650 |
| Collision gas flow (mL/min) | 0.14 |
| Nebuliser gas (Bar) | 7 |
| LM 1 Resolution | 2.8 |
| HM 1 Resolution | 14.8 |

| Mass spectral acquisition and data processing parameters |
| --- |
| Acquisition parameters |

| Compound | Function | m/z range | Start/end time (min) | Scan duration (s) | Ionization mode | Cone voltage (V) | Collision energy (V) | Total scan time (min) |
| --- | --- | --- | --- | --- | --- | --- | --- | --- |
| PG | Neutral Loss of 189.04 | 680 to 880 | 1.6 to 5 | 2 | ES+ | 30 | 8 | 3.4 |
| LPE, PE | Neutral Loss of 141.02 | 422 to 922 | 1.6 to 5 | 5 | ES+ | 40 | 20 | 3.4 |
| LPG | Parents of 152.9 | 420 to 570 | 5.2 to 10 | 0.75 | ES- | 30 | 38 | 4.8 |
| CL, MLCL | Parents of 153 | 865 to 1572 | 1.6 to 30 | 4.7 | ES- | 40 | 75 | 28.4 |

| Data processing parameters |  |  |
| --- | --- | --- |
| Subtraction | Smooth mean | Centroid mid peak width at half height (all top 50%) |
| 1, 40, 0.01 | 2 x 0.4 | 2 |
| 1, 40, 0.01 | 2 x 0.4 | 2 |
| 1, 40, 0.01 | 1 x 0.6 | 5 |
| 1, 40, 0.01 | 2 x 0.6 | 5 |

**Supplemental Table 7.** Transposon sequencing data summary

| **Growth experiment** | **Mutant pool pre-competition** | **Pool with WT *B. thai* - rep 1** | **Pool with ΔT6S *B. thai* - rep 1** |
| --- | --- | --- | --- |
| **fastq file name** | tn_mutant_pool.fastq | pool_v_wt_rep1.fastq | pool_v_deltaT6_rep1.fastq |
| **Total reads** | 515,434 | 5,788,728 | 6,452,899 |
| **Trimmed (valid Tn prefix)** | 515,293 | 5,786,427 | 6,450,594 |
| **Mapped** | 350,923 | 4,666,597 | 5,283,645 |
| **Mapped to TA sites** | 349,915 | 4,634,942 | 5,241,672 |
| **TA_sites** | 94,404 | 94,404 | 94,404 |
| **TAs_hit** | 50,348 | 57,644 | 60,331 |
| **Max reads per TA site** | 324 | 9,519 | 5,886 |
| **Mean reads per hit TA site** | 6.9 | 80.4 | 86.9 |

| **Pool with WT *B.thai* - rep 2** | **Pool with ΔT6S *B.thai* - rep 2** | **Pool with *tle3*<sup>S264A</sup> *B. thai*** | **Pool with ΔcolA *B. thai*** |
| --- | --- | --- | --- |
| pool_v_wt_rep2.fastq | pool_v_deltaT6_rep2.fastq | pool_v_tle3.fastq | pool_v_colA.fastq |
| 411,515 | 720,133 | 3,226,978 | 3,130,545 |
| 411,401 | 719,895 | 3,225,816 | 3,129,574 |
| 266,466 | 481,466 | 3,024,047 | 2,938,318 |
| 265,661 | 479,769 | 2,997,466 | 2,922,995 |
| 94,404 | 94,404 | 94,404 | 94,404 |
| 46,500 | 53,148 | 59,145 | 58,101 |
| 669 | 447 | 3,484 | 3,568 |
| 5.7 | 9.0 | 50.7 | 50.3 |

## References

1. Peterson, S. B., Bertolli, S. K. & Mougous, J. D. The Central Role of Interbacterial Antagonism in Bacterial Life. Curr Biol 30, R1203–R1214, doi:10.1016/j.cub.2020.06.103 (2020).

2. Granato, E. T., Meiller-Legrand, T. A. & Foster, K. R. The Evolution and Ecology of Bacterial Warfare. Curr Biol 29, R521–R537, doi:10.1016/j.cub.2019.04.024 (2019).

3. Hampton, H. G., Watson, B. N. J. & Fineran, P. C. The arms race between bacteria and their phage foes. Nature 577, 327–336, doi:10.1038/s41586-019-1894-8 (2020).

4. Bernheim, A. & Sorek, R. The pan-immune system of bacteria: antiviral defence as a community resource. Nat Rev Microbiol 18, 113–119, doi:10.1038/s41579-019-0278-2 (2020).

5. LeRoux, M. et al. Kin cell lysis is a danger signal that activates antibacterial pathways of Pseudomonas aeruginosa. Elife 4, doi:10.7554/eLife.05701 (2015).

6. LeRoux, M., Peterson, S. B. & Mougous, J. D. Bacterial danger sensing. J Mol Biol 427, 3744–3753, doi:10.1016/j.jmb.2015.09.018 (2015).

7. Robitaille, S., Trus, E. & Ross, B. D. Bacterial Defense against the Type VI Secretion System. Trends Microbiol 29, 187–190, doi:10.1016/j.tim.2020.09.001 (2021).

8. Hersch, S. J., Manera, K. & Dong, T. G. Defending against the Type Six Secretion System: beyond Immunity Genes. Cell Rep 33, 108259, doi:10.1016/j.celrep.2020.108259 (2020).

9. Ross, B. D. et al. Human gut bacteria contain acquired interbacterial defence systems. Nature 575, 224–228, doi:10.1038/s41586-019-1708-z (2019).

10. Hersch, S. J. et al. Envelope stress responses defend against type six secretion system attacks independently of immunity proteins. Nat Microbiol 5, 706–714, doi:10.1038/s41564-020-0672-6 (2020).

11. Toska, J., Ho, B. T. & Mekalanos, J. J. Exopolysaccharide protects Vibrio cholerae from exogenous attacks by the type 6 secretion system. Proc Natl Acad Sci U S A 115, 7997– 8002, doi:10.1073/pnas.1808469115 (2018).

12. Le, N. H. et al. Peptidoglycan editing provides immunity to Acinetobacter baumannii during bacterial warfare. Sci Adv 6, eabb5614, doi:10.1126/sciadv.abb5614 (2020).

13. Lories, B. et al. Biofilm Bacteria Use Stress Responses to Detect and Respond to Competitors. Curr Biol 30, 1231–1244 e1234, doi:10.1016/j.cub.2020.01.065 (2020).

14. Cornforth, D. M. & Foster, K. R. Competition sensing: the social side of bacterial stress responses. Nat Rev Microbiol 11, 285–293, doi:10.1038/nrmicro2977 (2013).

15. Kamal, F. et al. Differential Cellular Response to Translocated Toxic Effectors and Physical Penetration by the Type VI Secretion System. Cell Rep 31, 107766, doi:10.1016/j.celrep.2020.107766 (2020).

16. Goodman, A. L. et al. A signaling network reciprocally regulates genes associated with acute infection and chronic persistence in Pseudomonas aeruginosa. Dev Cell 7, 745–754 (2004).

17. Lapouge, K., Schubert, M., Allain, F. H. & Haas, D. Gac/Rsm signal transduction pathway of gamma-proteobacteria: from RNA recognition to regulation of social behaviour. Molecular microbiology 67, 241–253 (2008).

18. Hood, R. D. et al. A type VI secretion system of Pseudomonas aeruginosa targets a toxin to bacteria. Cell Host Microbe 7, 25–37 (2010).

19. Snider, J. & Houry, W. A. MoxR AAA+ ATPases: a novel family of molecular chaperones? J Struct Biol 156, 200–209, doi:10.1016/j.jsb.2006.02.009 (2006).

20. Tsai, Y. C. et al. Insights into the mechanism and regulation of the CbbQO-type Rubisco activase, a MoxR AAA+ ATPase. Proc Natl Acad Sci U S A 117, 381–387, doi:10.1073/pnas.1911123117 (2020).

21. Paludan, S. R., Pradeu, T., Masters, S. L. & Mogensen, T. H. Constitutive immune mechanisms: mediators of host defence and immune regulation. Nature reviews. Immunology 21, 137–150, doi:10.1038/s41577-020-0391-5 (2021).

22. Jorgensen, F. et al. RpoS-dependent stress tolerance in Pseudomonas aeruginosa. Microbiology (Reading) 145 **( Pt** **4**), 835–844, doi:10.1099/13500872-145-4-835 (1999).

23. Munguia, J. et al. The Mla pathway is critical for Pseudomonas aeruginosa resistance to outer membrane permeabilization and host innate immune clearance. Journal of molecular medicine 95, 1127–1136, doi:10.1007/s00109-017-1579-4 (2017).

24. Russell, A. B. et al. Diverse type VI secretion phospholipases are functionally plastic antibacterial effectors. Nature 496, 508–512, doi:10.1038/nature12074 (2013).

25. Salomon, D. et al. Marker for type VI secretion system effectors. Proc Natl Acad Sci U S A 111, 9271–9276, doi:10.1073/pnas.1406110111 (2014).

26. Russell, A. B. et al. A widespread bacterial type VI secretion effector superfamily identified using a heuristic approach. Cell Host Microbe 11, 538–549, doi:10.1016/j.chom.2012.04.007 (2012).

27. Gebhardt, M. J., Kambara, T. K., Ramsey, K. M. & Dove, S. L. Widespread targeting of nascent transcripts by RsmA in Pseudomonas aeruginosa. Proc Natl Acad Sci U S A 117, 10520–10529, doi:10.1073/pnas.1917587117 (2020).

28. Wilhelm, S., Tommassen, J. & Jaeger, K. E. A novel lipolytic enzyme located in the outer membrane of Pseudomonas aeruginosa. J Bacteriol 181, 6977–6986, doi:10.1128/JB.181.22.6977-6986.1999 (1999).

29. Filkin, S. Y., Lipkin, A. V. & Fedorov, A. N. Phospholipase Superfamily: Structure, Functions, and Biotechnological Applications. Biochemistry (Mosc) 85, S177–S195, doi:10.1134/S0006297920140096 (2020).

30. Hernandez, R. E., Gallegos-Monterrosa, R. & Coulthurst, S. J. Type VI secretion system effector proteins: Effective weapons for bacterial competitiveness. Cell Microbiol 22, e13241, doi:10.1111/cmi.13241 (2020).

31. Fridman, C. M., Keppel, K., Gerlic, M., Bosis, E. & Salomon, D. A comparative genomics methodology reveals a widespread family of membrane-disrupting T6SS effectors. Nat Commun 11, 1085, doi:10.1038/s41467-020-14951-4 (2020).

32. Zheng, L., Lin, Y., Lu, S., Zhang, J. & Bogdanov, M. Biogenesis, transport and remodeling of lysophospholipids in Gram-negative bacteria. Biochim Biophys Acta Mol Cell Biol Lipids 1862, 1404–1413, doi:10.1016/j.bbalip.2016.11.015 (2017).

33. Lin, Y. et al. The phospholipid-repair system LplT/Aas in Gram-negative bacteria protects the bacterial membrane envelope from host phospholipase A2 attack. J Biol Chem 293, 3386–3398, doi:10.1074/jbc.RA117.001231 (2018).

34. Morehouse, B. R. et al. STING cyclic dinucleotide sensing originated in bacteria. Nature 586, 429–433, doi:10.1038/s41586-020-2719-5 (2020).

35. Flores-Diaz, M., Monturiol-Gross, L., Naylor, C., Alape-Giron, A. & Flieger, A. Bacterial Sphingomyelinases and Phospholipases as Virulence Factors. Microbiol Mol Biol Rev 80, 597–628, doi:10.1128/MMBR.00082-15 (2016).

36. Stover, C. K. et al. Complete genome sequence of Pseudomonas aeruginosa PA01, an opportunistic pathogen. Nature 406, 959–964 (2000).

37. Yu, Y. et al. Genomic patterns of pathogen evolution revealed by comparison of Burkholderia pseudomallei, the causative agent of melioidosis, to avirulent Burkholderia thailandensis. BMC microbiology 6, 46 (2006).

38. Paulsen, I. T. et al. Complete genome sequence of the plant commensal Pseudomonas fluorescens Pf-5. Nat Biotechnol 23, 873–878, doi:nbt1110 [pii] 10.1038/nbt1110 (2005).

39. Rietsch, A., Vallet-Gely, I., Dove, S. L. & Mekalanos, J. J. ExsE, a secreted regulator of type III secretion genes in Pseudomonas aeruginosa. Proc Natl Acad Sci U S A 102, 8006–8011 (2005).

40. Chandler, J. R. et al. Mutational analysis of Burkholderia thailandensis quorum sensing and self-aggregation. J Bacteriol 191, 5901–5909 (2009).

41. Gibson, D. G. et al. Enzymatic assembly of DNA molecules up to several hundred kilobases. Nature methods 6, 343–345, doi:10.1038/nmeth.1318 (2009).

42. Hoang, T. T., Kutchma, A. J., Becher, A. & Schweizer, H. P. Integration-proficient plasmids for Pseudomonas aeruginosa: site-specific integration and use for engineering of reporter and expression strains. Plasmid 43, 59–72 (2000).

43. Leroux, M. et al. Quantitative single-cell characterization of bacterial interactions reveals type VI secretion is a double-edged sword. Proc Natl Acad Sci U S A 109, 19804–19809, doi:10.1073/pnas.1213963109 (2012).

44. Choi, K. H. et al. Genetic tools for select-agent-compliant manipulation of Burkholderia pseudomallei. Applied and environmental microbiology 74, 1064–1075 (2008).

45. Bligh, E. G. & Dyer, W. J. A rapid method of total lipid extraction and purification. Can J Biochem Physiol 37, 911–917, doi:10.1139/o59-099 (1959).

46. Shiva, S. et al. Lipidomic analysis of plant membrane lipids by direct infusion tandem mass spectrometry. *Methods in molecular biology (Clifton*, N.J 1009, 79–91, doi:10.1007/978-1-62703-401-2_9 (2013).

47. Lee, K. M. et al. A Genetic Screen Reveals Novel Targets to Render Pseudomonas aeruginosa Sensitive to Lysozyme and Cell Wall-Targeting Antibiotics. Front Cell Infect Microbiol 7, 59, doi:10.3389/fcimb.2017.00059 (2017).

48. Kulasekara, H. D. et al. A novel two-component system controls the expression of Pseudomonas aeruginosa fimbrial cup genes. Molecular microbiology 55, 368–380, doi:10.1111/j.1365-2958.2004.04402.x (2005).

49. Gallagher, L. A. Methods for Tn-Seq Analysis in Acinetobacter baumannii. *Methods in molecular biology (Clifton*, N.J 1946, 115–134, doi:10.1007/978-1-4939-9118-1_12 (2019).

50. Chen, I. A. et al. The IMG/M data management and analysis system v.6.0: new tools and advanced capabilities. Nucleic Acids Res 49, D751-D763, doi:10.1093/nar/gkaa939 (2021).

51. Dobson, L., Remenyi, I. & Tusnady, G. E. CCTOP: a Consensus Constrained TOPology prediction web server. Nucleic Acids Res 43, W408–412, doi:10.1093/nar/gkv451 (2015).

52. Lopalco, P., Stahl, J., Annese, C., Averhoff, B. & Corcelli, A. Identification of unique cardiolipin and monolysocardiolipin species in Acinetobacter baumannii. Sci Rep 7, 2972, doi:10.1038/s41598-017-03214-w (2017).

53. Harvat, E. M. et al. Lysophospholipid flipping across the Escherichia coli inner membrane catalyzed by a transporter (LplT) belonging to the major facilitator superfamily. J Biol Chem 280, 12028–12034, doi:10.1074/jbc.M414368200 (2005).

54. Lin, Y., Zheng, L. & Bogdanov, M. Measurement of Lysophospholipid Transport Across the Membrane Using Escherichia coli Spheroplasts. *Methods in molecular biology (Clifton*, N.J 1949, 165–180, doi:10.1007/978-1-4939-9136-5_13 (2019).

55. Silverman, J. M. et al. Separate inputs modulate phosphorylation-dependent and - independent type VI secretion activation. Molecular microbiology 82, 1277–1290, doi:10.1111/j.1365-2958.2011.07889.x (2011).

56. Mougous, J. D. et al. A virulence locus of Pseudomonas aeruginosa encodes a protein secretion apparatus. Science 312, 1526–1530 (2006).

